# Genie: An interactive real-time simulation for teaching genetic drift

**DOI:** 10.1101/268672

**Authors:** Andreina I. Castillo, Ben H. Roos, Michael S. Rosenberg, Reed A. Cartwright, Melissa A. Wilson

## Abstract

Neutral evolution is a fundamental concept in evolutionary biology but teaching this and other non-adaptive concepts is specially challenging. Here we present Genie, a browser-based educational tool that facilitates demonstration of concepts such as genetic drift, population isolation, gene flow, and genetic mutation. Because it does not need to be downloaded and installed, Genie can scale to large groups of students and is useful for both in-person and online instruction. Genie was used to teach genetic drift to Evolution students at Arizona State University during Spring 2016 and Spring 2017. The effectiveness of Genie to teach key genetic drift concepts and misconceptions was assessed with the Genetic Drift Inventory developed by Price et al. (2014). Overall, Genie performed comparably to that of traditional static methods across all evaluated classes. We have empirically demonstrated that Genie can be successfully integrated with traditional instruction to reduce misconceptions about genetic drift.

## 1. Introduction

A well-recognized challenge in biological science education is successfully teaching evolutionary concepts (Alters and Nelson 2002). However, even within the same discipline, some topics remain more challenging to teach than others, and the number and efficacy of tools available for instruction varies (Shulman 1987; Ziadie and Andrews 2018). For instance, multiple strategies have been developed to improve the teaching of concepts like natural selection (Ziadie and Andrews 2018). On the other hand, best practices to teach equally important topics such as non-adaptive evolution remain largely understudied (Kalinowski et al. 2013). This is particularly problematic for topics like genetic drift because concepts of adaptive and non-adaptive evolution form independent elements in evolutionary thinking, and better understanding of one does not necessarily implies better comprehension of the other (Beggrow and Nehm 2012). To address this, studies devoted to developing, improving, and testing teaching strategies for non-adaptive evolutionary concepts are needed.

Previous studies have created approaches aimed to identify student misconceptions regarding genetic drift (Andrews et al. 2012; Price et al. 2014), and study activities and software have been developed, tested, and made publicly available (Price et al. 2016; Revell 2019; Staub 2002). These serve as indicators that the knowledge gap regarding genetic drift instruction is being addressed. Nonetheless, diverse class environments, student cohorts, and even teaching styles require distinct sets of tools; therefore, new tools are important for improving evolutionary instruction. Furthermore, there is an academic push for improving the teaching strategies currently set in place and to utilize alternative instruction methods (Lee et al. 2017; Nelson 2008; Tanner and Allen 2005). In particular, teaching strategies that favor discussion and testing of evolutionary concepts among student have been shown to be most effective (George M. Slavich and Zimbardo 2012). As a result, tools that can be used to facilitate free in-class exploration of evolutionary concepts, are especially useful since they allow students to both learn these concepts and develop critical thinking skills.

Here, we developed a web application (Genie) designed to demonstrate several population genetics and evolutionary notions including genetic drift, gene flow, and random mutation. This application conducts a real time simulation of the change in allele frequencies in a finite population of spatially isolated individuals. Using colors, the application allows students to visualize changes in a population over time and understand how those visual changes translate to fluctuations in allele frequency, and eventually, fixation or loss of an allele. This web-based software is accessible to students and leads to increased knowledge of genetic drift concepts, as tested using a Genetic Drift Inventory (Price et al. 2014). These types of assessments have proven to be useful in capturing student’s understanding of other complex evolutionary concepts in the past (Perez et al. 2013). The Genie software requires no startup other than navigating to a web page, thus making the use of programmed stochastic simulations to demonstrate the concept of genetic drift practical to both educators and students in face-to-face and online instruction.

## 2. Methods

### 2.1 Genie simulation program

Genie (https://cartwrig.ht/apps/genie/) is a web-based, stochastic simulation app written in JavaScript. The simulation uses a spatially explicit Moran Model (Nei et al. 1976) to describe a finite population of 1,024 individuals on a 32 by 32 grid. Individuals are haploid with a single locus. The locus mutates according to the infinite alleles model (Nei et al. 1976). Genie works as follows:

- *Population Initialization*. The simulation begins when a population is randomly initialized according to Hoppe’s Urn (Perez et al. 2013). Briefly, the population is created one individual at a time, and each individual either carries a new, unique allele or is a copy of a previously created individual. The probability that individual *i* + 1 has a new allele is *θ/(θ+i)* and the probability that the individual copies an existing allele is *(i)/(θ+i)*, where *θ = 2Nμ, N* is the population size, and *μ* is the mutation rate. If an individual copies an allele, it randomly chooses a previously initialized individual uniformly. At initialization *μ* is *= 0.001* to ensure diversity within the initial population, but the mutation rate of each generation can be specified by the user, defaulting to 0.
- *Algorithm*. At each step of the simulation, a randomly selected individual dies, leaving its corresponding cell momentarily empty. A parent allele is then randomly selected from the eight immediate neighboring cells (including adjacent and diagonal). Cells on the edges and corners of the simulation have fewer neighbors than internal cells, causing a small edge effect. The probability that a new individual will have the same allele as its parent its 1-*μ*, and the probability that an individual has a new, unique allele is *μ*. Each ‘generation’ consists of 2000 death/birth steps after which the population is redrawn in the visualization window.
- *Running*. The application contains four components: a grid, where the population is displayed (Supplementary file 1a); a control panel, where users can manipulate the simulation’s mutation parameter (Supplementary file 1b); an upper graph, where users can see the number of alleles in the population at any given time (Supplementary file 1c); and a lower graph, where users can see the frequency of different alleles at any given time (Supplementary file 1d). Both graphs update in real time as the simulation runs. Each initial allele is assigned one of 18 basic colors, while each mutant allele is assigned one of six neon colors. A single button allows users to toggle between starting the simulation or pausing it. A reset button allows users to restart and reinitialize the simulation at any point.
- *Barriers*. Users can create a barrier in the population grid. To do so, users alter a cell (by clicking on it) or alter a set of cells (by clicking and dragging the cursor to select multiple cells). When a barrier is created, the color associated with the cell changes to black. Barriers act neither as parent cells (they are never replicated) nor die. Thus, for each created barrier cell the total population size declines by one. By building barriers, users can construct physical constraints that restrict the movement of alleles between subpopulations. Barriers can be used to create subpopulations of different sizes and shapes, as well as to study the effects of corridors on gene flow. Barriers can be removed by clicking on the chosen cell(s) a second time; this will set the cell color to white and designate the cell as unoccupied. Neighboring cells will replicate into unoccupied cells; unoccupied cells cannot serve as a parent of a neighboring cell.
- *Forced Mutation*. Users can force a mutation to occur in a manner similar to creating barriers. Cells can be mutated by holding the SHIFT button while clicking the cell, or while clicking and dragging the cursor across several cells. Forcing a mutation immediately creates a new, unique allele in each of the chosen cell(s).

### 2.2. Data collection

Genie’s efficacy as a tool for teaching Genetic Drift concepts was tested in the Evolution (BIO345) class at Arizona State University (ASU). Genie was used during the practical portion (recitation) of the BIO345 course in the Spring 2016 and Spring 2017 classes. All participants in the Spring 2016 class used Genie during practical class sessions. In the Spring 2017 class, half of the participants used the dynamic visualization of Genie while the other half used static illustrations. Participants in both the Spring 2016 and Spring 2017 classes were given the option to opt-in to the study at the end of the semester. In addition, participants were given the option to provide their demographic information: reported gender, reported ethnicity, and first-generation college student status. All research was reviewed and approved by Arizona State University’s IRB protocol STUDY00003707.

The impact of Genie as a tool for teaching concepts of genetic drift was evaluated using the Genetic Drift Inventory (Price et al. 2014). The inventory was used without changes (22 questions assessing different aspects of genetic drift) in pre- and post-recitation assessments. The pre- and post-recitation assessments (considered as homework for the entire class) were individually answered by each participant. The pre-recitation assessment was posted online on Blackboard two days before recitation. Participants were asked to answer all questions by 3:00 pm of the day of the recitation. The post-recitation assessment was posted on Blackboard at 9:00 pm after the last recitation session ended. Participants had two days to individually complete the post-recitation assessment. All participants were allowed the same amount of time to complete both the pre- and post-recitation assessments. Participant’s answers were recorded, and their individual pre- and post-recitation scores were calculated by summing the number of correctly answered questions (value 1 point) out of the 22 questions in the Genetic Drift Inventory.

### 2.3. Genie assessment

The complete dataset was divided into two major groups based on instruction year. These groups were: the entire Spring 2016 class (henceforth referred to as Genie 2016) and the entire Spring 2017 class. The 2017 class was further subdivided into groups based on the instruction method used during the practical class session. These groups were: participants that used Genie during the recitation session in 2017 (henceforth referred to as Genie 2017) and the participants who did not use Genie during the recitation session in 2017 (henceforth referred to as Non-Genie 2017). The Genie 2016 class was subsequently divided into eight in-class groups of roughly equal size, while each 2017 class was divided into four in-class groups of roughly equal size (two Genie and two Non-Genie). The groups were designated based on recitation start times, Graduate Teaching Assistants (TA) pairs; and in the case of 2017, on the use of dynamic (Genie) vs. static (Non-Genie) instruction methods. No more than 48 participants participated in each recitation session. All analyses and figures were developed using R v3.2. The code and datasets used are available (Supplementary files 2-12, https://github.com/AndreinaCastillo/Genie_manuscript_data_analysis).

The putative relationship between participants’ demographics and the pre- and post-recitation scores was evaluated using a two-way ANOVA. The following demographic parameters were used as explanatory variables: reported gender, reported ethnicity, and first-generation college student status. In the case of 2017, the use of Genie as an instruction tool was also considered as an explanatory variable. The two-way ANOVA was performed independently for Genie 2016, Genie 2017, and Non-Genie 2017. Next, we assessed if the pre- and post-recitation performance varied between the three class groups or among subgroups within each class. To conduct this analysis, the distribution of pre- and post-recitation scores was assessed using the ‘fitdistrplus’ (Delignette-Muller and Dutang 2015) and ‘betareg’ (Cribari-Neto and Zeileis 2010) R packages. Potential differences between pre- and post-recitation scores were evaluated both between classes and within each in-class group. In addition, a Cohen’s d was used to measure the effect size between pre- and post-recitation scores within each class, and to estimate differences in pre- and post-recitation scores between Genie 2017 and Non-Genie 2017.

In addition, potential differences in groups of participants based on their initial performance levels following instruction were assessed. Participant’s scores within Genie 2016, Genie 2017, and Non-Genie 2017 were divided into four quantiles based on their pre-recitation scores. The first quantile included participants with scores ranging from 0 to 0.25, the second quantile included participants with scores between 0.26-0.5, the third quantile included participants with scores of 0.51-0.75, and the fourth quantile included participants with scores of 0.76-1. For each quantile within each class, a paired Student’s t-test was performed in order to evaluate if participants with different performance levels (i.e. within each quantile) benefited differently from the use of Genie. Furthermore, a paired Student’s t-test was performed between individual participants’ pre- and post-recitation scores within each class.

Finally, question-specific performance was evaluated to determine how Genie aided participants in addressing the specific genetic drift concepts and misconceptions listed in the Genetic Drift Inventory (Price et al. 2014). The number of correct answers in pre- and post-recitation sessions associated with each question were calculated from participants’ individual answers, and the totals where then compiled by class. Differences between pre- and post-recitation scores for each question were assessed using a McNemars χ^2^ test. In addition, the difference in the number of correct answers per question in Genie 2017 vs. Non-Genie pre- and post-recitation sessions was assessed using a Fisher’s exact test.

## 3. Results

Demographic representation varied among cohorts (Table 1). Participants identifying as People of Color (‘POC’) (N = 168) were less represented compared to participants identifying as ‘White’ (N = 238) in Genie 2016. Both groups were roughly equal in Genie 2017 (‘POC’ = 136 and ‘White’ = 144) and Non-Genie 2017 (‘POC’ = 120 and ‘White’ = 112). On the other hand, participants identifying as ‘Female’ (N = 230 in Genie 2016, N = 190 in Genie 2017, and N = 140 in Non-Genie 2017) were more numerous than participants identifying as ‘Male’ (N = 176 in Genie 2016, N = 90 in Genie 2017, and N = 92 in Non-Genie 2017). Likewise, ‘First-generation’ college students were less numerous (N = 126 in Genie 2016, N = 46 in Genie 2017, and N = 36 in Non-Genie 2017) than ‘Not First-generation’ college students (N = 280 in Genie 2016, N = 234 in Genie 2017, and N = 196 in Non-Genie 2017). Regardless of these differences, pre- and post-recitation performance levels were similar in participants from different demographic backgrounds across the three evaluated class groups (Figure 1).

**Table 1.** Demographic breakdowns of participants in each year and section shows variable representation of different groups. The breakdown of participants in each year of the class who participated in the assessment, including those who self-identified as people of color (‘POC’) or ‘white’, ‘female’ or ‘male’, and ‘first-generation’ college students or not.

| Categories | Genie 2016 | Genie 2017 | Non-Genie 2017 |  |
| --- | --- | --- | --- | --- |
| POC | 168 | 136 | 120 | 403 |
| White | 238 | 144 | 112 |  |
| Female | 230 | 190 | 140 | 404 |
| Male | 176 | 90 | 92 |  |
| Not first-generation college | 280 | 234 | 196 |  |
| First-generation college | 126 | 46 | 36 | 405 |

**Figure 1.**
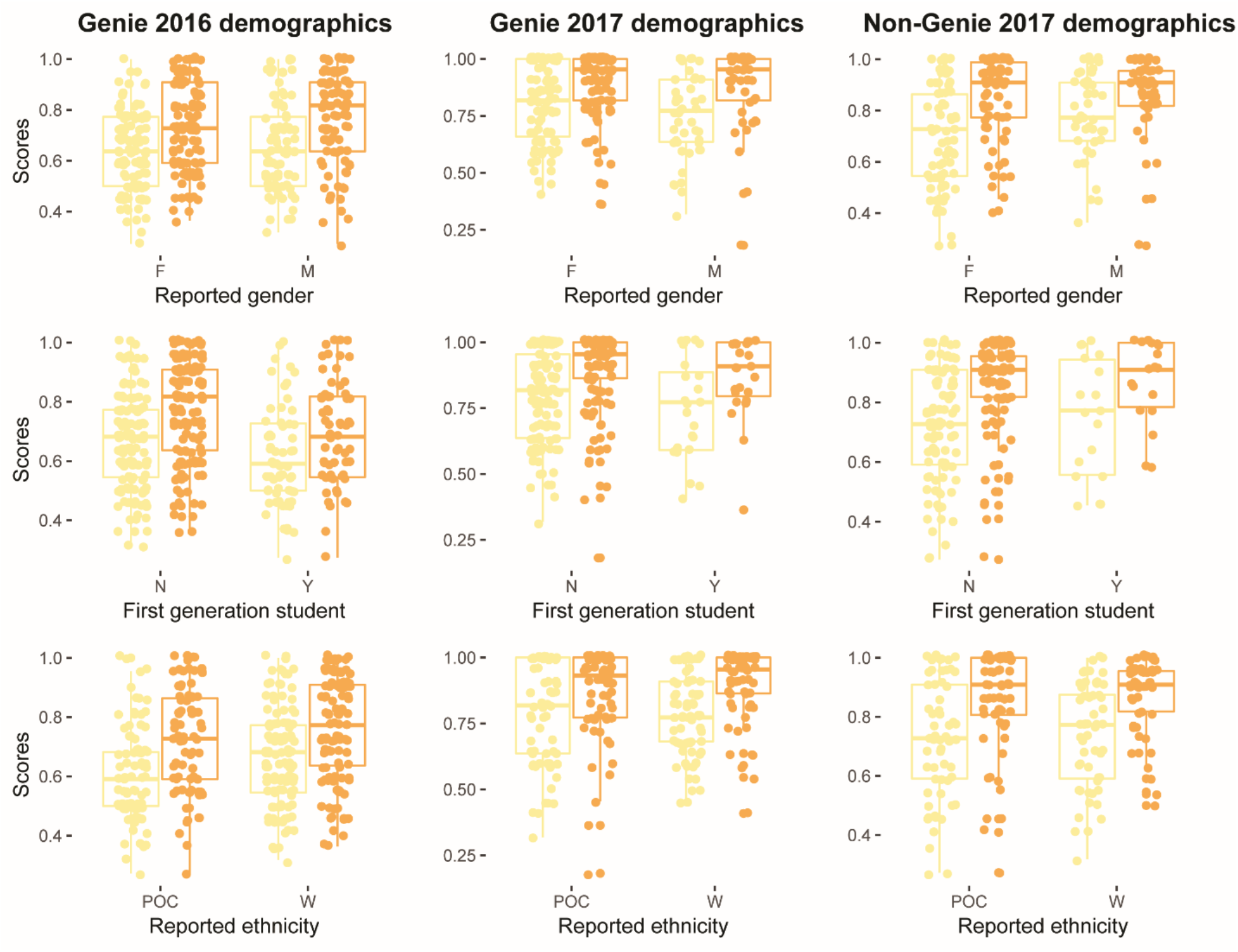
Pre- and post-recitation performance was comparable in participants from different demographic groups. Pre-recitation (pale yellow) and post-recitation (dark yellow) scores are plotted. Three demographic variables are plotted: ‘Reported gender’, ‘First-generation’ college, and ‘Reported ethnicity’.

A two-way ANOVA found that most demographic explanatory variables did not affect pre- and post-recitation scores (Table 2). For instance, the variable ‘Reported gender’ did not have a clear effect on pre- or post-recitation scores in either year (Genie 2016 pre-recitation F = 1.004, p-value = 0.318; Genie 2016 post-recitation F = 2.388, p-value = 0.124; 2017 pre-recitation F = 0.037, p-value = 0.848; 2017 post-recitation F = 0.010, p-value = 0.921). Likewise, the variable ‘Reported ethnicity’ did not have a clear effect in pre- (F = 1.046, p-value = 0.397) or post-recitation scores (F = 0.372, p-value = 0.896) in Genie 2016, or the post-recitation scores (F = 0.895, p-value = 0.511) in 2017. However, it did have a significant effect (F = 2.286, p-value = 0.029) in pre-recitation scores in 2017.

**Table 2.**
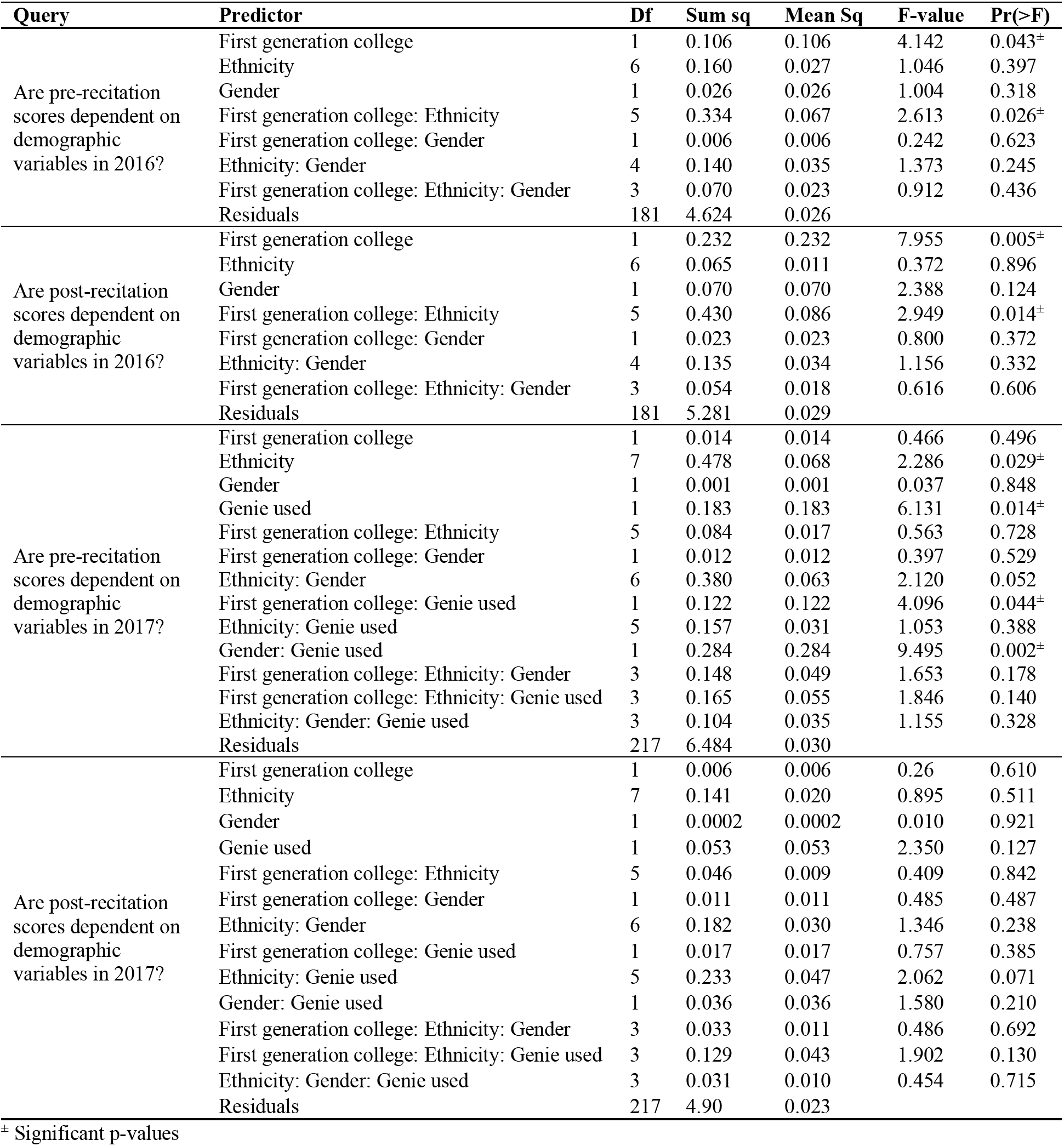
Most demographic predictors did not affect performance on the evaluation of genetic drift knowledge. The general linear regression of pre- and post-recitation scores with demographic predictors, the degrees of freedom (Df), and the summary statistics.

| Query | Predictor | Df | Sum sq | Mean Sq | F-value | Pr(>F) |
| --- | --- | --- | --- | --- | --- | --- |
| Are pre-recitation scores dependent on demographic variables in 2016? | First generation college | 1 | 0.106 | 0.106 | 4.142 | 0.043<sup>±</sup> |
|  | Ethnicity | 6 | 0.160 | 0.027 | 1.046 | 0.397 |
|  | Gender | 1 | 0.026 | 0.026 | 1.004 | 0.318 |
|  | First generation college: Ethnicity | 5 | 0.334 | 0.067 | 2.613 | 0.026<sup>±</sup> |
|  | First generation college: Gender | 1 | 0.006 | 0.006 | 0.242 | 0.623 |
|  | Ethnicity: Gender | 4 | 0.140 | 0.035 | 1.373 | 0.245 |
|  | First generation college: Ethnicity: Gender | 3 | 0.070 | 0.023 | 0.912 | 0.436 |
|  | Residuals | 181 | 4.624 | 0.026 |  |  |
| Are post-recitation scores dependent on demographic variables in 2016? | First generation college | 1 | 0.232 | 0.232 | 7.955 | 0.005<sup>±</sup> |
|  | Ethnicity | 6 | 0.065 | 0.011 | 0.372 | 0.896 |
|  | Gender | 1 | 0.070 | 0.070 | 2.388 | 0.124 |
|  | First generation college: Ethnicity | 5 | 0.430 | 0.086 | 2.949 | 0.014<sup>±</sup> |
|  | First generation college: Gender | 1 | 0.023 | 0.023 | 0.800 | 0.372 |
|  | Ethnicity: Gender | 4 | 0.135 | 0.034 | 1.156 | 0.332 |
|  | First generation college: Ethnicity: Gender | 3 | 0.054 | 0.018 | 0.616 | 0.606 |
|  | Residuals | 181 | 5.281 | 0.029 |  |  |
| Are pre-recitation scores dependent on demographic variables in 2017? | First generation college | 1 | 0.014 | 0.014 | 0.466 | 0.496 |
|  | Ethnicity | 7 | 0.478 | 0.068 | 2.286 | 0.029<sup>±</sup> |
|  | Gender | 1 | 0.001 | 0.001 | 0.037 | 0.848 |
|  | Genie used | 1 | 0.183 | 0.183 | 6.131 | 0.014<sup>±</sup> |
|  | First generation college: Ethnicity | 5 | 0.084 | 0.017 | 0.563 | 0.728 |
|  | First generation college: Gender | 1 | 0.012 | 0.012 | 0.397 | 0.529 |
|  | Ethnicity: Gender | 6 | 0.380 | 0.063 | 2.120 | 0.052 |
|  | First generation college: Genie used | 1 | 0.122 | 0.122 | 4.096 | 0.044<sup>±</sup> |
|  | Ethnicity: Genie used | 5 | 0.157 | 0.031 | 1.053 | 0.388 |
|  | Gender: Genie used | 1 | 0.284 | 0.284 | 9.495 | 0.002<sup>±</sup> |
|  | First generation college: Ethnicity: Gender | 3 | 0.148 | 0.049 | 1.653 | 0.178 |
|  | First generation college: Ethnicity: Genie used | 3 | 0.165 | 0.055 | 1.846 | 0.140 |
|  | Ethnicity: Gender: Genie used | 3 | 0.104 | 0.035 | 1.155 | 0.328 |
|  | Residuals | 217 | 6.484 | 0.030 |  |  |
| Are post-recitation scores dependent on demographic variables in 2017? | First generation college | 1 | 0.006 | 0.006 | 0.26 | 0.610 |
|  | Ethnicity | 7 | 0.141 | 0.020 | 0.895 | 0.511 |
|  | Gender | 1 | 0.0002 | 0.0002 | 0.010 | 0.921 |
|  | Genie used | 1 | 0.053 | 0.053 | 2.350 | 0.127 |
|  | First generation college: Ethnicity | 5 | 0.046 | 0.009 | 0.409 | 0.842 |
|  | First generation college: Gender | 1 | 0.011 | 0.011 | 0.485 | 0.487 |
|  | Ethnicity: Gender | 6 | 0.182 | 0.030 | 1.346 | 0.238 |
|  | First generation college: Genie used | 1 | 0.017 | 0.017 | 0.757 | 0.385 |
|  | Ethnicity: Genie used | 5 | 0.233 | 0.047 | 2.062 | 0.071 |
|  | Gender: Genie used | 1 | 0.036 | 0.036 | 1.580 | 0.210 |
|  | First generation college: Ethnicity: Gender | 3 | 0.033 | 0.011 | 0.486 | 0.692 |
|  | First generation college: Ethnicity: Genie used | 3 | 0.129 | 0.043 | 1.902 | 0.130 |
|  | Ethnicity: Gender: Genie used | 3 | 0.031 | 0.010 | 0.454 | 0.715 |
|  | Residuals | 217 | 4.90 | 0.023 |  |  |

<sup>±</sup> Significant p-values

The variable ‘First-generation’ college had a statistical effect in pre- (F = 4.142, p-value = 0.043) and post-recitation scores (F = 7.955, p-value = 0.005) in Genie 2016, but not in 2017 (pre-recitation F = 0.466, p-value = 0.496; post-recitation F = 0.260, p-value = 0.610). ‘First-generation’ college students had lower pre-recitations scores than ‘Not First-generation’ students in Genie 2016 and Genie 2017, while the opposite trend was observed in Non-Genie 2017 (Figure 1). While post-recitation scores were higher than pre-recitation scores in both groups, ‘First-generation’ college students showed slightly less improvement than ‘Not First-generation’ college students. For instance, ‘First-generation’ college students saw an increase of 0.09 compared to 0.13 for ‘Not First-generation’ college students in Genie 2016. The same trend was observed in Non-Genie 2017 (0.13 vs. 0.18 for ‘First-generation’ college students and ‘Not First-generation’ college students, respectively) and Genie 2017 (0.13 vs. 0.14 for ‘First-generation’ college students and ‘Not First-generation’ college students, respectively). It should be noted that the number of ‘First-generation’ college students was higher in Genie 2016 (N = 126) as well as Genie 2017 (N = 46), compared to Non-Genie 2017 (N = 36). The ‘Genie used’ variable showed a significant effect (F = 6.131, p-value = 0.014) in pre-recitation scores in 2017 but not in post-recitation scores (F = 2.350, p-value = 0.127). The difference between mean pre-recitation scores between the Genie and Non-Genie 2017 classes was small (0.0495, or approximately 1 out of 22 questions), with Genie 2017 having a higher mean pre-recitation score (0.7808) than Non-Genie 2017 (0.7313). The difference in post-recitation scores was 0.0368, with Genie 2017 still having a higher mean (0.8828) than Non-Genie 2017 (0.8460).

Overall, pre- and post-recitation scores were different among the classes analyzed. The mean pre-recitation score for Genie 2016 (0.6444) was lower than in either Genie 2017 or Non-Genie 2017 (see above). This same trend was maintained for post-recitation scores in Genie 2016 (0.7481) compared to Genie 2017 or Non-Genie 2017 (see above). Differences in post-recitation scores could be largely explained by the initial differences in pre-recitations scores, or in other words, the initial class performance (Table 3). Furthermore, in-class groups showed similar performance levels among Genie 2016, Genie 2017, and Non-Genie 2017. An outlier to this observation was the ‘TA Pair1 7:30pm’ in-class group (p-value=0.017) during Genie 2016, which was composed exclusively of honor students. Overall, the density curve of post-recitation scores showed that participant performance improved in all classes regardless of the instruction method used (Figure 2). Furthermore, Cohen’s d values (Table 4) showed a moderate improvement in post-recitation scores compared to the pre-recitation scores in Genie 2016 (0.608, CI: 0.408-0.807), Genie 2017 (0.632, CI: 0.410-0.855), and Non-Genie 2017 (0.658, CI: 0.430-0.886). In addition, Cohen’s d values also showed a small difference in pre- (0.272, CI: 0.051-0.493) and post-recitation (0.242, CI: 0.021-0.462) scores between Genie 2017 and Non-Genie 2017.

**Figure 2.**
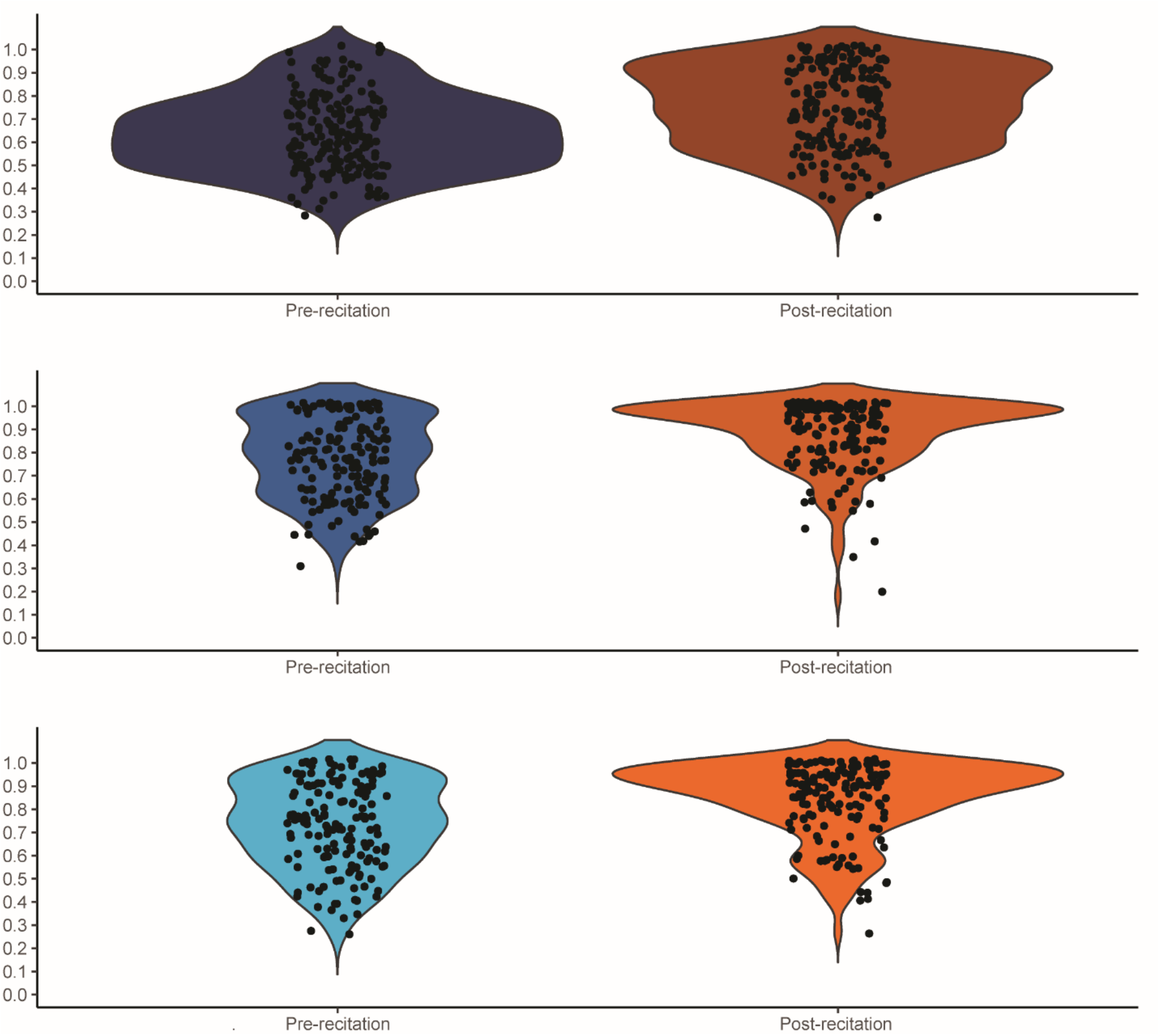
Post-recitation performance improved regardless of the instruction method. Pre-Recitation (blue) and post-recitation (orange) scores are plotted for each class.

**Table 3.** Post-recitation scores were mostly influenced by pre-recitation scores, but not class section. The Beta regression tests of pre- and post-recitation scores for specific queries, including the predictors, estimates, standard errors (SE), z-scores, and p-values.

| Query | Predictor | Estimate | SE | z-score | p-value |
| --- | --- | --- | --- | --- | --- |
| Are the pre-recitation scores different across classes (Genie 2016, Genie 2017, and Non-Genie 2017)? | Genie 2017 | 0.961 | 0.104 | 9.241 | $< 2 \times 10^{-16} \pm$ |
| | Non-Genie 2017 | 0.521 | 0.102 | 5.081 | $3.75 \times 10^{-07} \pm$ |
| Are the post-recitation scores dependent on the pre-recitation scores and/or class (Genie 2016, Genie 2017, and Non-Genie 2017)? | Pre-recitation scores | 3.453 | 0.237 | 14.55 | $< 2 \times 10^{-16} \pm$ |
| | Genie 2017 | 0.509 | 0.103 | 4.948 | $7.51 \times 10^{-07} \pm$ |
| | Non-Genie 2017 | 0.281 | 0.099 | 2.847 | $0.004 \pm$ |
| Are the post-recitation scores of 2016 dependent on the pre-recitation scores and/or the class section? | Pre-recitation score | 3.198 | 0.391 | 8.17 | $3.08 \times 10^{-16} \pm$ |
|  | TA Pair1 4:30pm | -0.384 | 0.217 | -1.764 | 0.078 |
|  | TA Pair1 6:00pm | -0.164 | 0.233 | -0.702 | 0.483 |
| | TA Pair1 7:30pm | -0.68 | 0.285 | -2.387 | $0.017 \pm$ |
|  | TA Pair2 3:00pm | 0.309 | 0.226 | 1.366 | 0.172 |
|  | TA Pair2 4:30pm | -0.096 | 0.205 | -0.468 | 0.64 |
|  | TA Pair2 6:00pm | -0.336 | 0.234 | -1.433 | 0.152 |
|  | TA Pair2 7:30pm | 0.052 | 0.242 | 0.213 | 0.831 |
| Are the post-recitation scores of 2017 dependent on the pre-recitation scores and/or the class section when using Genie? | Pre-recitation score | 3.52 | 0.452 | 7.797 | $6.64 \times 10^{-15} \pm$ |
|  | TA Pair1 7:00pm | -0.209 | 0.21 | -0.996 | 0.319 |
|  | TA Pair2 1:30pm | -0.032 | 0.209 | -0.152 | 0.879 |
|  | TA Pair2 7:00pm | 0.006 | 0.223 | 0.026 | 0.98 |
| Are the post-recitation scores of 2017 dependent on the pre-recitation scores and/or the class section when not using Genie? | Pre-recitation score | 3.583 | 0.37 | 9.695 | $< 2 \times 10^{-16} \pm$ |
|  | TA Pair1 4:30pm | -0.023 | 0.194 | -0.116 | 0.907 |
|  | TA Pair2 3:00pm | 0.33 | 0.188 | 1.75 | 0.08 |
|  | TA Pair2 4:30pm | -0.126 | 0.19 | -0.659 | 0.509 |

± Significant p-values

**Table 4.** Effect of Genie on learning outcomes. The size-effect analysis of recitation scores per year.

| Query | Cohen's d (Lower 95% CI, Upper 95% CI) |
| --- | --- |
| Are the scores different before and after instruction in 2016 (Genie)? | 0.608 (0.408, 0.807) |
| Are the scores different before and after instruction in 2017 (Genie)? | 0.632 (0.410, 0.855) |
| Are the scores different before and after instruction in 2017 (No-Genie)? | 0.658 (0.430, 0.886) |
| Are the pre-recitation scores different between 2017's Genie and Non-Genie classes? | 0.272 (0.051, 0.493) |
| Are the post-recitation scores different between 2017's Genie and Non-Genie classes? | 0.242 (0.021, 0.462) |
| Is the difference in pre- and post-recitation scores different between 2017's Genie and Non-Genie classes? | -0.087 (-0.307, 0.133) |

Post-recitation scores where higher than pre-recitation scores for most individual participants within all classes (Table 5, Supplementary file 13). Furthermore, participants across three out of four quantiles of initial pre-recitation scores showed improvement in their post-recitation scores (Table 6). There were too few participants (1-2 individuals across all groups) with pre-recitation scores between 0-0.25 (the lowest quantile), making unfeasible to statistically evaluate this group. In Genie 2016, the highest improvement was observed in participants with pre-recitation scores between 0.26-50 (T = −7.855, p-value = 4.294×10^−11^). In Genie 2017 (T = −7.118, p-value =1.191×10^−09^) and Non-Genie 2017 (T = −7.714, p-value = 4.412×10^−11^), the highest improvement was seen in participants with pre-recitation scores between 0.51-0.75.

**Table 5.** Individual participants in all classes showed higher post-recitation scores compared to their pre-recitation scores. Paired Student’s t-test for individual participants in pre- and post-recitation scores per class.

| Query | t -value (p-value) | Mean of the difference | Lower 95% CI, Upper 95% CI |
| --- | --- | --- | --- |
| Are paired post- and pre-recitation scores different in Genie 2016? | -9.747 (2.2x10<sup>-16</sup>)<sup>±</sup> | -0.104 | -0.125, -0.083 |
| Are paired post- and pre-recitation scores different in Genie 2017? | -8.966 (6.913x10<sup>-16</sup>)<sup>±</sup> | -0.102 | -0.124, -0.079 |
| Are paired post- and pre-recitation scores different in Non-Genie 2017? | -9.816 (2.2x10<sup>-16</sup>)<sup>±</sup> | -0.115 | -0.138, -0.092 |

<sup>±</sup> Significant p-values

**Table 6.** Participants from different pre-recitation scores quantiles showed improvement in their post-recitation scores. Student’s t-test for pre-to post-recitation scores depending on pre-recitation performance quantile.

| Query | Quantile (pre-recitation scores range) | t -value (p-value) | Lower 95% CI, Upper 95% CI | Mean pre-score | Mean post-score |
| --- | --- | --- | --- | --- | --- |
| Do different groups of participants have differences performances in Genie 2016? | 1<sup>st</sup> Quantile (0-0.25) | Not enough observations |  |  |  |
|  | 2<sup>nd</sup> Quantile (0.26-0.5) | -7.855 (4.294x10<sup>-11</sup>)<sup>±</sup> | -0.230, -0.137 | 0.448 | 0.631 |
|  | 3<sup>rd</sup> Quantile (0.51-0.75) | -4.897 (2.968x10<sup>-06</sup>)<sup>±</sup> | -0.120, -0.051 | 0.638 | 0.724 |
|  | 4<sup>th</sup> Quantile (0.76-1) | -3.630 (0.0004)<sup>±</sup> | -0.082, -0.024 | 0.859 | 0.913 |
| Do different groups of participants have differences performances in Genie 2017? | 1<sup>st</sup> Quantile (0-0.25) | Not enough observations |  |  |  |
|  | 2<sup>nd</sup> Quantile (0.26-0.5) | -5.186 (0.0014)<sup>±</sup> | -0.341, -0.142 | 0.448 | 0.689 |
|  | 3<sup>rd</sup> Quantile (0.51-0.75) | -7.118 (1.191x10<sup>-09</sup>)<sup>±</sup> | -0.222, -0.125 | 0.636 | 0.810 |
|  | 4<sup>th</sup> Quantile (0.76-1) | -3.835 (0.0002)<sup>±</sup> | -0.069, -0.022 | 0.903 | 0.948 |
| Do different groups of participants have differences performances in Non-Genie 2017? | 1<sup>st</sup> Quantile (0-0.25) | Not enough observations |  |  |  |
|  | 2<sup>nd</sup> Quantile (0.26-0.5) | -6.128 (1.203x10<sup>-06</sup>)<sup>±</sup> | -0.341, -0.170 | 0.421 | 0.676 |
|  | 3<sup>rd</sup> Quantile (0.51-0.75) | -7.714 (4.412x10<sup>-11</sup>)<sup>±</sup> | -0.214, -0.126 | 0.642 | 0.812 |
|  | 4<sup>th</sup> Quantile (0.76-1) | -2.337 (0.021)<sup>±</sup> | -0.059, -0.005 | 0.890 | 0.922 |

<sup>±</sup> Significant p-values

Participants from the top quantile (pre-recitation scores between 0.76-1), had the smallest improvement in their post-recitation scores, particularly in Non-Genie 2017 (Table 6).

Participants understanding of key genetic drift concepts and misconceptions statistically improved after instruction with or without Genie (Table 7, Figure 3). Post-recitation scores were generally higher in Genie 2017 than in Non-Genie 2017 except for two questions (Q10 and Q15, Figure 4). When the difference between pre- and post-recitation scores by question was plotted (Figure 5), there was variation in which questions had a higher score improvement in Non-Genie 2017 or Genie 2017. Non-Genie 2017 showed higher improvements in questions related to key concepts (Q1, Q10, and Q13) and misconceptions associated to genetic drift as natural selection/adaptation/acclimation to the environment that may result from a need to survive (M2), genetic drift as random mutation (Q9), and genetic drift as gene flow or migration (Q11). On the other hand, Genie 2017 showed higher improvements in misconceptions related to natural selection being always the most powerful mechanism of evolution, and as the primary agent of evolutionary change (M4). However, a Fisher’s exact test showed that instruction method (Genie vs. Non-Genie) was not associated with student’s switching answers from correct to incorrect or incorrect to correct between pre- and post-recitation (Table 8). Results were comparable with or without including students within honor sections (Supplementary file 14).

**Table 7.** For most individual questions, participant post-recitation performance improved across classes following instruction either with or without Genie. McNemar’s test performed on (in)correct to (in)correct pre- and post-recitation answers per question. McNemar statistic and p-value are provided. We show this for (a) Genie 2016, (b) Genie 2017, and (c) Non-Genie 2017.

| Evaluation | Question | Genie 2016 |  |  |  | McNemar stat. | p-value |
| --- | --- | --- | --- | --- | --- | --- | --- |
|  |  | Incorrect to incorrect switches | Incorrect to correct switches | Correct to incorrect switches | Correct to correct switches |  |  |
| Key concepts | Q1 | 8 | 22 | 14 | 159 | 1.778 | 0.182 |
|  | Q3 | 10 | 53 | 7 | 133 | 35.267 | 2.88x10<sup>-09±</sup> |
|  | Q15 | 9 | 26 | 15 | 153 | 2.951 | 0.086 |
|  | Q4 | 28 | 44 | 13 | 118 | 16.86 | 4.02x10<sup>-05±</sup> |
|  | Q10 | 21 | 28 | 18 | 136 | 2.174 | 0.14 |
|  | Q13 | 18 | 86 | 9 | 90 | 62.411 | 2.79x10<sup>-15±</sup> |
|  | Q16 | 58 | 28 | 43 | 74 | 3.169 | 0.075 |
| Misconceptions* | 1 Q7 | 10 | 32 | 19 | 142 | 3.314 | 0.069 |
|  | 2 Q5 | 38 | 51 | 15 | 99 | 19.636 | 9.37x10<sup>-06±</sup> |
|  | 2 Q6 | 69 | 53 | 19 | 62 | 16.056 | 6.15x10<sup>-05±</sup> |
|  | 2 Q8 | 71 | 46 | 22 | 64 | 8.471 | 0.004<sup>±</sup> |
|  | 3 Q2 | 39 | 41 | 27 | 96 | 2.882 | 0.09 |
|  | 4 Q9 | 8 | 28 | 21 | 146 | 1 | 0.317 |
|  | 4 Q12 | 36 | 60 | 22 | 85 | 17.610 | 2.71x10<sup>-05±</sup> |
|  | 4 Q17 | 49 | 58 | 12 | 84 | 30.229 | 3.84x10<sup>-08±</sup> |
|  | 4 Q20 | 22 | 40 | 18 | 123 | 8.345 | 0.004<sup>±</sup> |
|  | 5 Q14 | 61 | 46 | 32 | 64 | 2.513 | 0.113 |
|  | 5 Q19 | 28 | 36 | 39 | 100 | 0.12 | 0.729 |
|  | 5 Q22 | 45 | 30 | 30 | 98 | 0 | 1 |
|  | 6 Q11 | 17 | 50 | 17 | 119 | 16.254 | 5.54x10<sup>-05±</sup> |
|  | 6 Q18 | 12 | 29 | 8 | 154 | 11.919 | 5.56x10<sup>-04±</sup> |
|  | 6 Q21 | 20 | 24 | 28 | 131 | 0.308 | 0.579 |

<sup>±</sup> Significant p-values
\* Misconceptions (Price et al. 2014)

**Table 7. For most individual questions, participant post-recitation performance improved across classes following instruction either with or without Genie (cont.).**
| Evaluation | Question | Genie 2017 |  |  |  | McNemar stat. | p-value |
| --- | --- | --- | --- | --- | --- | --- | --- |
|  |  | Incorrect to incorrect switches | Incorrect to correct switches | Correct to incorrect switches | Correct to correct switches |  |  |
| Key concepts | Q1 | 0 | 6 | 3 | 155 | 1 | 0.317 |
|  | Q3 | 1 | 13 | 3 | 147 | 6.25 | 0.012<sup>±</sup> |
|  | Q15 | 5 | 17 | 8 | 134 | 3.24 | 0.072 |
|  | Q4 | 7 | 17 | 12 | 128 | 0.862 | 0.353 |
|  | Q10 | 9 | 22 | 6 | 127 | 9.143 | 0.002<sup>±</sup> |
|  | Q13 | 5 | 32 | 5 | 122 | 19.703 | 9.05x10<sup>-06±</sup> |
|  | Q16 | 31 | 29 | 17 | 87 | 3.13 | 0.077 |
| Misconceptions* | 1 Q7 | 6 | 23 | 11 | 124 | 4.235 | 0.04<sup>±</sup> |
|  | 2 Q5 | 7 | 31 | 6 | 120 | 16.892 | 3.96x10<sup>-05±</sup> |
|  | 2 Q6 | 23 | 47 | 5 | 89 | 33.923 | 5.73x10<sup>-09±</sup> |
|  | 2 Q8 | 28 | 35 | 8 | 93 | 16.953 | 3.83x10<sup>-05±</sup> |
|  | 3 Q2 | 28 | 23 | 12 | 101 | 3.457 | 0.063 |
|  | 4 Q9 | 7 | 19 | 5 | 133 | 8.167 | 0.004<sup>±</sup> |
|  | 4 Q12 | 21 | 31 | 6 | 106 | 16.892 | 3.96x10<sup>-05±</sup> |
|  | 4 Q17 | 28 | 44 | 7 | 85 | 26.843 | 2.21x10<sup>-07±</sup> |
|  | 4 Q20 | 8 | 23 | 4 | 129 | 13.37 | 2.56x10<sup>-04±</sup> |
|  | 5 Q14 | 17 | 30 | 8 | 109 | 12.737 | 3.59x10<sup>-04±</sup> |
|  | 5 Q19 | 9 | 15 | 14 | 126 | 0.034 | 0.853 |
|  | 5 Q22 | 10 | 23 | 9 | 122 | 6.125 | 0.013<sup>±</sup> |
|  | 6 Q11 | 4 | 23 | 7 | 130 | 8.533 | 0.003<sup>±</sup> |
|  | 6 Q18 | 2 | 14 | 4 | 144 | 5.556 | 0.018<sup>±</sup> |
|  | 6 Q21 | 5 | 13 | 2 | 144 | 8.067 | 0.005<sup>±</sup> |

<sup>±</sup> Significant p-values
\* Misconceptions (Price et al. 2014)

**Table 7. For most individual questions, participant post-recitation performance improved across classes following instruction either with or without Genie (cont.).**
| Evaluation | Question | Non-Genie 2017 |  |  |  | McNemar stat. | p-value |
| --- | --- | --- | --- | --- | --- | --- | --- |
|  |  | Incorrect to incorrect switches | Incorrect to correct switches | Correct to incorrect switches | Correct to correct switches |  |  |
| Key concepts | Q1 | 0 | 13 | 5 | 139 | 3.556 | 0.059 |
|  | Q3 | 4 | 16 | 8 | 129 | 2.667 | 0.102 |
|  | Q15 | 2 | 18 | 5 | 132 | 7.348 | 0.007<sup>±</sup> |
|  | Q4 | 13 | 14 | 11 | 119 | 0.36 | 0.549 |
|  | Q10 | 6 | 18 | 4 | 129 | 8.909 | 0.003<sup>±</sup> |
|  | Q13 | 8 | 39 | 3 | 107 | 30.857 | 2.78x10<sup>-08±</sup> |
|  | Q16 | 27 | 30 | 27 | 73 | 0.158 | 0.691 |
| Misconceptions* | 1 Q7 | 11 | 23 | 10 | 113 | 5.121 | 0.024<sup>±</sup> |
|  | 2 Q5 | 16 | 39 | 4 | 98 | 28.488 | 9.43x10<sup>-08±</sup> |
|  | 2 Q6 | 24 | 60 | 5 | 68 | 46.538 | 8.98x10<sup>-12±</sup> |
|  | 2 Q8 | 36 | 44 | 8 | 69 | 24.923 | 5.97x10<sup>-07±</sup> |
|  | 3 Q2 | 43 | 20 | 20 | 74 | 0 | 1 |
|  | 4 Q9 | 10 | 20 | 6 | 121 | 7.538 | 0.006<sup>±</sup> |
|  | 4 Q12 | 32 | 34 | 11 | 80 | 11.756 | 6.07x10<sup>-04±</sup> |
|  | 4 Q17 | 34 | 33 | 4 | 86 | 22.730 | 1.86x10<sup>-06±</sup> |
|  | 4 Q20 | 9 | 15 | 4 | 129 | 6.368 | 0.012<sup>±</sup> |
|  | 5 Q14 | 21 | 32 | 12 | 92 | 9.091 | 0.003<sup>±</sup> |
|  | 5 Q19 | 16 | 26 | 11 | 104 | 6.081 | 0.014<sup>±</sup> |
|  | 5 Q22 | 16 | 23 | 6 | 112 | 9.966 | 0.002<sup>±</sup> |
|  | 6 Q11 | 6 | 32 | 5 | 114 | 19.703 | 9.05x10<sup>-06±</sup> |
|  | 6 Q18 | 1 | 16 | 6 | 134 | 4.545 | 0.033<sup>±</sup> |
|  | 6 Q21 | 10 | 18 | 12 | 117 | 1.200 | 0.273 |

<sup>±</sup> Significant p-values
\* Misconceptions (Price et al. 2014)
1. Genetic drift is unpredictable because it has a random component. 2. Genetic drift is natural selection/adaptation/acclimation to the environment that may result from a need to survive. 3. Genetic drift is not evolution because it does not lead to directional change that increases fitness. 4. Natural selection is always the most powerful mechanism of evolution, and it is the primary agent of evolutionary change. 5. Genetic drift is random mutation. 6. Genetic drift is gene flow or migration.

**Figure 3.**
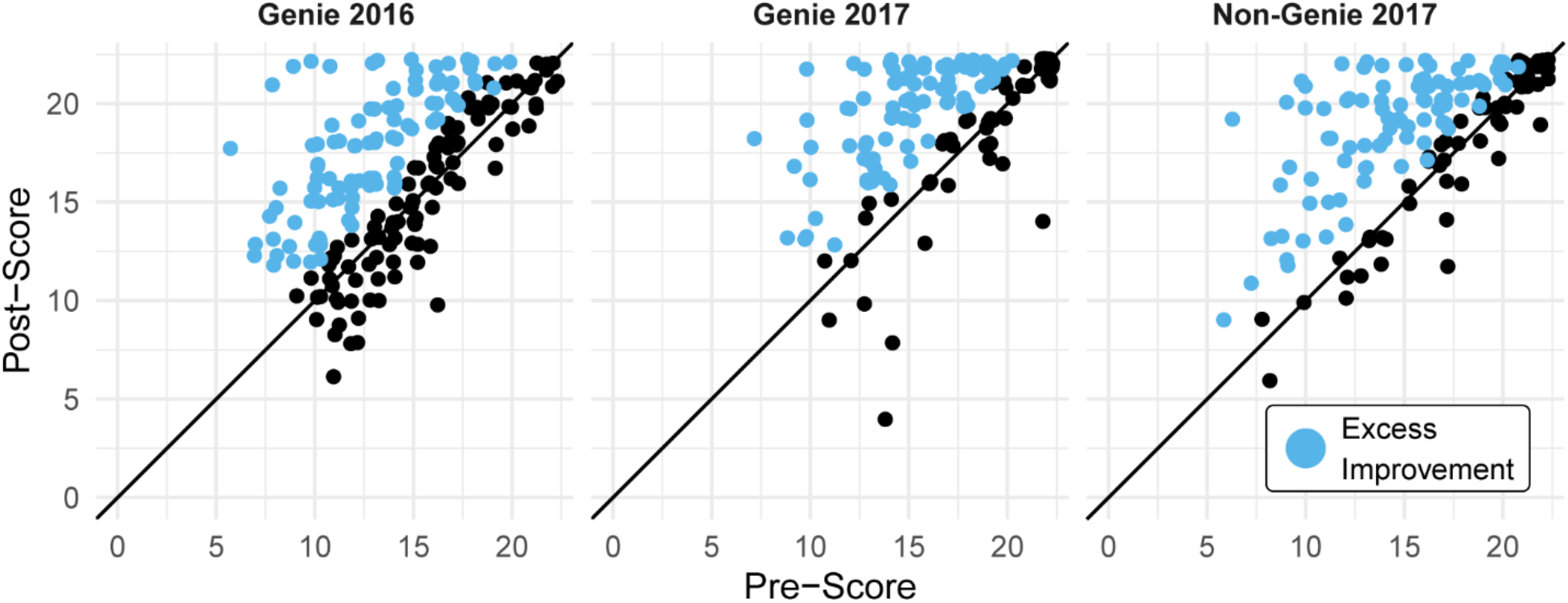
Students’ test scores generally improved after instruction. Blue dots represent excess improvement in class performance. The presence of blue points in a graph indicates that there were more students whose post-test score was better than their pre-test score. The number of blue points indicates how many more students improved their scores than students whose scores decreased.

**Figure 4.**
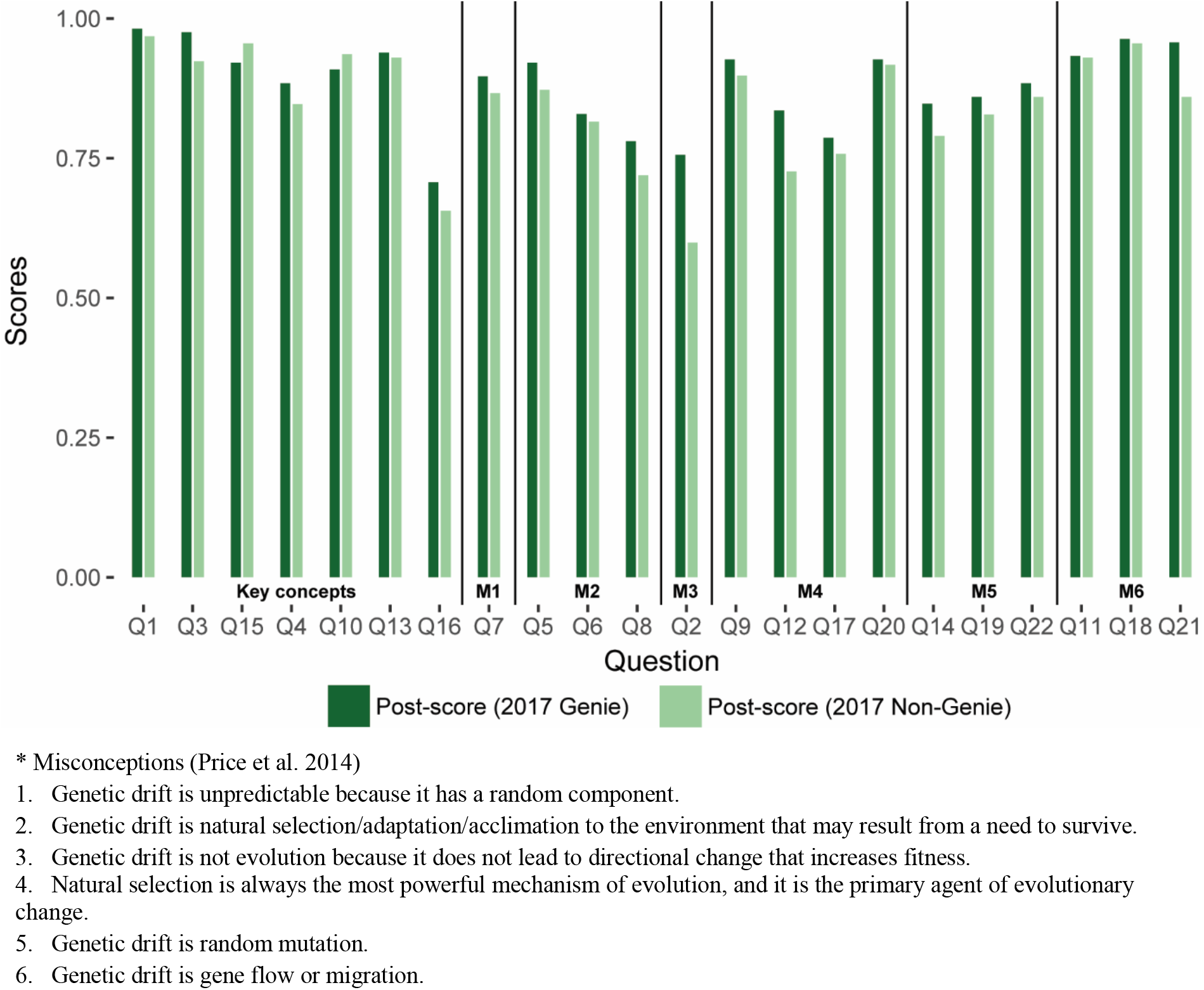
Post-recitation scores by question (Price et al. 2014) were generally higher in Genie 2017 compared to Non-Genie 2017. A bar plot comparing Genie 2017 (dark green) and Non-Genie 2017 (pale green) is shown. Questions have been grouped according to the classification provided by Price et al. (2014), with questions pertaining to Key concepts and misconceptions (M1-M6) separated by horizontal bars.

**Figure 5.**
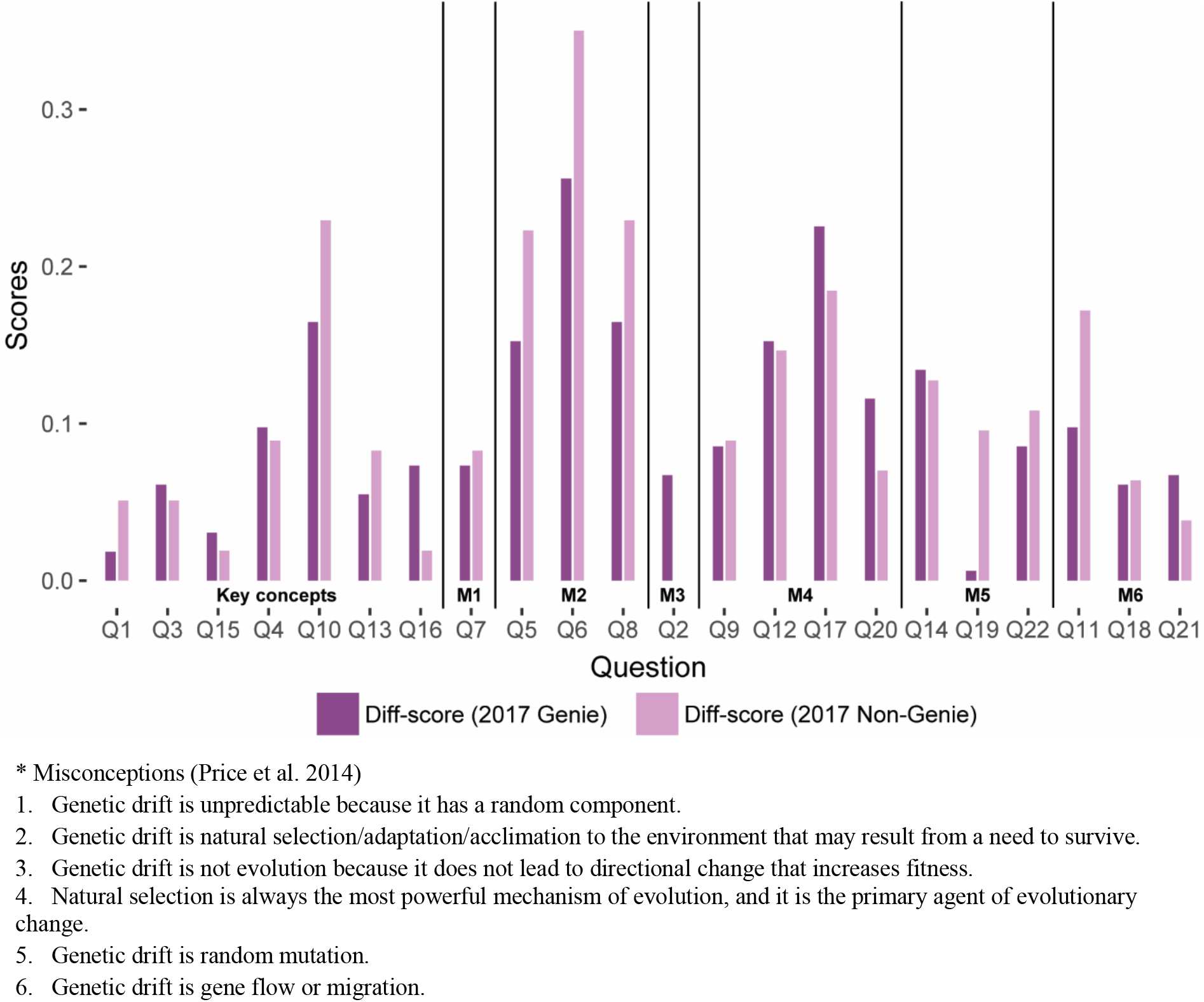
The difference between pre- and post-recitation scores by question (Price et al. 2014) shows that some questions saw higher improvement in Non-Genie 2017 while other showed higher improvement in Genie 2017. A bar plot comparing Genie 2017 (dark purple) and Non-Genie 2017 (pale purple) is shown. Questions have been grouped according to the classification provided by Price et al. (2014), with questions pertaining to Key concepts and misconceptions (M1-M6) separated by horizontal bars.

**Table 8.**
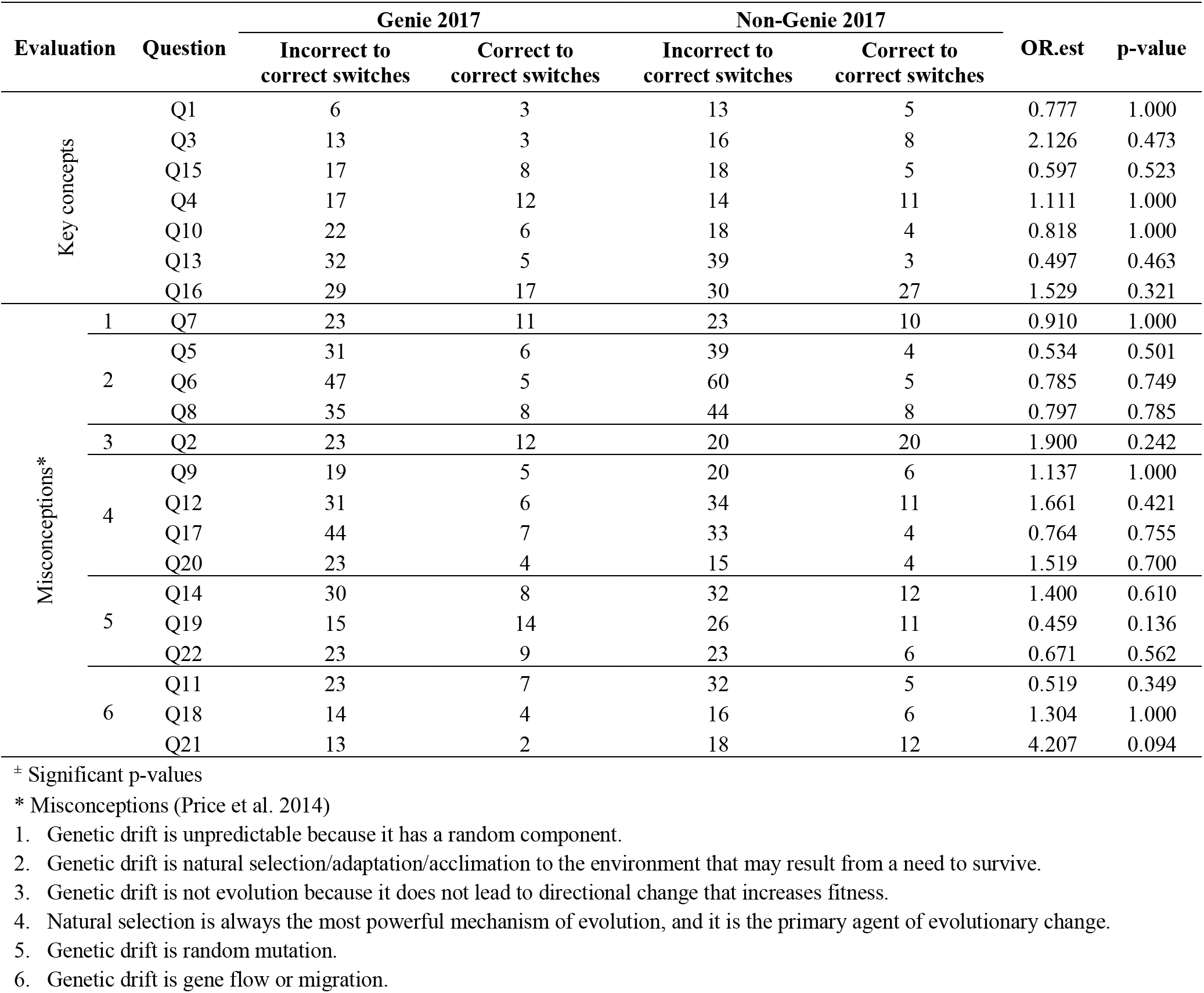
Comparison of performance between Genie 2017 and Non-Genie 2017, controlling by question. Fisher’s exact test testing the association between switches from ‘Incorrect to Correct’ and ‘Correct to Incorrect’ answers per question and by method of instruction (Genie 2017 and Non-Genie 2017).

## 4. Discussion

There are numerous other software capable of generating genetic drift simulations. Some of them can be easily downloaded and installed (Kliman et al. 2008; Revell 2019), others include an ample array of parameters to be modified by the user (http://evolution.gs.washington.edu/popgen/popg.html), and others can be found publicly available online (e.g. the Genetic Drift Simulator (http://www.biology.arizona.edu/evolution/act/drift/drift.html or Phyletica (http://phyletica.org/teaching/drift-simulator/). Some of this software even have a dynamic interface similar to that developed by Genie (http://virtualbiologylab.org/NetWebHTML_FilesJan2016/RandomEffectsModel.html). One notable software (Avida-ED) has a dynamic presentation and accessibility (https://avida-ed.msu.edu/app/AvidaED.html) alike Genie, but has additional parameters that permit for the evaluation of complex evolutionary hypotheses. While this list is not exhaustive, it provides a glimpse on how computational tools, and especially those found freely in web-interfaces, are becoming predominantly used in science teaching. The objective of this paper is not to compare Genie’s performance to all these tools, instead, the authors aim to present an additional teaching tool that can be added to an instructor’s repertoire. As such, we endeavor to show that Genie can be efficiently used alongside teacher-centered class instruction. A comparison was made between Genie-based instruction and instruction using static images (henceforth referred as teacher-centered instruction). The comparison was chosen since teacher-centered methods still are commonly used in science teaching (Tanner and Allen 2004) and have been traditionally used when teaching evolutionary topics in ASU.

There were no significant differences in the performance levels among participants from distinct demographic backgrounds. Despite differences in levels of representation across groups (e.g. more ‘Female’ than ‘Male’ participants in both Genie 2016 and 2017), pre- and post-recitation scores were similar. This suggests that Genie was as an effective teaching tool regardless of participant’s demographics. However, it should be mentioned that the variable ‘First-generation’ college students had statistical effect on participant’s performance. While participant’s performance increased in all methods of instruction, ‘First-generation’ college students showed slightly lower improvement than ‘Non-first generation’ college students. Multiple studies have attempted to address the social class gap among undergraduate students and explain why ‘First-generation’ college students, in occasion, perform more poorly than ‘Non-first generation’ college students (Grineski et al. 2018; Tibbetts et al. 2018). One finding pertinent to our assessment is that ‘First-generation’ college students tend to underperform when they know that their performance is going to be compared to that of other students in the class (Jury et al. 2015). This might be an unintended consequence of the in-class methods used here, which favored in-class discussion and student participation. However, while not possible to address here, is possible that these results point towards the unique disadvantages and social-related pressures that ‘First-generation’ college students face within ASU. These results should be evaluated in more detail in future studies.

We were unable to control for previous classes that BIO345 students took. Although, all students in BIO345 are required to have passed BIO340 (General Genetics), which typically includes instruction in evolutionary genetics; BIO340 is taught my multiple instructors, who do not teach evolutionary genetics equally. However, participants’ performance in all classes increased following instruction. Interestingly, mean scores showed that the increase in performance between pre- and post-recitation was ~0.1 regardless of the teaching method used. The main distinction were the pre-recitation scores, with some classes initially performing better than others. Taken together, these results are indicative that both teacher-centered and Genie-based teaching strategies led to a comparable improvement in participant’s scores, regardless of the initial performance level of the class. Thus, it is possible to conclude that Genie can perform as efficiently as traditionally teacher-centered instruction. In addition, when participant groups were divided based on the initial performance within their class (pre-recitation performance quantiles) both instruction methods lead to an increase in post-recitation scores. However, some quantiles saw more improvement than others. The highest improvement was observed in participants from the second pre-recitation score quantile in Genie 2016 and the third pre-recitation score quantile in 2017. On the other hand, participants with the highest pre-recitation scores (fourth quantile), showed the smallest improvement. The latter trend was more evident between Genie 2017 and Non-Genie 2017, with fourth quantile participants taking Genie-based classes having a slightly larger improvement than participants taking teacher-centered classes. This suggest that participants with initial lower understanding of the material similarly benefited from instruction with Genie as with other methods, while participants with initial higher understanding of the material benefited a little more when Genie was used.

In addition, participants’ performance was not affected by the instructor or the participant populations within the group, except for the ‘TA Pair1 7:30pm’ group during Genie 2016. The ‘TA Pair1 7:30pm’ group was formed by a small number of honors students; therefore, it is possible that this group simply performed better compared to the general class population in Genie 2016. Previous studies have found that instructors’ mastery of the content, as well as their overall teaching style play a critical role in students’ learning process (Alsharif and Qi 2014; Maleki et al. 2017). Thus, our results are indicative that Genie performs similarly well even with teachers using diverse teaching styles and having variable levels of expertise.

Overall, understanding of genetic drift key concepts and misconceptions improved following instruction with all teaching strategies. Nonetheless, it should be noted that there were small variations in how well participant’s performed in individual questions when using teaching-centered vs. Genie-based methods. Within 2017, teaching-centered methods performed better in questions related to key concepts on genetic drift as well as in understanding the difference between genetic drift and other non-adaptive or adaptive evolutionary mechanisms. On the other hand, Genie-based teaching aid the participants’ understanding on the importance of genetic drift as a source for evolutionary change and its relationship with natural selection. These results suggest that a combined teaching strategy using Genie alongside with traditional teacher-centered methods might help participants in gaining a more rounded comprehension of genetic drift concepts. In agreement with this conclusion, previous analyses have shown that a combination of traditional teaching-centered methods, with student-centered methods, and active learning strategies results in superior student performance (Dolan and Collins 2015; Shir et al. 2016; Wieman 2014).

In the case of evolution teaching, strategies that favor student’s development of critical thinking skills are especially useful. For instance, tools and methods that aid in creating and testing hypotheses have been effective in improving student’s understanding and acceptance of evolutionary theory (Lark et al. 2018; Smith et al. 2016). Likewise, instruction using computer simulation programs has proven to be valuable in facilitating student’s recognition of the breadth of evolutionary mechanisms that can act in a population (Kliman 2001). In that regard, flexible software that can be used to illustrate how diverse evolutionary forces are intrinsically linked can be particularly useful in teaching. For example, the award-winning software Avida-ED, has been effectively used to introduce evolutionary ideas to freshmen students via hypothesis testing (Abi Abdallah et al. 2020), understand the role of low-impact mutations in evolution (Nelson and Sanford 2011), and to test the robustness of genetic drift in small populations (Labar and Adami 2017). Nonetheless, it should not be forgotten than different students can master the same topic using different paths (Price et al. 2016), and that different classroom settings might be more suitable for distinct teaching methods. Simply put, there is no ‘fit all’ teaching strategy that can be universally implemented. Therefore, providing instructors with broad repertoire of teaching tools can aid them in finding those that better work for the topic being instructed, the specific class needs, and the instructor style. In this regard, we expect Genie can be added to the repertoire of higher education tools to be used for teaching genetic drift and other non-adaptive evolution concepts.

## 5. Conclusion

The present study shows that Genie can be successfully used for teaching undergraduate students concepts related with genetic drift and non-adaptive evolution. Genie performed comparably to traditional teacher-centered methods across all evaluated groups. Moreover, Genie-based and teacher-centered approaches led to participants understanding distinct key concepts and misconceptions of genetic drift. This indicates that Genie can be effectively used alongside other teaching strategies to provide a rounded view of non-adaptive evolution. In a related note, Genie provides a mean for participants to develop and test their own hypotheses, which can be useful in practicing critical thinking skills.

## Ethics approval and consent to participate

The study was approved by IRB protocol: STUDY00003707.

## Consent for publication

Not applicable.

## Availability of data and materials

A previous version of this manuscript is available as preprint (https://doi.org/10.1101/268672). All data and code used has been made available as supplementary materials. Genie is publicly available at https://cartwrig.ht/apps/genie/

## Authors’ contributions

AIC wrote and edited the manuscript, performed the statistical analyses, and participated in in-class instruction. BHR designed and wrote Genie. MSR revised the manuscript, designed the class study, and participated in in-class instruction. RAC designed and wrote Genie, edited the manuscript, and the performed statistical analyses. MAW edited the manuscript, designed the class study, and participated in in-class instruction prior to recitations using Genie.

## Acknowledgements

The authors would like to thank the students of Spring 2016 and Spring 2016, BIO345 course at ASU for their participation on this study.

