## Supplementary material for "Genie: An interactive real-time simulation for teaching genetic drift": SFile2_Genie.R

```
#install.packages("fitdistrplus")
#install.packages("logspline")
#install.packages("effsize")
#install.packages("betareg")
#install.packages("ggplot2")
#install.packages("PairedData")
#install.packages("tidyr")
#install.packages("tidyverse")

#Please change this path to your working directory
setwd("/Users/Andreina/Desktop/Genie_new/manuscript_all")

library(fitdistrplus)
library(logspline)
library(effsize)
library(betareg)
library(ggplot2)
library(PairedData)
library(RColorBrewer)
library(ggpubr)
library(dplyr)
options(tidyverse.quiet = TRUE)
library(tidyverse)
library(glue)
display.brewer.all(n=NULL, type="all", select=NULL, exact.n=TRUE, 
                   colorblindFriendly=TRUE)


#Demographic analyses
demographics_2016 <- read.table("Sfile3_BIO345_DEMOGRAPHICS_2016.txt", header = TRUE, sep = "\t")
head(demographics_2016)
demographics_2017_GENIE <- read.table("Sfile4_BIO345_DEMOGRAPHICS_2017_GENIE.txt", header = TRUE, sep = "\t")
head(demographics_2017_GENIE)
demographics_2017_NOGENIE <- read.table("Sfile5_BIO345_DEMOGRAPHICS_2017_NOGENIE.txt", header = TRUE, sep = "\t")
head(demographics_2017_NOGENIE)


require(gridExtra)
gen_2016 <- ggplot(demographics_2016, aes(x=Gender, y=score, color=time)) +
  labs(title="Genie 2016 demographics", x="Reported gender", y ="Scores")+
  theme_bw() + 
  theme(panel.border = element_blank(), panel.grid.major = element_blank(), panel.grid.minor = element_blank(), axis.line = element_line(colour = "black")) +
  theme(legend.position = "none") +
  geom_boxplot(position=position_dodge(0.8)) +
  geom_jitter(position = position_jitterdodge(jitter.height = 0.01))

gen_2017_GENIE <- ggplot(demographics_2017_GENIE, aes(x=Gender, y=score, color=time)) + 
  labs(title="Genie 2017 demographics", x="Reported gender", y="")+
  theme_bw() + 
  theme(panel.border = element_blank(), panel.grid.major = element_blank(), panel.grid.minor = element_blank(), axis.line = element_line(colour = "black")) +
  theme(legend.position = "none") +
  geom_boxplot(position=position_dodge(0.8)) +
  geom_jitter(position = position_jitterdodge(jitter.height = 0.01))

gen_2017_NOGENIE <- ggplot(demographics_2017_NOGENIE, aes(x=Gender, y=score, color=time)) + 
  labs(title="Non-Genie 2017 demographics", x="Reported gender", y="")+
  theme_bw() + 
  theme(panel.border = element_blank(), panel.grid.major = element_blank(), panel.grid.minor = element_blank(), axis.line = element_line(colour = "black")) +
  theme(legend.position = "none") +
  geom_boxplot(position=position_dodge(0.8)) +
  geom_jitter(position = position_jitterdodge(jitter.height = 0.01))

first_gen_2016 <- ggplot(demographics_2016, aes(x=First_Gen, y=score, color=time)) + 
  labs(x="First generation student", y ="Scores")+
  theme_bw() + 
  theme(panel.border = element_blank(), panel.grid.major = element_blank(), panel.grid.minor = element_blank(), axis.line = element_line(colour = "black")) +
  theme(legend.position = "none") +
  geom_boxplot(position=position_dodge(0.8)) +
  geom_jitter(position = position_jitterdodge(jitter.height = 0.01))

first_gen_2017_GENIE <- ggplot(demographics_2017_GENIE, aes(x=First_Gen, y=score, color=time)) + 
  labs(x="First generation student", y="")+
  theme_bw() + 
  theme(panel.border = element_blank(), panel.grid.major = element_blank(), panel.grid.minor = element_blank(), axis.line = element_line(colour = "black")) +
  theme(legend.position = "none") +
  geom_boxplot(position=position_dodge(0.8)) +
  geom_jitter(position = position_jitterdodge(jitter.height = 0.01))

first_gen_2017_NOGENIE <- ggplot(demographics_2017_NOGENIE, aes(x=First_Gen, y=score, color=time)) + 
  labs(x="First generation student", y="")+
  theme_bw() + 
  theme(panel.border = element_blank(), panel.grid.major = element_blank(), panel.grid.minor = element_blank(), axis.line = element_line(colour = "black")) +
  theme(legend.position = "none") +
  geom_boxplot(position=position_dodge(0.8)) +
  geom_jitter(position = position_jitterdodge(jitter.height = 0.01))

ethn_2016 <- ggplot(demographics_2016, aes(x=Ethnicity, y=score, color=time)) + 
  labs(x="Reported ethnicity", y ="Scores")+
  theme_bw() + 
  theme(panel.border = element_blank(), panel.grid.major = element_blank(), panel.grid.minor = element_blank(), axis.line = element_line(colour = "black")) +
  theme(legend.position = "none") +
  geom_boxplot(position=position_dodge(0.8)) +
  geom_jitter(position = position_jitterdodge(jitter.height = 0.01))

ethn_2017_GENIE <- ggplot(demographics_2017_GENIE, aes(x=Ethnicity, y=score, color=time)) + 
  labs(x="Reported ethnicity", y="")+
  theme_bw() + 
  theme(panel.border = element_blank(), panel.grid.major = element_blank(), panel.grid.minor = element_blank(), axis.line = element_line(colour = "black")) +
  theme(legend.position = "none") +
  geom_boxplot(position=position_dodge(0.8)) +
  geom_jitter(position = position_jitterdodge(jitter.height = 0.01))

ethn_2017_NOGENIE <- ggplot(demographics_2017_NOGENIE, aes(x=Ethnicity, y=score, color=time)) + 
  labs(x="Reported ethnicity", y="")+
  theme_bw() + 
  theme(panel.border = element_blank(), panel.grid.major = element_blank(), panel.grid.minor = element_blank(), axis.line = element_line(colour = "black")) +
  theme(legend.position = "none") +
  geom_boxplot(position=position_dodge(0.8)) +
  geom_jitter(position = position_jitterdodge(jitter.height = 0.01))

pdf(file="F1.pdf")

F1 <- ggarrange(gen_2016 + scale_color_brewer(palette="YlOrRd"), 
          gen_2017_GENIE + scale_color_brewer(palette="YlOrRd"), 
          gen_2017_NOGENIE + scale_color_brewer(palette="YlOrRd"), 
          first_gen_2016 + scale_color_brewer(palette="YlOrRd"), 
          first_gen_2017_GENIE + scale_color_brewer(palette="YlOrRd"), 
          first_gen_2017_NOGENIE + scale_color_brewer(palette="YlOrRd"), 
          ethn_2016 + scale_color_brewer(palette="YlOrRd"), 
          ethn_2017_GENIE + scale_color_brewer(palette="YlOrRd"), 
          ethn_2017_NOGENIE + scale_color_brewer(palette="YlOrRd"),
          nrow=3, ncol=3)

print(F1)
dev.off()

##############################################################################################
#Counting the number of invididuals per demographic group per class
year <- rep("2016", 406)
demographics_2016 <- cbind(demographics_2016, year)
demo2016 <- select(demographics_2016, time, Gender, First_Gen, Ethnicity, year)
year <- rep("2017G", 280)
demographics_2017_GENIE <- cbind(demographics_2017_GENIE, year)
demo2017G <- select(demographics_2017_GENIE, time, Gender, First_Gen, Ethnicity, year)
year <- rep("2017NG", 232)
demographics_2017_NOGENIE <- cbind(demographics_2017_NOGENIE, year)
demo2017NG <- select(demographics_2017_NOGENIE, time, Gender, First_Gen, Ethnicity, year)

total <- rbind(demo2016, demo2017G, demo2017NG)

Ethinicy_contingency_table <- table(total$Ethnicity, total$year)
Gender_contingency_table <- table(total$Gender, total$year)
First_Gen_contingency_table <- table(total$First_Gen, total$year)

#Writing output to text
sink('T1_demographic_output.txt', append=TRUE)
Ethinicy_contingency_table
Gender_contingency_table
First_Gen_contingency_table
sink()

############################################################################################
#Are scores demographic dependent on either 2016 or 2017 classes?
demographics_2016_glm <- read.table("Sfile6_BIO345_DEMOGRAPHICS_2016.txt", header = TRUE, sep = "\t")
head(demographics_2016_glm)
demographics_2017_glm <- read.table("Sfile7_BIO345_DEMOGRAPHICS_2017.txt", header = TRUE, sep = "\t")
head(demographics_2017_glm)

#Pre-recitation scores 2016
glm_demo_2016_pre <- lm(score_pre ~ First_Gen * Ethnicity * Gender, data=demographics_2016_glm)
anova(glm_demo_2016_pre)
#Post-recitation scores 2016
glm_demo_2016_post <- lm(score_post ~ First_Gen * Ethnicity * Gender, data=demographics_2016_glm)
anova(glm_demo_2016_post)
#Pre-recitation scores 2017
glm_demo_2017_pre <- lm(score_pre ~ First_Gen * Ethnicity * Gender * Genie_used, data=demographics_2017_glm)
anova(glm_demo_2017_pre)
#Post-recitation scores 2017
glm_demo_2017_post <- lm(score_post ~ First_Gen * Ethnicity * Gender * Genie_used, data=demographics_2017_glm)
anova(glm_demo_2017_post)


#Writing output to text
sink('T2_anova_output.txt', append=TRUE)
cat("ANOVA on pre-recitation scores 2016\n")
anova(glm_demo_2016_pre)
cat("\n\n")
cat("=============================\n")
cat("ANOVA on post-recitation scores 2016\n")
anova(glm_demo_2016_post)
cat("\n\n")
cat("=============================\n")
cat("ANOVA on pre-recitation scores 2017\n")
anova(glm_demo_2017_pre)
cat("\n\n")
cat("=============================\n")
cat("ANOVA on post-recitation scores 2017\n")
anova(glm_demo_2017_post)
sink()


############################################################################################
#Input the results of both years and treatments
combined_years <- read.table("Sfile8_COMBINED_TABLE_FOR_DF.txt", header = TRUE, sep = "\t")
head(combined_years)

#Splits the data frame into years and treatments
split_years <- split(combined_years, combined_years$year)
df_2016 <- split_years[[1]]
head(df_2016)
df_2017_G <- split_years[[2]]
head(df_2017_G)
df_2017_NG <- split_years[[3]]
head(df_2017_NG)

#Fitting the data to a distribution
descdist(df_2016$score_pre, discrete = F) #beta distributionn
descdist(df_2016$score_post, discrete = F) #beta distribution
fit.beta_pre_2016 <- fitdist(df_2016$score_pre, "beta", method = "mme")
fit.beta_post_2016 <- fitdist(df_2016$score_post, "beta", method = "mme")
plot(fit.beta_pre_2016)
plot(fit.beta_post_2016)

descdist(df_2017_G$score_pre, discrete = F) #beta distributionn
descdist(df_2017_G$score_post, discrete = F) #beta distribution
fit.beta_pre_2017_G <- fitdist(df_2017_G$score_pre, "beta", method = "mme")
fit.beta_post_2017_G <- fitdist(df_2017_G$score_post, "beta", method = "mme")
plot(fit.beta_pre_2017_G)
plot(fit.beta_post_2017_G)

descdist(df_2017_NG$score_pre, discrete = F) #beta distributionn
descdist(df_2017_NG$score_post, discrete = F) #beta distribution
fit.beta_pre_2017_NG <- fitdist(df_2017_NG$score_pre, "beta", method = "mme")
fit.beta_post_2017_NG <- fitdist(df_2017_NG$score_post, "beta", method = "mme")
plot(fit.beta_pre_2017_NG)
plot(fit.beta_post_2017_NG)


############################################################################################
#Beta regression models
#Function to transform y values to be used on a betareg distribution
y.transf.betareg <- function(y){
  n.obs <- sum(!is.na(y))
  (y * (n.obs - 1) + 0.5) / n.obs
}

#Are the pre-recitation scores different across years (use of Genie or not)?
effect_combined_years_pre <- betareg(y.transf.betareg(score_pre) ~  year, data=combined_years, link = "logit")
summary(effect_combined_years_pre)
#Are the post-recitation scores across years dependent of the pre-recitation scores and/or the year of instruction (use of Genie or not)?
effect_combined_years <- betareg(y.transf.betareg(score_post) ~ score_pre + year, data=combined_years, link = "logit")
summary(effect_combined_years)
#Are the post-recitation scores on 2016 dependent of the pre-recitation scores and/or the class section?
effect_test_2016 <- betareg(y.transf.betareg(score_post) ~ score_pre + class, data=df_2016, link = "logit")
summary(effect_test_2016)
#Are the post-recitation scores on 2017 dependent of the pre-recitation scores and/or the class section when using Genie?
effect_test_2017_G <- betareg(y.transf.betareg(score_post) ~ score_pre + class, data=df_2017_G, link = "logit")
summary(effect_test_2017_G)
#Are the post-recitation scores on 2017 dependent of the pre-recitation scores and/or the class section when not using Genie?
effect_test_2017_NG <- betareg(y.transf.betareg(score_post) ~ score_pre + class, data=df_2017_NG, link = "logit")
summary(effect_test_2017_NG)


#Writing output to text
sink('T3_betareg_output.txt', append=TRUE)
cat("Betareg on pre-recitation scores across years\n")
summary(effect_combined_years_pre)
cat("\n\n")
cat("=============================\n")
cat("Betareg on pre-recitation scores across years\n")
summary(effect_combined_years)
cat("\n\n")
cat("=============================\n")
cat("Betareg on postrecitation scores 2016\n")
summary(effect_test_2016)
cat("\n\n")
cat("=============================\n")
cat("Betareg on postrecitation scores 2017 Genie\n")
summary(effect_test_2017_G)
cat("\n\n")
cat("=============================\n")
cat("Betareg on postrecitation scores 2017 Non-genie\n")
summary(effect_test_2017_NG)
sink()


############################################################################################
#Size effect testing
#Transforms the scores columns on each data frame into a numeric vector
score_post_2016 <- as.numeric(as.character(df_2016$score_post))
score_pre_2016 <- as.numeric(as.character(df_2016$score_pre))
score_diff_2016 <- as.numeric(as.character(df_2016$diff))

score_post_2017_G <- as.numeric(as.character(df_2017_G$score_post))
score_pre_2017_G <- as.numeric(as.character(df_2017_G$score_pre))
score_diff_2017_G <- as.numeric(as.character(df_2017_G$diff))

score_post_2017_NG <- as.numeric(as.character(df_2017_NG$score_post))
score_pre_2017_NG <- as.numeric(as.character(df_2017_NG$score_pre))
score_diff_2017_NG <- as.numeric(as.character(df_2017_NG$diff))

#Are the scores different before and after instruction on 2016 (Genie)? 
cohens_prepost_2016 <- cohen.d(score_post_2016,score_pre_2016)
#Are the scores different before and after instruction on 2017 (Genie)? 
cohens_prepost_2017G <- cohen.d(score_post_2017_G,score_pre_2017_G)
#Are the scores different before and after instruction on 2017 (No Genie)? 
cohens_prepost_2017NG <- cohen.d(score_post_2017_NG,score_pre_2017_NG)
#Are the pre-recitation scores different between 2017's Genie and Non Genie classes? 
cohens_pre_2017GNG <- cohen.d(score_pre_2017_G,score_pre_2017_NG)
#Are the post-recitation scores different between 2017's Genie and Non Genie classes? 
cohens_post_2017GNG <- cohen.d(score_post_2017_G,score_post_2017_NG)
#Are the difference in pre- and post-recitation scores different between 2017's Genie and Non Genie classes? 
cohens_diff_2017GNG <- cohen.d(score_diff_2017_G,score_diff_2017_NG)

mean(score_pre_2016)
mean(score_post_2016)
mean(score_pre_2017_G)
mean(score_post_2017_G)
mean(score_pre_2017_NG)
mean(score_post_2017_NG)

#Writing output to text
sink('T4_cohenD_output.txt', append=TRUE)
cat("Cohen's d for pre- vs. post-recitation 2016\n")
cohens_prepost_2016
cat("=============================\n")
cat("Cohen's d for pre- vs. post-recitation 2017 Genie\n")
cohens_prepost_2017G
cat("=============================\n")
cat("Cohen's d for pre- vs. post-recitation 2017 Non-genie\n")
cohens_prepost_2017NG
cat("=============================\n")
cat("Cohen's d for pre-recitation 2017 Genie vs. Non-genie\n")
cohens_pre_2017GNG
cat("=============================\n")
cat("Cohen's d for post-recitation 2017 Genie vs. Non-genie\n")
cohens_post_2017GNG
cat("=============================\n")
cat("Cohen's d for diff between pre- and post-recitation 2017 Genie vs. Non-genie\n")
cohens_diff_2017GNG
sink()


############################################################################################
#Paired Student t-test of pre- vs post-recitation performance
#2016
t.test(score_pre_2016, score_post_2016, paired = TRUE)
#2017G
t.test(score_pre_2017_G, score_post_2017_G, paired = TRUE)
#2017NG
t.test(score_pre_2017_NG, score_post_2017_NG, paired = TRUE)


#Writing output to text
sink('T5_paired_t_test.txt', append=TRUE)
cat("Paired Student t-test of pre- vs post-recitation performance in 2016\n")
t.test(score_pre_2016, score_post_2016, paired = TRUE)
cat("=============================\n")
cat("Paired Student t-test of pre- vs post-recitation performance in 2017 Genie\n")
t.test(score_pre_2017_G, score_post_2017_G, paired = TRUE)
cat("=============================\n")
cat("Paired Student t-test of pre- vs post-recitation performance in 2017 Non-genie\n")
t.test(score_pre_2017_NG, score_post_2017_NG, paired = TRUE)
sink()

############################################################################################
#Student t-test of performance by student quantiles
#Reorder individual data frames by the pre-recitation scores
df_2016_order_pre_score <- df_2016[order(df_2016$score_pre),] 
head(df_2016_order_pre_score)
df_2017_G_order_pre_score <- df_2017_G[order(df_2017_G$score_pre),] 
head(df_2017_G_order_pre_score)
df_2017_NG_order_pre_score <- df_2017_NG[order(df_2017_NG$score_pre),] 
head(df_2017_NG_order_pre_score)

#Split the dataframe into four quantiles based on pre-recitation scores
#2016
range = c(0, 0.25, 0.5, 0.75, 1)
df_2016_order_pre_score_quantiles <- split(df_2016_order_pre_score, cut(df_2016_order_pre_score$score_pre, range))
df_2016_order_pre_score_first_quantile <- df_2016_order_pre_score_quantiles[[1]]
df_2016_order_pre_score_first_quantile_numeric <- as.numeric(as.character(df_2016_order_pre_score_first_quantile$score_pre))
df_2016_order_post_score_first_quantile_numeric <- as.numeric(as.character(df_2016_order_pre_score_first_quantile$score_post))

df_2016_order_pre_score_second_quantile <- df_2016_order_pre_score_quantiles[[2]]
df_2016_order_pre_score_second_quantile_numeric <- as.numeric(as.character(df_2016_order_pre_score_second_quantile$score_pre))
df_2016_order_post_score_second_quantile_numeric <- as.numeric(as.character(df_2016_order_pre_score_second_quantile$score_post))

df_2016_order_pre_score_third_quantile <- df_2016_order_pre_score_quantiles[[3]]
df_2016_order_pre_score_third_quantile_numeric <- as.numeric(as.character(df_2016_order_pre_score_third_quantile$score_pre))
df_2016_order_post_score_third_quantile_numeric <- as.numeric(as.character(df_2016_order_pre_score_third_quantile$score_post))

df_2016_order_pre_score_fourth_quantile <- df_2016_order_pre_score_quantiles[[4]]
df_2016_order_pre_score_fourth_quantile_numeric <- as.numeric(as.character(df_2016_order_pre_score_fourth_quantile$score_pre))
df_2016_order_post_score_fourth_quantile_numeric <- as.numeric(as.character(df_2016_order_pre_score_fourth_quantile$score_post))

#2017 Genie classes
df_2017_G_order_pre_score_quantiles <- split(df_2017_G_order_pre_score, cut(df_2017_G_order_pre_score$score_pre, range))
df_2017_G_order_pre_score_first_quantile <- df_2017_G_order_pre_score_quantiles[[1]]
df_2017_G_order_pre_score_first_quantile_numeric <- as.numeric(as.character(df_2017_G_order_pre_score_first_quantile$score_pre))
df_2017_G_order_post_score_first_quantile_numeric <- as.numeric(as.character(df_2017_G_order_pre_score_first_quantile$score_post))

df_2017_G_order_pre_score_second_quantile <- df_2017_G_order_pre_score_quantiles[[2]]
df_2017_G_order_pre_score_second_quantile_numeric <- as.numeric(as.character(df_2017_G_order_pre_score_second_quantile$score_pre))
df_2017_G_order_post_score_second_quantile_numeric <- as.numeric(as.character(df_2017_G_order_pre_score_second_quantile$score_post))

df_2017_G_order_pre_score_third_quantile <- df_2017_G_order_pre_score_quantiles[[3]]
df_2017_G_order_pre_score_third_quantile_numeric <- as.numeric(as.character(df_2017_G_order_pre_score_third_quantile$score_pre))
df_2017_G_order_post_score_third_quantile_numeric <- as.numeric(as.character(df_2017_G_order_pre_score_third_quantile$score_post))

df_2017_G_order_pre_score_fourth_quantile <- df_2017_G_order_pre_score_quantiles[[4]]
df_2017_G_order_pre_score_fourth_quantile_numeric <- as.numeric(as.character(df_2017_G_order_pre_score_fourth_quantile$score_pre))
df_2017_G_order_post_score_fourth_quantile_numeric <- as.numeric(as.character(df_2017_G_order_pre_score_fourth_quantile$score_post))

#2017 No Genie classes
df_2017_NG_order_pre_score_quantiles <- split(df_2017_NG_order_pre_score, cut(df_2017_NG_order_pre_score$score_pre, range))
df_2017_NG_order_pre_score_first_quantile <- df_2017_NG_order_pre_score_quantiles[[1]]
df_2017_NG_order_pre_score_first_quantile_numeric <- as.numeric(as.character(df_2017_NG_order_pre_score_first_quantile$score_pre))
df_2017_NG_order_post_score_first_quantile_numeric <- as.numeric(as.character(df_2017_NG_order_pre_score_first_quantile$score_post))

df_2017_NG_order_pre_score_second_quantile <- df_2017_NG_order_pre_score_quantiles[[2]]
df_2017_NG_order_pre_score_second_quantile_numeric <- as.numeric(as.character(df_2017_NG_order_pre_score_second_quantile$score_pre))
df_2017_NG_order_post_score_second_quantile_numeric <- as.numeric(as.character(df_2017_NG_order_pre_score_second_quantile$score_post))

df_2017_NG_order_pre_score_third_quantile <- df_2017_NG_order_pre_score_quantiles[[3]]
df_2017_NG_order_pre_score_third_quantile_numeric <- as.numeric(as.character(df_2017_NG_order_pre_score_third_quantile$score_pre))
df_2017_NG_order_post_score_third_quantile_numeric <- as.numeric(as.character(df_2017_NG_order_pre_score_third_quantile$score_post))

df_2017_NG_order_pre_score_fourth_quantile <- df_2017_NG_order_pre_score_quantiles[[4]]
df_2017_NG_order_pre_score_fourth_quantile_numeric <- as.numeric(as.character(df_2017_NG_order_pre_score_fourth_quantile$score_pre))
df_2017_NG_order_post_score_fourth_quantile_numeric <- as.numeric(as.character(df_2017_NG_order_pre_score_fourth_quantile$score_post))

#Do different groups of students have differences performances? 
#2016
t.test(df_2016_order_pre_score_first_quantile_numeric,df_2016_order_post_score_first_quantile_numeric,  paired = FALSE)
t.test(df_2016_order_pre_score_second_quantile_numeric,df_2016_order_post_score_second_quantile_numeric,  paired = FALSE)
t.test(df_2016_order_pre_score_third_quantile_numeric,df_2016_order_post_score_third_quantile_numeric,  paired = FALSE)
t.test(df_2016_order_pre_score_fourth_quantile_numeric,df_2016_order_post_score_fourth_quantile_numeric,  paired = FALSE)

#2017 Genie classes
t.test(df_2017_G_order_pre_score_first_quantile_numeric,df_2017_G_order_post_score_first_quantile_numeric,  paired = FALSE)
t.test(df_2017_G_order_pre_score_second_quantile_numeric,df_2017_G_order_post_score_second_quantile_numeric,  paired = FALSE)
t.test(df_2017_G_order_pre_score_third_quantile_numeric,df_2017_G_order_post_score_third_quantile_numeric,  paired = FALSE)
t.test(df_2017_G_order_pre_score_fourth_quantile_numeric,df_2017_G_order_post_score_fourth_quantile_numeric,  paired = FALSE)

#2017 No Genie classes
t.test(df_2017_NG_order_pre_score_first_quantile_numeric,df_2017_NG_order_post_score_first_quantile_numeric,  paired = FALSE)
t.test(df_2017_NG_order_pre_score_second_quantile_numeric,df_2017_NG_order_post_score_second_quantile_numeric,  paired = FALSE)
t.test(df_2017_NG_order_pre_score_third_quantile_numeric,df_2017_NG_order_post_score_third_quantile_numeric,  paired = FALSE)
t.test(df_2017_NG_order_pre_score_fourth_quantile_numeric,df_2017_NG_order_post_score_fourth_quantile_numeric,  paired = FALSE)


#Writing output to text
sink('T6_performance_by_quantile_output.txt', append=TRUE)
cat("Performance by quantile on 2016\n\n")
cat("First quantile\n")
t.test(df_2016_order_pre_score_first_quantile_numeric,df_2016_order_post_score_first_quantile_numeric,  paired = FALSE)
cat("Not enough observations\n\n")
cat("Second quantile")
t.test(df_2016_order_pre_score_second_quantile_numeric,df_2016_order_post_score_second_quantile_numeric,  paired = FALSE)
cat("Third quantile")
t.test(df_2016_order_pre_score_third_quantile_numeric,df_2016_order_post_score_third_quantile_numeric,  paired = FALSE)
cat("Fourth quantile")
t.test(df_2016_order_pre_score_fourth_quantile_numeric,df_2016_order_post_score_fourth_quantile_numeric,  paired = FALSE)
cat("===============================================\n")
cat("Performance by quantile on 2017 Genie\n\n")
cat("First quantile\n")
t.test(df_2017_G_order_pre_score_first_quantile_numeric,df_2017_G_order_post_score_first_quantile_numeric,  paired = FALSE)
cat("Not enough observations\n\n")
cat("Second quantile")
t.test(df_2017_G_order_pre_score_second_quantile_numeric,df_2017_G_order_post_score_second_quantile_numeric,  paired = FALSE)
cat("Third quantile")
t.test(df_2017_G_order_pre_score_third_quantile_numeric,df_2017_G_order_post_score_third_quantile_numeric,  paired = FALSE)
cat("Fourth quantile")
t.test(df_2017_G_order_pre_score_fourth_quantile_numeric,df_2017_G_order_post_score_fourth_quantile_numeric,  paired = FALSE)
cat("===============================================\n")
cat("Performance by quantile on 2017 Non-Genie\n\n")
cat("First quantile\n")
t.test(df_2017_NG_order_pre_score_first_quantile_numeric,df_2017_NG_order_post_score_first_quantile_numeric,  paired = FALSE)
cat("Not enough observations\n\n")
cat("Second quantile")
t.test(df_2017_NG_order_pre_score_second_quantile_numeric,df_2017_NG_order_post_score_second_quantile_numeric,  paired = FALSE)
cat("Third quantile")
t.test(df_2017_NG_order_pre_score_third_quantile_numeric,df_2017_NG_order_post_score_third_quantile_numeric,  paired = FALSE)
cat("Fourth quantile")
t.test(df_2017_NG_order_pre_score_fourth_quantile_numeric,df_2017_NG_order_post_score_fourth_quantile_numeric,  paired = FALSE)
sink()


############################################################################################
#Visualization of performance on each year via violin plots
VIOLINPLOT_2016_PRE_POST <- read.table("Sfile10_VIOLINPLOT_2016_PRE_POST.txt", header = TRUE, sep = "\t")
head(VIOLINPLOT_2016_PRE_POST)

VIOLINPLOT_2017_GENIE_PRE_POST <- read.table("Sfile11_VIOLINPLOT_2017_GENIE_PRE_POST.txt", header = TRUE, sep = "\t")
head(VIOLINPLOT_2017_GENIE_PRE_POST)

VIOLINPLOT_2017_NOGENIE_PRE_POST <- read.table("Sfile12_VIOLINPLOT_2017_NOGENIE_PRE_POST.txt", header = TRUE, sep = "\t")
head(VIOLINPLOT_2017_NOGENIE_PRE_POST)


require(gridExtra)
VIOLINPLOT_2016 <- ggplot(VIOLINPLOT_2016_PRE_POST, aes(x = year, y = score, fill=year)) +
  scale_y_continuous(limits = c(0, 1.1), breaks = seq(0, 1.1, by = 0.1)) +
  scale_x_discrete(labels=c(".Pre-recitation_score_2016" = "Pre-recitation", "Post-recitation_score_2016" = "Post-recitation")) + xlab("") +
  labs(title="Genie 2016", y ="Scores")+
  theme_bw() + 
  theme(panel.border = element_blank(), panel.grid.major = element_blank(), panel.grid.minor = element_blank(), axis.line = element_line(colour = "black")) +
  theme(legend.position = "none") +
  scale_fill_brewer(palette="Dark2") + 
  geom_violin(trim=FALSE) + 
  geom_jitter(shape=16, position=position_jitter(0.1), col="black")

VIOLINPLOT_2017_GENIE <- ggplot(VIOLINPLOT_2017_GENIE_PRE_POST, aes(x = year, y = score, fill=year)) +
  scale_y_continuous(limits = c(0, 1.1), breaks = seq(0, 1.1, by = 0.1)) +
  scale_x_discrete(labels=c(".Pre-recitation_score_2017" = "Pre-recitation", "Post-recitation_score_2017" = "Post-recitation")) + xlab("") +
  labs(title="Genie 2017", y ="Scores")+
  theme_bw() + 
  theme(panel.border = element_blank(), panel.grid.major = element_blank(), panel.grid.minor = element_blank(), axis.line = element_line(colour = "black")) +
  theme(legend.position = "none") +
  scale_fill_brewer(palette="Dark2") + 
  geom_violin(trim=FALSE) + 
  geom_jitter(shape=16, position=position_jitter(0.1), col="black")

VIOLINPLOT_2017_NOGENIE <- ggplot(VIOLINPLOT_2017_NOGENIE_PRE_POST, aes(x = year, y = score, fill=year)) +
  scale_y_continuous(limits = c(0, 1.1), breaks = seq(0, 1.1, by = 0.1)) +
  scale_x_discrete(labels=c(".Pre-recitation_score_2017" = "Pre-recitation", "Post-recitation_score_2017" = "Post-recitation")) + xlab("") +
  labs(title="Non-Genie 2017", y ="Scores")+
  theme_bw() + 
  theme(panel.border = element_blank(), panel.grid.major = element_blank(), panel.grid.minor = element_blank(), axis.line = element_line(colour = "black")) +
  theme(legend.position = "none") +
  scale_fill_brewer(palette="Dark2") + 
  geom_violin(trim=FALSE) + 
  geom_jitter(shape=16, position=position_jitter(0.1), col="black")

pdf(file="F2.pdf")

F2 <- ggarrange(VIOLINPLOT_2016, VIOLINPLOT_2017_GENIE, VIOLINPLOT_2017_NOGENIE, nrow=3)

print(F2)
dev.off()

#Paired student plot
pd <- position_dodge(0.01)

PAIRED_STUDENT_2016 <- ggplot(VIOLINPLOT_2016_PRE_POST, aes(x = year, y = score)) +
  scale_y_continuous(limits = c(0, 1.1), breaks = seq(0, 1.1, by = 0.1)) +
  scale_x_discrete(labels=c(".Pre-recitation_score_2016" = "Pre-recitation", "Post-recitation_score_2016" = "Post-recitation")) + xlab("") +
  labs(title="Genie 2016", y ="Scores")+
  theme_bw() + 
  theme(panel.border = element_blank(), panel.grid.major = element_blank(), panel.grid.minor = element_blank(), axis.line = element_line(colour = "black")) +
  theme(legend.position = "none") +
  geom_line(aes(group = student, color=student), position = pd) +
  geom_point(position = pd)

PAIRED_STUDENT_2017_GENIE <- ggplot(VIOLINPLOT_2017_GENIE_PRE_POST, aes(x = year, y = score)) +
  scale_y_continuous(limits = c(0, 1.1), breaks = seq(0, 1.1, by = 0.1)) +
  scale_x_discrete(labels=c(".Pre-recitation_score_2017" = "Pre-recitation", "Post-recitation_score_2017" = "Post-recitation")) + xlab("") +
  labs(title="Genie 2017", y ="")+
  theme_bw() + 
  theme(panel.border = element_blank(), panel.grid.major = element_blank(), panel.grid.minor = element_blank(), axis.line = element_line(colour = "black")) +
  theme(legend.position = "none") +
  geom_line(aes(group = student, color=student), position = pd) +
  geom_point(position = pd)

PAIRED_STUDENT_2017_NOGENIE <- ggplot(VIOLINPLOT_2017_NOGENIE_PRE_POST, aes(x = year, y = score)) +
  scale_y_continuous(limits = c(0, 1.1), breaks = seq(0, 1.1, by = 0.1)) +
  scale_x_discrete(labels=c(".Pre-recitation_score_2017" = "Pre-recitation", "Post-recitation_score_2017" = "Post-recitation")) + xlab("") +
  labs(title="Non-Genie 2017", y ="") +
  theme_bw() + 
  theme(panel.border = element_blank(), panel.grid.major = element_blank(), panel.grid.minor = element_blank(), axis.line = element_line(colour = "black")) +
  theme(legend.position = "none") +
  geom_line(aes(group = student, color=student), position = pd) +
  geom_point(position = pd)

pdf(file="FS13.pdf")

FS13 <- ggarrange(PAIRED_STUDENT_2016, PAIRED_STUDENT_2017_GENIE, PAIRED_STUDENT_2017_NOGENIE, ncol=3)

print(FS13)
dev.off()


############################################################################################
#Testing performance per question
rawdat <- read_tsv("Sfile9_RESULTS_COMPILED_BY_QUESTION.txt")

# Calculate McNemar's Test Statistic
# https://en.wikipedia.org/wiki/McNemar%27s_test
mcnemar_stat <- function(b,c) {
  (b-c)^2/(b+c)
}

# Reshape the input data so that each year-student-question has its own line.
# Each observation will have a Pre and Post Stat
dat <- rawdat %>% pivot_longer(!c(year,class,student)) %>%
  separate("name", c("quiz", "question"), sep="_") %>%
  mutate(question = as.integer(question)) %>%
  pivot_wider(names_from="quiz",values_from="value")

# Count for each year-question the interaction of pre and post.
# use these counts to measure McNemar's statistic and its pvalue.
datn1 <- dat %>% mutate(result = glue("ans_{Pre}_{Post}")) %>%
  count(year, question, result) %>%
  pivot_wider(names_from="result",values_from="n", values_fill=list(n=0L)) %>%
  mutate(mcnemar_stat =  mcnemar_stat(ans_1_0, ans_0_1),
         mcnemar_pvalue = pchisq(mcnemar_stat, 1L, lower.tail = FALSE))

# Save Results
write_tsv(datn1,"T7_mcnemar.tsv")


############################################################################################
#Comparing 2017 genie vs non-genie classes performance per question
rawdat <- read_tsv("Sfile9_RESULTS_COMPILED_BY_QUESTION.txt")

dat0 <- rawdat %>% pivot_longer(!c(year,class,student)) %>%
  separate("name", c("quiz", "question"), sep="_") %>%
  mutate(question = as.integer(question)) %>%
  pivot_wider(names_from="quiz",values_from="value")

write_tsv(dat0,"pre_and_post.tsv")

dat <- read_tsv("pre_and_post.tsv")

# Compare 2017 Genie and 2017 Not Genie on pre and post
# Use Fisher's exact tests to test the association between question switching and
# type of instruction.

dat1 <- dat %>% mutate(result = str_glue("ans_{Pre}_{Post}")) %>%
  count(year, question, result) %>%
  pivot_wider(names_from="result",values_from="n", values_fill=list(n=0L)) %>%
  filter( year %in% c("2017_Genie", "2017_No_Genie"))

dat2 <- dat1 %>% select(year,question,ans_0_1,ans_1_0) %>%
  pivot_wider(names_from="year", values_from=c(ans_0_1,ans_1_0))

f <- function(a,b,c,d) {
  m <- matrix(c(a,b,c,d),nrow=2)
  v <- fisher.test(m)
  data.frame(OR.est=v$estimate, p.value=v$p.value)
}

dat3 <- dat2 %>% rowwise() %>% summarize(f(ans_0_1_2017_Genie, ans_0_1_2017_No_Genie, ans_1_0_2017_Genie, ans_1_0_2017_No_Genie))
dat4 <- bind_cols(dat2,dat3)

write_tsv(dat4,"T8_genie_vs_non.tsv")

pal_okabe_ito <- c("#E69F00", "#56B4E9", "#009E73", "#F0E442", "#0072B2", "#D55E00", "#CC79A7", "#000000")

dat <- read_tsv("pre_and_post.tsv")


dat.A <- dat %>% mutate(result = str_glue("ans_{Pre}_{Post}")) %>%
  count(year, student, result) %>%
  pivot_wider(names_from="result",values_from="n", values_fill=list(n=0L)) %>%
  mutate(pre_acc = (ans_1_1+ans_1_0),
         post_acc = (ans_1_1+ans_0_1))
dat.B <- dat.A %>% mutate(diff = post_acc - pre_acc, diffs = sign(diff), excess = "no") %>%
  arrange(diff)

dat4 <- dat.B %>% group_by(year) %>% group_modify(function(tbl,y){
  j <- tail(1:nrow(tbl),sum(tbl$diffs))
  tbl$excess[j] <- "yes"
  tbl
})

pretty_year <- c(
  `2016_Genie` = "Genie 2016",
  `2017_Genie` = "Genie 2017",
  `2017_No_Genie` = "Non-Genie 2017"
)

ann_tbl <- tibble(pre_acc=10,post_acc=2.5,year="2017_No_Genie",excess="yes")

pdf(file="F3.pdf")

set.seed(0x90210)
gg <- ggplot(dat4, aes(x=pre_acc,y=post_acc,color=excess)) + geom_jitter(width=0.3,height=0.3)
gg <- gg + geom_abline(slope=1,intercept=0)
gg <- gg + labs(x = "Pre-Score", y = "Post-Score")
gg <- gg + theme_minimal() + coord_fixed(xlim=c(0,22),ylim=c(0,22))
gg <- gg + facet_wrap(~year, labeller = labeller(year = pretty_year))
gg <- gg + scale_color_manual(values=pal_okabe_ito[c(8,2)])
gg <- gg + theme(legend.position = "none")
gg <- gg + geom_label(data=ann_tbl,label="        Excess\n        Improvement",hjust="left", nudge_x=-2,color="black",size=3) +
  geom_point(data=ann_tbl,size=5)
print(gg)
dev.off()


############################################################################################
compiled_years_by_question <- read.table("Sfile9_RESULTS_COMPILED_BY_QUESTION.txt", header = TRUE, sep = "\t")
split_years_by_year <- split(compiled_years_by_question, compiled_years_by_question$year)
df_2016_by_year <- split_years_by_year[[1]]
df_2017_G_by_year <- split_years_by_year[[2]]
df_2017_NG_by_year <- split_years_by_year[[3]]

#Transforms the pre-recitation scores columns on each data frame into a numeric vector
score_pre1_2017_G <- as.numeric(as.character(df_2017_G_by_year$Pre_1))
score_pre2_2017_G <- as.numeric(as.character(df_2017_G_by_year$Pre_2))
score_pre3_2017_G <- as.numeric(as.character(df_2017_G_by_year$Pre_3))
score_pre4_2017_G <- as.numeric(as.character(df_2017_G_by_year$Pre_4))
score_pre5_2017_G <- as.numeric(as.character(df_2017_G_by_year$Pre_5))
score_pre6_2017_G <- as.numeric(as.character(df_2017_G_by_year$Pre_6))
score_pre7_2017_G <- as.numeric(as.character(df_2017_G_by_year$Pre_7))
score_pre8_2017_G <- as.numeric(as.character(df_2017_G_by_year$Pre_8))
score_pre9_2017_G <- as.numeric(as.character(df_2017_G_by_year$Pre_9))
score_pre10_2017_G <- as.numeric(as.character(df_2017_G_by_year$Pre_10))
score_pre11_2017_G <- as.numeric(as.character(df_2017_G_by_year$Pre_11))
score_pre12_2017_G <- as.numeric(as.character(df_2017_G_by_year$Pre_12))
score_pre13_2017_G <- as.numeric(as.character(df_2017_G_by_year$Pre_13))
score_pre14_2017_G <- as.numeric(as.character(df_2017_G_by_year$Pre_14))
score_pre15_2017_G <- as.numeric(as.character(df_2017_G_by_year$Pre_15))
score_pre16_2017_G <- as.numeric(as.character(df_2017_G_by_year$Pre_16))
score_pre17_2017_G <- as.numeric(as.character(df_2017_G_by_year$Pre_17))
score_pre18_2017_G <- as.numeric(as.character(df_2017_G_by_year$Pre_18))
score_pre19_2017_G <- as.numeric(as.character(df_2017_G_by_year$Pre_19))
score_pre20_2017_G <- as.numeric(as.character(df_2017_G_by_year$Pre_20))
score_pre21_2017_G <- as.numeric(as.character(df_2017_G_by_year$Pre_21))
score_pre22_2017_G <- as.numeric(as.character(df_2017_G_by_year$Pre_22))

score_pre1_2017_NG <- as.numeric(as.character(df_2017_NG_by_year$Pre_1))
score_pre2_2017_NG <- as.numeric(as.character(df_2017_NG_by_year$Pre_2))
score_pre3_2017_NG <- as.numeric(as.character(df_2017_NG_by_year$Pre_3))
score_pre4_2017_NG <- as.numeric(as.character(df_2017_NG_by_year$Pre_4))
score_pre5_2017_NG <- as.numeric(as.character(df_2017_NG_by_year$Pre_5))
score_pre6_2017_NG <- as.numeric(as.character(df_2017_NG_by_year$Pre_6))
score_pre7_2017_NG <- as.numeric(as.character(df_2017_NG_by_year$Pre_7))
score_pre8_2017_NG <- as.numeric(as.character(df_2017_NG_by_year$Pre_8))
score_pre9_2017_NG <- as.numeric(as.character(df_2017_NG_by_year$Pre_9))
score_pre10_2017_NG <- as.numeric(as.character(df_2017_NG_by_year$Pre_10))
score_pre11_2017_NG <- as.numeric(as.character(df_2017_NG_by_year$Pre_11))
score_pre12_2017_NG <- as.numeric(as.character(df_2017_NG_by_year$Pre_12))
score_pre13_2017_NG <- as.numeric(as.character(df_2017_NG_by_year$Pre_13))
score_pre14_2017_NG <- as.numeric(as.character(df_2017_NG_by_year$Pre_14))
score_pre15_2017_NG <- as.numeric(as.character(df_2017_NG_by_year$Pre_15))
score_pre16_2017_NG <- as.numeric(as.character(df_2017_NG_by_year$Pre_16))
score_pre17_2017_NG <- as.numeric(as.character(df_2017_NG_by_year$Pre_17))
score_pre18_2017_NG <- as.numeric(as.character(df_2017_NG_by_year$Pre_18))
score_pre19_2017_NG <- as.numeric(as.character(df_2017_NG_by_year$Pre_19))
score_pre20_2017_NG <- as.numeric(as.character(df_2017_NG_by_year$Pre_20))
score_pre21_2017_NG <- as.numeric(as.character(df_2017_NG_by_year$Pre_21))
score_pre22_2017_NG <- as.numeric(as.character(df_2017_NG_by_year$Pre_22))

#Gets score for each question
score_pre1_2017_G_val <- sum(score_pre1_2017_G)/length(score_pre1_2017_G)
score_pre2_2017_G_val <- sum(score_pre2_2017_G)/length(score_pre2_2017_G)
score_pre3_2017_G_val <- sum(score_pre3_2017_G)/length(score_pre3_2017_G)
score_pre4_2017_G_val <- sum(score_pre4_2017_G)/length(score_pre4_2017_G)
score_pre5_2017_G_val <- sum(score_pre5_2017_G)/length(score_pre5_2017_G)
score_pre6_2017_G_val <- sum(score_pre6_2017_G)/length(score_pre6_2017_G)
score_pre7_2017_G_val <- sum(score_pre7_2017_G)/length(score_pre7_2017_G)
score_pre8_2017_G_val <- sum(score_pre8_2017_G)/length(score_pre8_2017_G)
score_pre9_2017_G_val <- sum(score_pre9_2017_G)/length(score_pre9_2017_G)
score_pre10_2017_G_val <- sum(score_pre10_2017_G)/length(score_pre10_2017_G)
score_pre11_2017_G_val <- sum(score_pre11_2017_G)/length(score_pre11_2017_G)
score_pre12_2017_G_val <- sum(score_pre12_2017_G)/length(score_pre12_2017_G)
score_pre13_2017_G_val <- sum(score_pre13_2017_G)/length(score_pre13_2017_G)
score_pre14_2017_G_val <- sum(score_pre14_2017_G)/length(score_pre14_2017_G)
score_pre15_2017_G_val <- sum(score_pre15_2017_G)/length(score_pre15_2017_G)
score_pre16_2017_G_val <- sum(score_pre16_2017_G)/length(score_pre16_2017_G)
score_pre17_2017_G_val <- sum(score_pre17_2017_G)/length(score_pre17_2017_G)
score_pre18_2017_G_val <- sum(score_pre18_2017_G)/length(score_pre18_2017_G)
score_pre19_2017_G_val <- sum(score_pre19_2017_G)/length(score_pre19_2017_G)
score_pre20_2017_G_val <- sum(score_pre20_2017_G)/length(score_pre20_2017_G)
score_pre21_2017_G_val <- sum(score_pre21_2017_G)/length(score_pre21_2017_G)
score_pre22_2017_G_val <- sum(score_pre22_2017_G)/length(score_pre22_2017_G)

score_pre1_2017_NG_val <- sum(score_pre1_2017_NG)/length(score_pre1_2017_NG)
score_pre2_2017_NG_val <- sum(score_pre2_2017_NG)/length(score_pre2_2017_NG)
score_pre3_2017_NG_val <- sum(score_pre3_2017_NG)/length(score_pre3_2017_NG)
score_pre4_2017_NG_val <- sum(score_pre4_2017_NG)/length(score_pre4_2017_NG)
score_pre5_2017_NG_val <- sum(score_pre5_2017_NG)/length(score_pre5_2017_NG)
score_pre6_2017_NG_val <- sum(score_pre6_2017_NG)/length(score_pre6_2017_NG)
score_pre7_2017_NG_val <- sum(score_pre7_2017_NG)/length(score_pre7_2017_NG)
score_pre8_2017_NG_val <- sum(score_pre8_2017_NG)/length(score_pre8_2017_NG)
score_pre9_2017_NG_val <- sum(score_pre9_2017_NG)/length(score_pre9_2017_NG)
score_pre10_2017_NG_val <- sum(score_pre10_2017_NG)/length(score_pre10_2017_NG)
score_pre11_2017_NG_val <- sum(score_pre11_2017_NG)/length(score_pre11_2017_NG)
score_pre12_2017_NG_val <- sum(score_pre12_2017_NG)/length(score_pre12_2017_NG)
score_pre13_2017_NG_val <- sum(score_pre13_2017_NG)/length(score_pre13_2017_NG)
score_pre14_2017_NG_val <- sum(score_pre14_2017_NG)/length(score_pre14_2017_NG)
score_pre15_2017_NG_val <- sum(score_pre15_2017_NG)/length(score_pre15_2017_NG)
score_pre16_2017_NG_val <- sum(score_pre16_2017_NG)/length(score_pre16_2017_NG)
score_pre17_2017_NG_val <- sum(score_pre17_2017_NG)/length(score_pre17_2017_NG)
score_pre18_2017_NG_val <- sum(score_pre18_2017_NG)/length(score_pre18_2017_NG)
score_pre19_2017_NG_val <- sum(score_pre19_2017_NG)/length(score_pre19_2017_NG)
score_pre20_2017_NG_val <- sum(score_pre20_2017_NG)/length(score_pre20_2017_NG)
score_pre21_2017_NG_val <- sum(score_pre21_2017_NG)/length(score_pre21_2017_NG)
score_pre22_2017_NG_val <- sum(score_pre22_2017_NG)/length(score_pre22_2017_NG)


#Transforms the post-recitation scores columns on each data frame into a numeric vector
score_post1_2017_G <- as.numeric(as.character(df_2017_G_by_year$Post_1))
score_post2_2017_G <- as.numeric(as.character(df_2017_G_by_year$Post_2))
score_post3_2017_G <- as.numeric(as.character(df_2017_G_by_year$Post_3))
score_post4_2017_G <- as.numeric(as.character(df_2017_G_by_year$Post_4))
score_post5_2017_G <- as.numeric(as.character(df_2017_G_by_year$Post_5))
score_post6_2017_G <- as.numeric(as.character(df_2017_G_by_year$Post_6))
score_post7_2017_G <- as.numeric(as.character(df_2017_G_by_year$Post_7))
score_post8_2017_G <- as.numeric(as.character(df_2017_G_by_year$Post_8))
score_post9_2017_G <- as.numeric(as.character(df_2017_G_by_year$Post_9))
score_post10_2017_G <- as.numeric(as.character(df_2017_G_by_year$Post_10))
score_post11_2017_G <- as.numeric(as.character(df_2017_G_by_year$Post_11))
score_post12_2017_G <- as.numeric(as.character(df_2017_G_by_year$Post_12))
score_post13_2017_G <- as.numeric(as.character(df_2017_G_by_year$Post_13))
score_post14_2017_G <- as.numeric(as.character(df_2017_G_by_year$Post_14))
score_post15_2017_G <- as.numeric(as.character(df_2017_G_by_year$Post_15))
score_post16_2017_G <- as.numeric(as.character(df_2017_G_by_year$Post_16))
score_post17_2017_G <- as.numeric(as.character(df_2017_G_by_year$Post_17))
score_post18_2017_G <- as.numeric(as.character(df_2017_G_by_year$Post_18))
score_post19_2017_G <- as.numeric(as.character(df_2017_G_by_year$Post_19))
score_post20_2017_G <- as.numeric(as.character(df_2017_G_by_year$Post_20))
score_post21_2017_G <- as.numeric(as.character(df_2017_G_by_year$Post_21))
score_post22_2017_G <- as.numeric(as.character(df_2017_G_by_year$Post_22))

score_post1_2017_NG <- as.numeric(as.character(df_2017_NG_by_year$Post_1))
score_post2_2017_NG <- as.numeric(as.character(df_2017_NG_by_year$Post_2))
score_post3_2017_NG <- as.numeric(as.character(df_2017_NG_by_year$Post_3))
score_post4_2017_NG <- as.numeric(as.character(df_2017_NG_by_year$Post_4))
score_post5_2017_NG <- as.numeric(as.character(df_2017_NG_by_year$Post_5))
score_post6_2017_NG <- as.numeric(as.character(df_2017_NG_by_year$Post_6))
score_post7_2017_NG <- as.numeric(as.character(df_2017_NG_by_year$Post_7))
score_post8_2017_NG <- as.numeric(as.character(df_2017_NG_by_year$Post_8))
score_post9_2017_NG <- as.numeric(as.character(df_2017_NG_by_year$Post_9))
score_post10_2017_NG <- as.numeric(as.character(df_2017_NG_by_year$Post_10))
score_post11_2017_NG <- as.numeric(as.character(df_2017_NG_by_year$Post_11))
score_post12_2017_NG <- as.numeric(as.character(df_2017_NG_by_year$Post_12))
score_post13_2017_NG <- as.numeric(as.character(df_2017_NG_by_year$Post_13))
score_post14_2017_NG <- as.numeric(as.character(df_2017_NG_by_year$Post_14))
score_post15_2017_NG <- as.numeric(as.character(df_2017_NG_by_year$Post_15))
score_post16_2017_NG <- as.numeric(as.character(df_2017_NG_by_year$Post_16))
score_post17_2017_NG <- as.numeric(as.character(df_2017_NG_by_year$Post_17))
score_post18_2017_NG <- as.numeric(as.character(df_2017_NG_by_year$Post_18))
score_post19_2017_NG <- as.numeric(as.character(df_2017_NG_by_year$Post_19))
score_post20_2017_NG <- as.numeric(as.character(df_2017_NG_by_year$Post_20))
score_post21_2017_NG <- as.numeric(as.character(df_2017_NG_by_year$Post_21))
score_post22_2017_NG <- as.numeric(as.character(df_2017_NG_by_year$Post_22))

#Gets score for each question
score_post1_2017_G_val <- sum(score_post1_2017_G)/length(score_post1_2017_G)
score_post2_2017_G_val <- sum(score_post2_2017_G)/length(score_post2_2017_G)
score_post3_2017_G_val <- sum(score_post3_2017_G)/length(score_post3_2017_G)
score_post4_2017_G_val <- sum(score_post4_2017_G)/length(score_post4_2017_G)
score_post5_2017_G_val <- sum(score_post5_2017_G)/length(score_post5_2017_G)
score_post6_2017_G_val <- sum(score_post6_2017_G)/length(score_post6_2017_G)
score_post7_2017_G_val <- sum(score_post7_2017_G)/length(score_post7_2017_G)
score_post8_2017_G_val <- sum(score_post8_2017_G)/length(score_post8_2017_G)
score_post9_2017_G_val <- sum(score_post9_2017_G)/length(score_post9_2017_G)
score_post10_2017_G_val <- sum(score_post10_2017_G)/length(score_post10_2017_G)
score_post11_2017_G_val <- sum(score_post11_2017_G)/length(score_post11_2017_G)
score_post12_2017_G_val <- sum(score_post12_2017_G)/length(score_post12_2017_G)
score_post13_2017_G_val <- sum(score_post13_2017_G)/length(score_post13_2017_G)
score_post14_2017_G_val <- sum(score_post14_2017_G)/length(score_post14_2017_G)
score_post15_2017_G_val <- sum(score_post15_2017_G)/length(score_post15_2017_G)
score_post16_2017_G_val <- sum(score_post16_2017_G)/length(score_post16_2017_G)
score_post17_2017_G_val <- sum(score_post17_2017_G)/length(score_post17_2017_G)
score_post18_2017_G_val <- sum(score_post18_2017_G)/length(score_post18_2017_G)
score_post19_2017_G_val <- sum(score_post19_2017_G)/length(score_post19_2017_G)
score_post20_2017_G_val <- sum(score_post20_2017_G)/length(score_post20_2017_G)
score_post21_2017_G_val <- sum(score_post21_2017_G)/length(score_post21_2017_G)
score_post22_2017_G_val <- sum(score_post22_2017_G)/length(score_post22_2017_G)

score_post1_2017_NG_val <- sum(score_post1_2017_NG)/length(score_post1_2017_NG)
score_post2_2017_NG_val <- sum(score_post2_2017_NG)/length(score_post2_2017_NG)
score_post3_2017_NG_val <- sum(score_post3_2017_NG)/length(score_post3_2017_NG)
score_post4_2017_NG_val <- sum(score_post4_2017_NG)/length(score_post4_2017_NG)
score_post5_2017_NG_val <- sum(score_post5_2017_NG)/length(score_post5_2017_NG)
score_post6_2017_NG_val <- sum(score_post6_2017_NG)/length(score_post6_2017_NG)
score_post7_2017_NG_val <- sum(score_post7_2017_NG)/length(score_post7_2017_NG)
score_post8_2017_NG_val <- sum(score_post8_2017_NG)/length(score_post8_2017_NG)
score_post9_2017_NG_val <- sum(score_post9_2017_NG)/length(score_post9_2017_NG)
score_post10_2017_NG_val <- sum(score_post10_2017_NG)/length(score_post10_2017_NG)
score_post11_2017_NG_val <- sum(score_post11_2017_NG)/length(score_post11_2017_NG)
score_post12_2017_NG_val <- sum(score_post12_2017_NG)/length(score_post12_2017_NG)
score_post13_2017_NG_val <- sum(score_post13_2017_NG)/length(score_post13_2017_NG)
score_post14_2017_NG_val <- sum(score_post14_2017_NG)/length(score_post14_2017_NG)
score_post15_2017_NG_val <- sum(score_post15_2017_NG)/length(score_post15_2017_NG)
score_post16_2017_NG_val <- sum(score_post16_2017_NG)/length(score_post16_2017_NG)
score_post17_2017_NG_val <- sum(score_post17_2017_NG)/length(score_post17_2017_NG)
score_post18_2017_NG_val <- sum(score_post18_2017_NG)/length(score_post18_2017_NG)
score_post19_2017_NG_val <- sum(score_post19_2017_NG)/length(score_post19_2017_NG)
score_post20_2017_NG_val <- sum(score_post20_2017_NG)/length(score_post20_2017_NG)
score_post21_2017_NG_val <- sum(score_post21_2017_NG)/length(score_post21_2017_NG)
score_post22_2017_NG_val <- sum(score_post22_2017_NG)/length(score_post22_2017_NG)


#Vector of pre-recitation and post-recitation scores by question on 2017 genie classes
pre_recitation_scores_2017_G2AT3 <- c(score_pre1_2017_G_val,
                                      score_pre3_2017_G_val,
                                      score_pre15_2017_G_val,
                                      score_pre4_2017_G_val,
                                      score_pre10_2017_G_val,
                                      score_pre13_2017_G_val,
                                      score_pre16_2017_G_val,
                                      score_pre7_2017_G_val,
                                      score_pre5_2017_G_val,
                                      score_pre6_2017_G_val,
                                      score_pre8_2017_G_val,
                                      score_pre2_2017_G_val,
                                      score_pre9_2017_G_val,
                                      score_pre12_2017_G_val,
                                      score_pre17_2017_G_val,
                                      score_pre20_2017_G_val,
                                      score_pre14_2017_G_val,
                                      score_pre19_2017_G_val,
                                      score_pre22_2017_G_val,
                                      score_pre11_2017_G_val,
                                      score_pre18_2017_G_val,
                                      score_pre21_2017_G_val)


post_recitation_scores_2017_G2AT3 <- c(score_post1_2017_G_val,
                                       score_post3_2017_G_val,
                                       score_post15_2017_G_val,
                                       score_post4_2017_G_val,
                                       score_post10_2017_G_val,
                                       score_post13_2017_G_val,
                                       score_post16_2017_G_val,
                                       score_post7_2017_G_val,
                                       score_post5_2017_G_val,
                                       score_post6_2017_G_val,
                                       score_post8_2017_G_val,
                                       score_post2_2017_G_val,
                                       score_post9_2017_G_val,
                                       score_post12_2017_G_val,
                                       score_post17_2017_G_val,
                                       score_post20_2017_G_val,
                                       score_post14_2017_G_val,
                                       score_post19_2017_G_val,
                                       score_post22_2017_G_val,
                                       score_post11_2017_G_val,
                                       score_post18_2017_G_val,
                                       score_post21_2017_G_val)


pre_recitation_scores_2017_NG2AT3 <- c(score_pre1_2017_NG_val,
                                       score_pre3_2017_NG_val,
                                       score_pre15_2017_NG_val,
                                       score_pre4_2017_NG_val,
                                       score_pre10_2017_NG_val,
                                       score_pre13_2017_NG_val,
                                       score_pre16_2017_NG_val,
                                       score_pre7_2017_NG_val,
                                       score_pre5_2017_NG_val,
                                       score_pre6_2017_NG_val,
                                       score_pre8_2017_NG_val,
                                       score_pre2_2017_NG_val,
                                       score_pre9_2017_NG_val,
                                       score_pre12_2017_NG_val,
                                       score_pre17_2017_NG_val,
                                       score_pre20_2017_NG_val,
                                       score_pre14_2017_NG_val,
                                       score_pre19_2017_NG_val,
                                       score_pre22_2017_NG_val,
                                       score_pre11_2017_NG_val,
                                       score_pre18_2017_NG_val,
                                       score_pre21_2017_NG_val)


post_recitation_scores_2017_NG2AT3 <- c(score_post1_2017_NG_val,
                                        score_post3_2017_NG_val,
                                        score_post15_2017_NG_val,
                                        score_post4_2017_NG_val,
                                        score_post10_2017_NG_val,
                                        score_post13_2017_NG_val,
                                        score_post16_2017_NG_val,
                                        score_post7_2017_NG_val,
                                        score_post5_2017_NG_val,
                                        score_post6_2017_NG_val,
                                        score_post8_2017_NG_val,
                                        score_post2_2017_NG_val,
                                        score_post9_2017_NG_val,
                                        score_post12_2017_NG_val,
                                        score_post17_2017_NG_val,
                                        score_post20_2017_NG_val,
                                        score_post14_2017_NG_val,
                                        score_post19_2017_NG_val,
                                        score_post22_2017_NG_val,
                                        score_post11_2017_NG_val,
                                        score_post18_2017_NG_val,
                                        score_post21_2017_NG_val)


#get diifferece between pre- and post-recitation scores
score_diff1_2017_G <- score_post1_2017_G_val-score_pre1_2017_G_val
score_diff2_2017_G <- score_post2_2017_G_val-score_pre2_2017_G_val
score_diff3_2017_G <- score_post3_2017_G_val-score_pre3_2017_G_val
score_diff4_2017_G <- score_post4_2017_G_val-score_pre4_2017_G_val
score_diff5_2017_G <- score_post5_2017_G_val-score_pre5_2017_G_val
score_diff6_2017_G <- score_post6_2017_G_val-score_pre6_2017_G_val
score_diff7_2017_G <- score_post7_2017_G_val-score_pre7_2017_G_val
score_diff8_2017_G <- score_post8_2017_G_val-score_pre8_2017_G_val
score_diff9_2017_G <- score_post9_2017_G_val-score_pre9_2017_G_val
score_diff10_2017_G <- score_post10_2017_G_val-score_pre10_2017_G_val
score_diff11_2017_G <- score_post11_2017_G_val-score_pre11_2017_G_val
score_diff12_2017_G <- score_post12_2017_G_val-score_pre12_2017_G_val
score_diff13_2017_G <- score_post13_2017_G_val-score_pre13_2017_G_val
score_diff14_2017_G <- score_post14_2017_G_val-score_pre14_2017_G_val
score_diff15_2017_G <- score_post15_2017_G_val-score_pre15_2017_G_val
score_diff16_2017_G <- score_post16_2017_G_val-score_pre16_2017_G_val
score_diff17_2017_G <- score_post17_2017_G_val-score_pre17_2017_G_val
score_diff18_2017_G <- score_post18_2017_G_val-score_pre18_2017_G_val
score_diff19_2017_G <- score_post19_2017_G_val-score_pre19_2017_G_val
score_diff20_2017_G <- score_post20_2017_G_val-score_pre20_2017_G_val
score_diff21_2017_G <- score_post21_2017_G_val-score_pre21_2017_G_val
score_diff22_2017_G <- score_post22_2017_G_val-score_pre22_2017_G_val


diff_recitation_scores_2017_G2AT3 <- c(score_diff1_2017_G,
                                   score_diff3_2017_G,
                                   score_diff4_2017_G,
                                   score_diff10_2017_G,
                                   score_diff13_2017_G,
                                   score_diff15_2017_G,
                                   score_diff16_2017_G,
                                   score_diff7_2017_G,
                                   score_diff5_2017_G,
                                   score_diff6_2017_G,
                                   score_diff8_2017_G,
                                   score_diff2_2017_G,
                                   score_diff9_2017_G,
                                   score_diff12_2017_G,
                                   score_diff17_2017_G,
                                   score_diff20_2017_G,
                                   score_diff14_2017_G,
                                   score_diff19_2017_G,
                                   score_diff22_2017_G,
                                   score_diff11_2017_G,
                                   score_diff18_2017_G,
                                   score_diff21_2017_G)

score_diff1_2017_NG <- score_post1_2017_NG_val-score_pre1_2017_NG_val
score_diff2_2017_NG <- score_post2_2017_NG_val-score_pre2_2017_NG_val
score_diff3_2017_NG <- score_post3_2017_NG_val-score_pre3_2017_NG_val
score_diff4_2017_NG <- score_post4_2017_NG_val-score_pre4_2017_NG_val
score_diff5_2017_NG <- score_post5_2017_NG_val-score_pre5_2017_NG_val
score_diff6_2017_NG <- score_post6_2017_NG_val-score_pre6_2017_NG_val
score_diff7_2017_NG <- score_post7_2017_NG_val-score_pre7_2017_NG_val
score_diff8_2017_NG <- score_post8_2017_NG_val-score_pre8_2017_NG_val
score_diff9_2017_NG <- score_post9_2017_NG_val-score_pre9_2017_NG_val
score_diff10_2017_NG <- score_post10_2017_NG_val-score_pre10_2017_NG_val
score_diff11_2017_NG <- score_post11_2017_NG_val-score_pre11_2017_NG_val
score_diff12_2017_NG <- score_post12_2017_NG_val-score_pre12_2017_NG_val
score_diff13_2017_NG <- score_post13_2017_NG_val-score_pre13_2017_NG_val
score_diff14_2017_NG <- score_post14_2017_NG_val-score_pre14_2017_NG_val
score_diff15_2017_NG <- score_post15_2017_NG_val-score_pre15_2017_NG_val
score_diff16_2017_NG <- score_post16_2017_NG_val-score_pre16_2017_NG_val
score_diff17_2017_NG <- score_post17_2017_NG_val-score_pre17_2017_NG_val
score_diff18_2017_NG <- score_post18_2017_NG_val-score_pre18_2017_NG_val
score_diff19_2017_NG <- score_post19_2017_NG_val-score_pre19_2017_NG_val
score_diff20_2017_NG <- score_post20_2017_NG_val-score_pre20_2017_NG_val
score_diff21_2017_NG <- score_post21_2017_NG_val-score_pre21_2017_NG_val
score_diff22_2017_NG <- score_post22_2017_NG_val-score_pre22_2017_NG_val

diff_recitation_scores_2017_NG2AT3 <- c(score_diff1_2017_NG,
                                    score_diff3_2017_NG,
                                    score_diff4_2017_NG,
                                    score_diff10_2017_NG,
                                    score_diff13_2017_NG,
                                    score_diff15_2017_NG,
                                    score_diff16_2017_NG,
                                    score_diff7_2017_NG,
                                    score_diff5_2017_NG,
                                    score_diff6_2017_NG,
                                    score_diff8_2017_NG,
                                    score_diff2_2017_NG,
                                    score_diff9_2017_NG,
                                    score_diff12_2017_NG,
                                    score_diff17_2017_NG,
                                    score_diff20_2017_NG,
                                    score_diff14_2017_NG,
                                    score_diff19_2017_NG,
                                    score_diff22_2017_NG,
                                    score_diff11_2017_NG,
                                    score_diff18_2017_NG,
                                    score_diff21_2017_NG)
                                                       

#Questions on 2AT3 order in Price et al 2014
question2AT3 <- c("Q1","Q3","Q15","Q4","Q10","Q13","Q16","Q7","Q5","Q6","Q8","Q2","Q9","Q12","Q17","Q20","Q14","Q19","Q22","Q11","Q18","Q21")
time_diffG <- rep("Diff-score Genie", 22)
time_diffNG <- rep("Diff-score no Genie", 22)
df_diff_scores_by_question_GA2T3 <- data.frame(diff_recitation_scores = diff_recitation_scores_2017_G2AT3,
                                           question=question2AT3,
                                           time=time_diffG)

df_diff_scores_by_question_NGA2T3 <- data.frame(diff_recitation_scores = diff_recitation_scores_2017_NG2AT3,
                                            question=question2AT3,
                                            time=time_diffNG)

df_diff_scores_by_question_GA2T3$question <- factor(df_diff_scores_by_question_GA2T3$question,
                                                levels = df_diff_scores_by_question_GA2T3$question)

df_diff_scores_by_question_NGA2T3$question <- factor(df_diff_scores_by_question_NGA2T3$question,
                                                 levels = df_diff_scores_by_question_NGA2T3$question)

df_diff_scores_by_question_combinedA2T3 <- rbind(df_diff_scores_by_question_GA2T3, df_diff_scores_by_question_NGA2T3)

pdf(file="F5.pdf")

my_purple_colors <- c("orchid4","plum")
diff_per_question_plotA2T3 <- ggplot(data=df_diff_scores_by_question_combinedA2T3, aes(x=question, y=diff_recitation_scores, fill=time)) +
  geom_bar(width=0.4, stat="identity", position=position_dodge(0.5)) +
  scale_fill_manual(values = my_purple_colors) +
  xlab("Question") +
  ylab("Scores") +
  labs(fill = "Recitation") +
  scale_x_discrete(limits= df_diff_scores_by_question_combinedA2T3$p.question) +
  theme(panel.grid.major = element_blank(), panel.grid.minor = element_blank(),
          panel.background = element_blank(), axis.line = element_line(colour = "black"))

print(diff_per_question_plotA2T3)
dev.off()


question2AT3 <- c("Q1","Q3","Q15","Q4","Q10","Q13","Q16","Q7","Q5","Q6","Q8","Q2","Q9","Q12","Q17","Q20","Q14","Q19","Q22","Q11","Q18","Q21")
time_postG <- rep("Post-score Genie", 22)
time_postNG <- rep("Post-score no Genie", 22)
df_post_scores_by_question_GA2T3 <- data.frame(post_recitation_scores = post_recitation_scores_2017_G2AT3,
                                           question=question2AT3,
                                           time=time_postG)

df_post_scores_by_question_NGA2T3 <- data.frame(post_recitation_scores = post_recitation_scores_2017_NG2AT3,
                                            question=question2AT3,
                                            time=time_postNG)


df_post_scores_by_question_GA2T3$question <- factor(df_post_scores_by_question_GA2T3$question,
                                                levels = df_post_scores_by_question_GA2T3$question)

df_post_scores_by_question_NGA2T3$question <- factor(df_post_scores_by_question_NGA2T3$question,
                                                 levels = df_post_scores_by_question_NGA2T3$question)

df_post_scores_by_question_combinedA2T3 <- rbind(df_post_scores_by_question_GA2T3, df_post_scores_by_question_NGA2T3)


pdf(file="F4.pdf")

my_green_colors <- c("darkgreen","darkseagreen3")
post_per_question_plotA2T3 <- ggplot(data=df_post_scores_by_question_combinedA2T3, aes(x=question, y=post_recitation_scores, fill=time)) +
  geom_bar(width=0.4, stat="identity", position=position_dodge(0.5)) +
  #geom_text(aes(label=combined_diff_2017), vjust=1.6, color="red",
  #          position = position_dodge(0.9), size=3.5)+
  scale_fill_manual(values = my_green_colors) +
  xlab("Question") +
  ylab("Scores") +
  labs(fill = "Recitation") +
  scale_x_discrete(limits= df_post_scores_by_question_combinedA2T3$p.question)+
  theme(panel.grid.major = element_blank(), panel.grid.minor = element_blank(),
        panel.background = element_blank(), axis.line = element_line(colour = "black"))

print(post_per_question_plotA2T3)
dev.off()

#########################################################################################
#########################################################################################
#########################################################################################
#########################################################################################
#########################################################################################
#ANALYSIS REMOVING HONOR SECTIONS

#The results for both years and treatments removing the honors recitation section on each year
combined_years_without_honors <- combined_years[!(combined_years$class=="Pair1_730" & combined_years$year=="2016_Genie"),]
combined_years_without_honors <- combined_years_without_honors[!(combined_years_without_honors$class=="Pair2_130" & combined_years_without_honors$year=="2017_Genie"),]
head(combined_years_without_honors)

#Creates a violin plot of the difference in scores for each year and treatment
plotdiff <- ggplot(combined_years_without_honors, aes(x = year, y = diff, fill=year)) +
  scale_y_continuous(limits = c(0, 1), breaks = seq(0, 1, by = 0.1)) +
  scale_x_discrete() + xlab("Treatment Group") +
  ylab("Difference in recitation scores") + 
  theme(legend.position = "none") +
  scale_fill_brewer(palette="Paired") + 
  geom_violin(trim=FALSE) +  
  geom_jitter(shape=16, position=position_jitter(0.1), col="black")

plotdiff <- plotdiff + theme(panel.grid.major = element_blank(), panel.grid.minor = element_blank(),
                             panel.background = element_blank(), axis.line = element_line(colour = "black"))
plotdiff


############################################################################################
#Splits the data frame into years and treatments
split_years <- split(combined_years_without_honors, combined_years_without_honors$year)
df_2016_without_honors <- split_years[[1]]
df_2017_G_without_honors <- split_years[[2]]
df_2017_NG_without_honors <- split_years[[3]]

#Fitting the data to a distribution
descdist(df_2016$score_pre, discrete = F) #beta distributionn
descdist(df_2016$score_post, discrete = F) #beta distribution
fit.beta_pre_2016 <- fitdist(df_2016_without_honors$score_pre, "beta", method = "mme")
fit.beta_post_2016 <- fitdist(df_2016_without_honors$score_post, "beta", method = "mme")
plot(fit.beta_pre_2016)
plot(fit.beta_post_2016)

descdist(df_2017_G$score_pre, discrete = F) #beta distributionn
descdist(df_2017_G$score_post, discrete = F) #beta distribution
fit.beta_pre_2017_G <- fitdist(df_2017_G_without_honors$score_pre, "beta", method = "mme")
fit.beta_post_2017_G <- fitdist(df_2017_G_without_honors$score_post, "beta", method = "mme")
plot(fit.beta_pre_2017_G)
plot(fit.beta_post_2017_G)

descdist(df_2017_NG$score_pre, discrete = F) #beta distributionn
descdist(df_2017_NG$score_post, discrete = F) #beta distribution
fit.beta_pre_2017_NG <- fitdist(df_2017_NG_without_honors$score_pre, "beta", method = "mme")
fit.beta_post_2017_NG <- fitdist(df_2017_NG_without_honors$score_post, "beta", method = "mme")
plot(fit.beta_pre_2017_NG)
plot(fit.beta_post_2017_NG)


######Beta regression models#######
#Function to transform y values to be used on a betareg distribution
y.transf.betareg <- function(y){
  n.obs <- sum(!is.na(y))
  (y * (n.obs - 1) + 0.5) / n.obs
}

#Are the pre-recitation scores different across years (use of Genie or not)?
effect_combined_years_without_honors_pre <- betareg(y.transf.betareg(score_pre) ~  year, data=combined_years_without_honors, link = "logit")
#Are the post-recitation scores across years dependent of the pre-recitation scores and/or the year of instruction (use of Genie or not)?
effect_combined_years_without_honors <- betareg(y.transf.betareg(score_post) ~ score_pre + year, data=combined_years_without_honors, link = "logit")
#Are the post-recitation scores on 2016 dependent of the pre-recitation scores and/or the class section?
effect_test_2016_without_honors <- betareg(y.transf.betareg(score_post) ~ score_pre + class, data=df_2016_without_honors, link = "logit")
#Are the post-recitation scores on 2017 dependent of the pre-recitation scores and/or the class section when using Genie?
effect_test_2017_G_without_honors <- betareg(y.transf.betareg(score_post) ~ score_pre + class, data=df_2017_G_without_honors, link = "logit")
#Are the post-recitation scores on 2017 dependent of the pre-recitation scores and/or the class section when not using Genie?
effect_test_2017_NG_without_honors <- betareg(y.transf.betareg(score_post) ~ score_pre + class, data=df_2017_NG_without_honors, link = "logit")


#Writing output to text
sink('nohonors_betareg_output.txt', append=TRUE)
cat("Betareg on pre-recitation scores across years nohonors\n")
summary(effect_combined_years_without_honors_pre)
cat("\n\n")
cat("=============================\n")
cat("Betareg on pre-recitation scores across years nohonors\n")
summary(effect_combined_years_without_honors)
cat("\n\n")
cat("=============================\n")
cat("Betareg on postrecitation scores 2016 nohonors\n")
summary(effect_test_2016_without_honors)
cat("\n\n")
cat("=============================\n")
cat("Betareg on postrecitation scores 2017 Genie nohonors\n")
summary(effect_test_2017_G_without_honors)
cat("\n\n")
cat("=============================\n")
cat("Betareg on postrecitation scores 2017 Non-genie nohonors\n")
summary(effect_test_2017_NG_without_honors)
sink()

############################################################################################
#Size effect testing
#Transforms the scores columns on each data frame into a numeric vector
score_post_2016_without_honors <- as.numeric(as.character(df_2016_without_honors$score_post))
score_pre_2016_without_honors <- as.numeric(as.character(df_2016_without_honors$score_pre))
score_diff_2016_without_honors <- as.numeric(as.character(df_2016_without_honors$diff))

score_post_2017_G_without_honors <- as.numeric(as.character(df_2017_G_without_honors$score_post))
score_pre_2017_G_without_honors <- as.numeric(as.character(df_2017_G_without_honors$score_pre))
score_diff_2017_G_without_honors <- as.numeric(as.character(df_2017_G_without_honors$diff))

score_post_2017_NG_without_honors <- as.numeric(as.character(df_2017_NG_without_honors$score_post))
score_pre_2017_NG_without_honors <- as.numeric(as.character(df_2017_NG_without_honors$score_pre))
score_diff_2017_NG_without_honors <- as.numeric(as.character(df_2017_NG_without_honors$diff))


#Are the scores different before and after instruction on 2016 (Genie)? 
cohen.d(score_post_2016_without_honors,score_pre_2016_without_honors)
#Are the scores different before and after instruction on 2017 (Genie)? 
cohen.d(score_post_2017_G_without_honors,score_pre_2017_G_without_honors)
#Are the scores different before and after instruction on 2017 (No Genie)? 
cohen.d(score_post_2017_NG_without_honors,score_pre_2017_NG_without_honors)
#Are the pre-recitation scores different between 2017's Genie and Non Genie classes? 
cohen.d(score_pre_2017_G_without_honors,score_pre_2017_NG_without_honors)
#Are the post-recitation scores different between 2017's Genie and Non Genie classes? 
cohen.d(score_post_2017_G_without_honors,score_post_2017_NG_without_honors)
#Are the difference in pre- and post-recitation scores different between 2017's Genie and Non Genie classes? 
cohen.d(score_diff_2017_G_without_honors,score_diff_2017_NG_without_honors)


#Writing output to text
sink('nohonors_cohenD_output.txt', append=TRUE)
cat("Cohen's d for pre- vs. post-recitation 2016 nohonors\n")
cohen.d(score_post_2016_without_honors,score_pre_2016_without_honors)
cat("=============================\n")
cat("Cohen's d for pre- vs. post-recitation 2017 Genie nohonors\n")
cohen.d(score_post_2017_G_without_honors,score_pre_2017_G_without_honors)
cat("=============================\n")
cat("Cohen's d for pre- vs. post-recitation 2017 Non-genie nohonors\n")
cohen.d(score_post_2017_NG_without_honors,score_pre_2017_NG_without_honors)
cat("=============================\n")
cat("Cohen's d for pre-recitation 2017 Genie vs. Non-genie nohonors\n")
cohen.d(score_pre_2017_G_without_honors,score_pre_2017_NG_without_honors)
cat("=============================\n")
cat("Cohen's d for post-recitation 2017 Genie vs. Non-genie nohonors\n")
cohen.d(score_post_2017_G_without_honors,score_post_2017_NG_without_honors)
cat("=============================\n")
cat("Cohen's d for diff between pre- and post-recitation 2017 Genie vs. Non-genie nohonors\n")
cohen.d(score_diff_2017_G_without_honors,score_diff_2017_NG_without_honors)
sink()


############################################################################################
#Paired Student t-test of pre- vs post-recitation performance
#2016
t.test(score_pre_2016_without_honors, score_post_2016_without_honors, paired = TRUE)
#2017G
t.test(score_pre_2017_G_without_honors, score_post_2017_G_without_honors, paired = TRUE)
#2017NG
t.test(score_pre_2017_NG_without_honors, score_post_2017_NG_without_honors, paired = TRUE)

#Writing output to text
sink('nohonors_paired_t_test.txt', append=TRUE)
cat("Paired Student t-test of pre- vs post-recitation performance in 2016 nohonors\n")
t.test(score_pre_2016_without_honors, score_post_2016_without_honors, paired = TRUE)
cat("=============================\n")
cat("Paired Student t-test of pre- vs post-recitation performance in 2017 Genie nohonors\n")
t.test(score_pre_2017_G_without_honors, score_post_2017_G_without_honors, paired = TRUE)
cat("=============================\n")
cat("Paired Student t-test of pre- vs post-recitation performance in 2017 Non-genie nohonors\n")
t.test(score_pre_2017_NG_without_honors, score_post_2017_NG_without_honors, paired = TRUE)
sink()

############################################################################################
#Paired-test of performance by student quantiles
#Reorder individual data frames by the pre-recitation scores
df_2016_order_pre_score_without_honors <- df_2016_without_honors[order(df_2016_without_honors$score_pre),] 
df_2017_G_order_pre_score_without_honors <- df_2017_G_without_honors[order(df_2017_G_without_honors$score_pre),] 
df_2017_NG_order_pre_score_without_honors <- df_2017_NG_without_honors[order(df_2017_NG_without_honors$score_pre),] 

#Split the dataframe into four quantiles based on pre-recitation scores
#2016
range = c(0, 0.25, 0.5, 0.75, 1)
df_2016_order_pre_score_quantiles_without_honors <- split(df_2016_order_pre_score_without_honors, cut(df_2016_order_pre_score_without_honors$score_pre, range))
df_2016_order_pre_score_second_quantile_without_honors <- df_2016_order_pre_score_quantiles_without_honors[[2]]
df_2016_order_pre_score_second_quantile_numeric_without_honors <- as.numeric(as.character(df_2016_order_pre_score_second_quantile_without_honors$score_pre))
df_2016_order_post_score_second_quantile_numeric_without_honors <- as.numeric(as.character(df_2016_order_pre_score_second_quantile_without_honors$score_post))

df_2016_order_pre_score_third_quantile_without_honors <- df_2016_order_pre_score_quantiles_without_honors[[3]]
df_2016_order_pre_score_third_quantile_numeric_without_honors <- as.numeric(as.character(df_2016_order_pre_score_third_quantile_without_honors$score_pre))
df_2016_order_post_score_third_quantile_numeric_without_honors <- as.numeric(as.character(df_2016_order_pre_score_third_quantile_without_honors$score_post))

df_2016_order_pre_score_fourth_quantile_without_honors <- df_2016_order_pre_score_quantiles_without_honors[[4]]
df_2016_order_pre_score_fourth_quantile_numeric_without_honors <- as.numeric(as.character(df_2016_order_pre_score_fourth_quantile_without_honors$score_pre))
df_2016_order_post_score_fourth_quantile_numeric_without_honors <- as.numeric(as.character(df_2016_order_pre_score_fourth_quantile_without_honors$score_post))

#2017 Genie classes
df_2017_G_order_pre_score_quantiles_without_honors <- split(df_2017_G_order_pre_score_without_honors, cut(df_2017_G_order_pre_score_without_honors$score_pre, range))
df_2017_G_order_pre_score_second_quantile_without_honors <- df_2017_G_order_pre_score_quantiles_without_honors[[2]]
df_2017_G_order_pre_score_second_quantile_numeric_without_honors <- as.numeric(as.character(df_2017_G_order_pre_score_second_quantile_without_honors$score_pre))
df_2017_G_order_post_score_second_quantile_numeric_without_honors <- as.numeric(as.character(df_2017_G_order_pre_score_second_quantile_without_honors$score_post))

df_2017_G_order_pre_score_third_quantile_without_honors <- df_2017_G_order_pre_score_quantiles_without_honors[[3]]
df_2017_G_order_pre_score_third_quantile_numeric_without_honors <- as.numeric(as.character(df_2017_G_order_pre_score_third_quantile_without_honors$score_pre))
df_2017_G_order_post_score_third_quantile_numeric_without_honors <- as.numeric(as.character(df_2017_G_order_pre_score_third_quantile_without_honors$score_post))

df_2017_G_order_pre_score_fourth_quantile_without_honors <- df_2017_G_order_pre_score_quantiles_without_honors[[4]]
df_2017_G_order_pre_score_fourth_quantile_numeric_without_honors <- as.numeric(as.character(df_2017_G_order_pre_score_fourth_quantile_without_honors$score_pre))
df_2017_G_order_post_score_fourth_quantile_numeric_without_honors <- as.numeric(as.character(df_2017_G_order_pre_score_fourth_quantile_without_honors$score_post))

#2017 No Genie classes
df_2017_NG_order_pre_score_quantiles_without_honors <- split(df_2017_NG_order_pre_score_without_honors, cut(df_2017_NG_order_pre_score_without_honors$score_pre, range))
df_2017_NG_order_pre_score_second_quantile_without_honors <- df_2017_NG_order_pre_score_quantiles_without_honors[[2]]
df_2017_NG_order_pre_score_second_quantile_numeric_without_honors <- as.numeric(as.character(df_2017_NG_order_pre_score_second_quantile_without_honors$score_pre))
df_2017_NG_order_post_score_second_quantile_numeric_without_honors <- as.numeric(as.character(df_2017_NG_order_pre_score_second_quantile_without_honors$score_post))

df_2017_NG_order_pre_score_third_quantile_without_honors <- df_2017_NG_order_pre_score_quantiles_without_honors[[3]]
df_2017_NG_order_pre_score_third_quantile_numeric_without_honors <- as.numeric(as.character(df_2017_NG_order_pre_score_third_quantile_without_honors$score_pre))
df_2017_NG_order_post_score_third_quantile_numeric_without_honors <- as.numeric(as.character(df_2017_NG_order_pre_score_third_quantile_without_honors$score_post))

df_2017_NG_order_pre_score_fourth_quantile_without_honors <- df_2017_NG_order_pre_score_quantiles_without_honors[[4]]
df_2017_NG_order_pre_score_fourth_quantile_numeric_without_honors <- as.numeric(as.character(df_2017_NG_order_pre_score_fourth_quantile_without_honors$score_pre))
df_2017_NG_order_post_score_fourth_quantile_numeric_without_honors <- as.numeric(as.character(df_2017_NG_order_pre_score_fourth_quantile_without_honors$score_post))

#Do different groups of students have differences performances? 
#2016
t.test(df_2016_order_pre_score_second_quantile_numeric_without_honors,df_2016_order_post_score_second_quantile_numeric_without_honors,  paired = FALSE)
t.test(df_2016_order_pre_score_third_quantile_numeric_without_honors,df_2016_order_post_score_third_quantile_numeric_without_honors,  paired = FALSE)
t.test(df_2016_order_pre_score_fourth_quantile_numeric_without_honors,df_2016_order_post_score_fourth_quantile_numeric_without_honors,  paired = FALSE)

#2017 Genie classes
t.test(df_2017_G_order_pre_score_second_quantile_numeric_without_honors,df_2017_G_order_post_score_second_quantile_numeric_without_honors,  paired = FALSE)
t.test(df_2017_G_order_pre_score_third_quantile_numeric_without_honors,df_2017_G_order_post_score_third_quantile_numeric_without_honors,  paired = FALSE)
t.test(df_2017_G_order_pre_score_fourth_quantile_numeric_without_honors,df_2017_G_order_post_score_fourth_quantile_numeric_without_honors,  paired = FALSE)

#2017 No Genie classes
t.test(df_2017_NG_order_pre_score_second_quantile_numeric_without_honors,df_2017_NG_order_post_score_second_quantile_numeric_without_honors,  paired = FALSE)
t.test(df_2017_NG_order_pre_score_third_quantile_numeric_without_honors,df_2017_NG_order_post_score_third_quantile_numeric_without_honors,  paired = FALSE)
t.test(df_2017_NG_order_pre_score_fourth_quantile_numeric_without_honors,df_2017_NG_order_post_score_fourth_quantile_numeric_without_honors,  paired = FALSE)


#Writing output to text
sink('nohonors_performance_by_quantile_output.txt', append=TRUE)
cat("Performance by quantile on 2016\n\n")
cat("Second quantile")
t.test(df_2016_order_pre_score_second_quantile_numeric_without_honors,df_2016_order_post_score_second_quantile_numeric_without_honors,  paired = FALSE)
cat("Third quantile")
t.test(df_2016_order_pre_score_third_quantile_numeric_without_honors,df_2016_order_post_score_third_quantile_numeric_without_honors,  paired = FALSE)
cat("Fourth quantile")
t.test(df_2016_order_pre_score_fourth_quantile_numeric_without_honors,df_2016_order_post_score_fourth_quantile_numeric_without_honors,  paired = FALSE)
cat("===============================================\n")
cat("Performance by quantile on 2017 Genie\n\n")
cat("Second quantile")
t.test(df_2017_G_order_pre_score_second_quantile_numeric_without_honors,df_2017_G_order_post_score_second_quantile_numeric_without_honors,  paired = FALSE)
cat("Third quantile")
t.test(df_2017_G_order_pre_score_third_quantile_numeric_without_honors,df_2017_G_order_post_score_third_quantile_numeric_without_honors,  paired = FALSE)
cat("Fourth quantile")
t.test(df_2017_G_order_pre_score_fourth_quantile_numeric_without_honors,df_2017_G_order_post_score_fourth_quantile_numeric_without_honors,  paired = FALSE)
cat("===============================================\n")
cat("Performance by quantile on 2017 Non-Genie\n\n")
cat("Second quantile")
t.test(df_2017_NG_order_pre_score_second_quantile_numeric_without_honors,df_2017_NG_order_post_score_second_quantile_numeric_without_honors,  paired = FALSE)
cat("Third quantile")
t.test(df_2017_NG_order_pre_score_third_quantile_numeric_without_honors,df_2017_NG_order_post_score_third_quantile_numeric_without_honors,  paired = FALSE)
cat("Fourth quantile")
t.test(df_2017_NG_order_pre_score_fourth_quantile_numeric_without_honors,df_2017_NG_order_post_score_fourth_quantile_numeric_without_honors,  paired = FALSE)
sink()


############################################################################################
#Visualization of performance on each year via violin plots
VIOLINPLOT_2016_PRE_POST <- read.table("Sfile10_VIOLINPLOT_2016_PRE_POST.txt", header = TRUE, sep = "\t")
head(VIOLINPLOT_2016_PRE_POST)
VIOLINPLOT_2016_PRE_POST_without_honors <- VIOLINPLOT_2016_PRE_POST[!(VIOLINPLOT_2016_PRE_POST$class=="Pair1_730"),]
head(VIOLINPLOT_2016_PRE_POST_without_honors)

VIOLINPLOT_2017_GENIE_PRE_POST <- read.table("Sfile11_VIOLINPLOT_2017_GENIE_PRE_POST.txt", header = TRUE, sep = "\t")
head(VIOLINPLOT_2017_GENIE_PRE_POST)
VIOLINPLOT_2017_GENIE_PRE_POST_without_honors <- VIOLINPLOT_2017_GENIE_PRE_POST[!(VIOLINPLOT_2017_GENIE_PRE_POST$class=="Pair2_130"),]
head(VIOLINPLOT_2017_GENIE_PRE_POST_without_honors)

VIOLINPLOT_2017_NOGENIE_PRE_POST <- read.table("Sfile12_VIOLINPLOT_2017_NOGENIE_PRE_POST.txt", header = TRUE, sep = "\t")
head(VIOLINPLOT_2017_NOGENIE_PRE_POST)

require(gridExtra)
VIOLINPLOT_2016_without_honors <- ggplot(VIOLINPLOT_2016_PRE_POST_without_honors, aes(x = year, y = score, fill=year)) +
  scale_y_continuous(limits = c(0, 1.1), breaks = seq(0, 1.1, by = 0.1)) +
  scale_x_discrete() + xlab("") +
  ylab("2016") +
  theme(legend.position = "none") +
  scale_fill_brewer(palette="Dark2") + 
  geom_violin(trim=FALSE) + 
  geom_jitter(shape=16, position=position_jitter(0.1), col="black")

VIOLINPLOT_2016_without_honors <- VIOLINPLOT_2016_without_honors + theme(panel.grid.major = element_blank(), panel.grid.minor = element_blank(),
                                                                         panel.background = element_blank(), axis.line = element_line(colour = "black"))

VIOLINPLOT_2017_GENIE_without_honors <- ggplot(VIOLINPLOT_2017_GENIE_PRE_POST_without_honors, aes(x = year, y = score, fill=year)) +
  scale_y_continuous(limits = c(0, 1.1), breaks = seq(0, 1.1, by = 0.1)) +
  scale_x_discrete() + xlab("") +
  ylab("Evaluation scores 2017 (Genie)") +
  theme(legend.position = "none") +
  scale_fill_brewer(palette="Dark2") + 
  geom_violin(trim=FALSE) + 
  geom_jitter(shape=16, position=position_jitter(0.1), col="black")

VIOLINPLOT_2017_GENIE_without_honors <- VIOLINPLOT_2017_GENIE_without_honors + theme(panel.grid.major = element_blank(), panel.grid.minor = element_blank(),
                                                                                     panel.background = element_blank(), axis.line = element_line(colour = "black"))

VIOLINPLOT_2017_NOGENIE <- ggplot(VIOLINPLOT_2017_NOGENIE_PRE_POST, aes(x = year, y = score, fill=year)) +
  scale_y_continuous(limits = c(0, 1.1), breaks = seq(0, 1.1, by = 0.1)) +
  scale_x_discrete() + xlab("") +
  ylab("2017 (No Genie)") +
  theme(legend.position = "none") +
  scale_fill_brewer(palette="Dark2") + 
  geom_violin(trim=FALSE) + 
  geom_jitter(shape=16, position=position_jitter(0.1), col="black")


VIOLINPLOT_2017_NOGENIE <- VIOLINPLOT_2017_NOGENIE + theme(panel.grid.major = element_blank(), panel.grid.minor = element_blank(),
                                                           panel.background = element_blank(), axis.line = element_line(colour = "black"))


pdf(file="nohonors_F2.pdf")

F2 <- ggarrange(VIOLINPLOT_2016_without_honors, VIOLINPLOT_2017_GENIE_without_honors, VIOLINPLOT_2017_NOGENIE, nrow=3)

print(F2)
dev.off()


#Paired student plot
pd <- position_dodge(0.01)

PAIRED_STUDENT_2016_without_honors <- ggplot(VIOLINPLOT_2016_PRE_POST_without_honors, aes(x = year, y = score)) +
  scale_y_continuous(limits = c(0, 1.1), breaks = seq(0, 1.1, by = 0.1)) +
  scale_x_discrete(labels=c(".Pre-recitation_score_2016" = "Pre-recitation", "Post-recitation_score_2016" = "Post-recitation")) + xlab("") +
  labs(title="Genie 2016", y ="Scores")+
  theme_bw() + 
  theme(panel.border = element_blank(), panel.grid.major = element_blank(), panel.grid.minor = element_blank(), axis.line = element_line(colour = "black")) +
  theme(legend.position = "none") +
  geom_line(aes(group = student, color=student), position = pd) +
  geom_point(position = pd)

PAIRED_STUDENT_2017_GENIE_without_honors <- ggplot(VIOLINPLOT_2017_GENIE_PRE_POST_without_honors, aes(x = year, y = score)) +
  scale_y_continuous(limits = c(0, 1.1), breaks = seq(0, 1.1, by = 0.1)) +
  scale_x_discrete(labels=c(".Pre-recitation_score_2017" = "Pre-recitation", "Post-recitation_score_2017" = "Post-recitation")) + xlab("") +
  labs(title="Genie 2017", y ="")+
  theme_bw() + 
  theme(panel.border = element_blank(), panel.grid.major = element_blank(), panel.grid.minor = element_blank(), axis.line = element_line(colour = "black")) +
  theme(legend.position = "none") +
  geom_line(aes(group = student, color=student), position = pd) +
  geom_point(position = pd)

PAIRED_STUDENT_2017_NOGENIE <- ggplot(VIOLINPLOT_2017_NOGENIE_PRE_POST, aes(x = year, y = score)) +
  scale_y_continuous(limits = c(0, 1.1), breaks = seq(0, 1.1, by = 0.1)) +
  scale_x_discrete(labels=c(".Pre-recitation_score_2017" = "Pre-recitation", "Post-recitation_score_2017" = "Post-recitation")) + xlab("") +
  labs(title="Non-Genie 2017", y ="") +
  theme_bw() + 
  theme(panel.border = element_blank(), panel.grid.major = element_blank(), panel.grid.minor = element_blank(), axis.line = element_line(colour = "black")) +
  theme(legend.position = "none") +
  geom_line(aes(group = student, color=student), position = pd) +
  geom_point(position = pd)


pdf(file="nohonors_FS13.pdf")

FS13 <- ggarrange(PAIRED_STUDENT_2016_without_honors, PAIRED_STUDENT_2017_GENIE_without_honors, PAIRED_STUDENT_2017_NOGENIE, ncol=3)

print(FS13)
dev.off()


############################################################################################
#Performance by question
compiled_years_by_question <- read.table("Sfile9_RESULTS_COMPILED_BY_QUESTION.txt", header = TRUE, sep = "\t")
head(compiled_years_by_question)

#The results for both years and treatments removing the honors recitation section on each year
compiled_years_by_question_without_honors <- compiled_years_by_question[!(compiled_years_by_question$class=="Pair1_730" & compiled_years_by_question$year=="2016_Genie"),]
compiled_years_by_question_without_honors <- compiled_years_by_question_without_honors[!(compiled_years_by_question_without_honors$class=="Pair2_130" & compiled_years_by_question_without_honors$year=="2017_Genie"),]
head(compiled_years_by_question_without_honors)

split_years_by_year <- split(compiled_years_by_question_without_honors, compiled_years_by_question_without_honors$year)
df_2016_by_year_without_honors <- split_years_by_year[[1]]
head(df_2016_by_year_without_honors)
df_2017_G_by_year_without_honors <- split_years_by_year[[2]]
head(df_2017_G_by_year_without_honors)
df_2017_NG_by_year_without_honors <- split_years_by_year[[3]]
head(df_2017_NG_by_year_without_honors)

# Calculate McNemar's Test Statistic
# https://en.wikipedia.org/wiki/McNemar%27s_test
mcnemar_stat <- function(b,c) {
  (b-c)^2/(b+c)
}

# Reshape the input data so that each year-student-question has its own line.
# Each observation will have a Pre and Post Stat
dat <- compiled_years_by_question_without_honors %>% pivot_longer(!c(year,class,student)) %>%
  separate("name", c("quiz", "question"), sep="_") %>%
  mutate(question = as.integer(question)) %>%
  pivot_wider(names_from="quiz",values_from="value")

# Count for each year-question the interaction of pre and post.
# use these counts to measure McNemar's statistic and its pvalue.
dat1 <- dat %>% mutate(result = glue("ans_{Pre}_{Post}")) %>%
  count(year, question, result) %>%
  pivot_wider(names_from="result",values_from="n", values_fill=list(n=0L)) %>%
  mutate(mcnemar_stat =  mcnemar_stat(ans_1_0, ans_0_1),
         mcnemar_pvalue = pchisq(mcnemar_stat, 1L, lower.tail = FALSE))

# Save Results
write_tsv(dat1,"nohonors_mcnemar.tsv")


############################################################################################
#Comparing 2017 genie vs non-genie classes performance per question
rawdat <- read.table("Sfile9_RESULTS_COMPILED_BY_QUESTION.txt", header = TRUE, sep = "\t")
head(rawdat)

#The results for both years and treatments removing the honors recitation section on each year
compiled_years_by_question_without_honors <- rawdat[!(rawdat$class=="Pair1_730" & rawdat$year=="2016_Genie"),]
compiled_years_by_question_without_honors <- rawdat[!(rawdat$class=="Pair2_130" & rawdat$year=="2017_Genie"),]
head(compiled_years_by_question_without_honors)

dat0 <- compiled_years_by_question_without_honors %>% pivot_longer(!c(year,class,student)) %>%
  separate("name", c("quiz", "question"), sep="_") %>%
  mutate(question = as.integer(question)) %>%
  pivot_wider(names_from="quiz",values_from="value")

write_tsv(dat0,"nohonors_pre_and_post.tsv")

dat <- read_tsv("nohonors_pre_and_post.tsv")

# Compare 2017 Genie and 2017 Not Genie on pre and post
# Use Fisher's exact tests to test the association between question switching and
# type of instruction.

dat1 <- dat %>% mutate(result = str_glue("ans_{Pre}_{Post}")) %>%
  count(year, question, result) %>%
  pivot_wider(names_from="result",values_from="n", values_fill=list(n=0L)) %>%
  filter( year %in% c("2017_Genie", "2017_No_Genie"))

dat2 <- dat1 %>% select(year,question,ans_0_1,ans_1_0) %>%
  pivot_wider(names_from="year", values_from=c(ans_0_1,ans_1_0))

f <- function(a,b,c,d) {
  m <- matrix(c(a,b,c,d),nrow=2)
  v <- fisher.test(m)
  data.frame(OR.est=v$estimate, p.value=v$p.value)
}

dat3 <- dat2 %>% rowwise() %>% summarize(f(ans_0_1_2017_Genie, ans_0_1_2017_No_Genie, ans_1_0_2017_Genie, ans_1_0_2017_No_Genie))
dat4 <- bind_cols(dat2,dat3)

write_tsv(dat4,"nohonors_genie_vs_non_genie.tsv")


############################################################################################
pal_okabe_ito <- c("#E69F00", "#56B4E9", "#009E73", "#F0E442", "#0072B2", "#D55E00", "#CC79A7", "#000000")

dat <- read_tsv("pre_and_post_nohonors.tsv")
dat.A <- dat %>% mutate(result = str_glue("ans_{Pre}_{Post}")) %>%
  count(year, student, result) %>%
  pivot_wider(names_from="result",values_from="n", values_fill=list(n=0L)) %>%
  mutate(pre_acc = (ans_1_1+ans_1_0),
         post_acc = (ans_1_1+ans_0_1))
dat.B <- dat.A %>% mutate(diff = post_acc - pre_acc, diffs = sign(diff), excess = "no") %>%
  arrange(diff)

dat4 <- dat.B %>% group_by(year) %>% group_modify(function(tbl,y){
  j <- tail(1:nrow(tbl),sum(tbl$diffs))
  tbl$excess[j] <- "yes"
  tbl
})

pretty_year <- c(
  `2016_Genie` = "Genie 2016",
  `2017_Genie` = "Genie 2017",
  `2017_No_Genie` = "Non-Genie 2017"
)

ann_tbl <- tibble(pre_acc=10,post_acc=2.5,year="2017_No_Genie",excess="yes")

pdf(file="nohonors_F3.pdf")

set.seed(0x90210)
gg <- ggplot(dat4, aes(x=pre_acc,y=post_acc,color=excess)) + geom_jitter(width=0.3,height=0.3)
gg <- gg + geom_abline(slope=1,intercept=0)
gg <- gg + labs(x = "Pre-Score", y = "Post-Score")
gg <- gg + theme_minimal() + coord_fixed(xlim=c(0,22),ylim=c(0,22))
gg <- gg + facet_wrap(~year, labeller = labeller(year = pretty_year))
gg <- gg + scale_color_manual(values=pal_okabe_ito[c(8,2)])
gg <- gg + theme(legend.position = "none")
gg <- gg + geom_label(data=ann_tbl,label="        Excess\n        Improvement",hjust="left", nudge_x=-2,color="black",size=3) +
  geom_point(data=ann_tbl,size=5)

print(gg)
dev.off()

############################################################################################
#Visualization per question
#Transforms the pre-recitation scores columns on each data frame into a numeric vector
score_pre1_2017_G_without_honors <- as.numeric(as.character(df_2017_G_by_year_without_honors$Pre_1))
score_pre2_2017_G_without_honors <- as.numeric(as.character(df_2017_G_by_year_without_honors$Pre_2))
score_pre3_2017_G_without_honors <- as.numeric(as.character(df_2017_G_by_year_without_honors$Pre_3))
score_pre4_2017_G_without_honors <- as.numeric(as.character(df_2017_G_by_year_without_honors$Pre_4))
score_pre5_2017_G_without_honors <- as.numeric(as.character(df_2017_G_by_year_without_honors$Pre_5))
score_pre6_2017_G_without_honors <- as.numeric(as.character(df_2017_G_by_year_without_honors$Pre_6))
score_pre7_2017_G_without_honors <- as.numeric(as.character(df_2017_G_by_year_without_honors$Pre_7))
score_pre8_2017_G_without_honors <- as.numeric(as.character(df_2017_G_by_year_without_honors$Pre_8))
score_pre9_2017_G_without_honors <- as.numeric(as.character(df_2017_G_by_year_without_honors$Pre_9))
score_pre10_2017_G_without_honors <- as.numeric(as.character(df_2017_G_by_year_without_honors$Pre_10))
score_pre11_2017_G_without_honors <- as.numeric(as.character(df_2017_G_by_year_without_honors$Pre_11))
score_pre12_2017_G_without_honors <- as.numeric(as.character(df_2017_G_by_year_without_honors$Pre_12))
score_pre13_2017_G_without_honors <- as.numeric(as.character(df_2017_G_by_year_without_honors$Pre_13))
score_pre14_2017_G_without_honors <- as.numeric(as.character(df_2017_G_by_year_without_honors$Pre_14))
score_pre15_2017_G_without_honors <- as.numeric(as.character(df_2017_G_by_year_without_honors$Pre_15))
score_pre16_2017_G_without_honors <- as.numeric(as.character(df_2017_G_by_year_without_honors$Pre_16))
score_pre17_2017_G_without_honors <- as.numeric(as.character(df_2017_G_by_year_without_honors$Pre_17))
score_pre18_2017_G_without_honors <- as.numeric(as.character(df_2017_G_by_year_without_honors$Pre_18))
score_pre19_2017_G_without_honors <- as.numeric(as.character(df_2017_G_by_year_without_honors$Pre_19))
score_pre20_2017_G_without_honors <- as.numeric(as.character(df_2017_G_by_year_without_honors$Pre_20))
score_pre21_2017_G_without_honors <- as.numeric(as.character(df_2017_G_by_year_without_honors$Pre_21))
score_pre22_2017_G_without_honors <- as.numeric(as.character(df_2017_G_by_year_without_honors$Pre_22))

score_pre1_2017_NG_without_honors <- as.numeric(as.character(df_2017_NG_by_year_without_honors$Pre_1))
score_pre2_2017_NG_without_honors <- as.numeric(as.character(df_2017_NG_by_year_without_honors$Pre_2))
score_pre3_2017_NG_without_honors <- as.numeric(as.character(df_2017_NG_by_year_without_honors$Pre_3))
score_pre4_2017_NG_without_honors <- as.numeric(as.character(df_2017_NG_by_year_without_honors$Pre_4))
score_pre5_2017_NG_without_honors <- as.numeric(as.character(df_2017_NG_by_year_without_honors$Pre_5))
score_pre6_2017_NG_without_honors <- as.numeric(as.character(df_2017_NG_by_year_without_honors$Pre_6))
score_pre7_2017_NG_without_honors <- as.numeric(as.character(df_2017_NG_by_year_without_honors$Pre_7))
score_pre8_2017_NG_without_honors <- as.numeric(as.character(df_2017_NG_by_year_without_honors$Pre_8))
score_pre9_2017_NG_without_honors <- as.numeric(as.character(df_2017_NG_by_year_without_honors$Pre_9))
score_pre10_2017_NG_without_honors <- as.numeric(as.character(df_2017_NG_by_year_without_honors$Pre_10))
score_pre11_2017_NG_without_honors <- as.numeric(as.character(df_2017_NG_by_year_without_honors$Pre_11))
score_pre12_2017_NG_without_honors <- as.numeric(as.character(df_2017_NG_by_year_without_honors$Pre_12))
score_pre13_2017_NG_without_honors <- as.numeric(as.character(df_2017_NG_by_year_without_honors$Pre_13))
score_pre14_2017_NG_without_honors <- as.numeric(as.character(df_2017_NG_by_year_without_honors$Pre_14))
score_pre15_2017_NG_without_honors <- as.numeric(as.character(df_2017_NG_by_year_without_honors$Pre_15))
score_pre16_2017_NG_without_honors <- as.numeric(as.character(df_2017_NG_by_year_without_honors$Pre_16))
score_pre17_2017_NG_without_honors <- as.numeric(as.character(df_2017_NG_by_year_without_honors$Pre_17))
score_pre18_2017_NG_without_honors <- as.numeric(as.character(df_2017_NG_by_year_without_honors$Pre_18))
score_pre19_2017_NG_without_honors <- as.numeric(as.character(df_2017_NG_by_year_without_honors$Pre_19))
score_pre20_2017_NG_without_honors <- as.numeric(as.character(df_2017_NG_by_year_without_honors$Pre_20))
score_pre21_2017_NG_without_honors <- as.numeric(as.character(df_2017_NG_by_year_without_honors$Pre_21))
score_pre22_2017_NG_without_honors <- as.numeric(as.character(df_2017_NG_by_year_without_honors$Pre_22))

#Gets score for that question
score_pre1_2017_G_val_without_honors <- sum(score_pre1_2017_G_without_honors)/length(score_pre1_2017_G_without_honors)
score_pre2_2017_G_val_without_honors <- sum(score_pre2_2017_G_without_honors)/length(score_pre2_2017_G_without_honors)
score_pre3_2017_G_val_without_honors <- sum(score_pre3_2017_G_without_honors)/length(score_pre3_2017_G_without_honors)
score_pre4_2017_G_val_without_honors <- sum(score_pre4_2017_G_without_honors)/length(score_pre4_2017_G_without_honors)
score_pre5_2017_G_val_without_honors <- sum(score_pre5_2017_G_without_honors)/length(score_pre5_2017_G_without_honors)
score_pre6_2017_G_val_without_honors <- sum(score_pre6_2017_G_without_honors)/length(score_pre6_2017_G_without_honors)
score_pre7_2017_G_val_without_honors <- sum(score_pre7_2017_G_without_honors)/length(score_pre7_2017_G_without_honors)
score_pre8_2017_G_val_without_honors <- sum(score_pre8_2017_G_without_honors)/length(score_pre8_2017_G_without_honors)
score_pre9_2017_G_val_without_honors <- sum(score_pre9_2017_G_without_honors)/length(score_pre9_2017_G_without_honors)
score_pre10_2017_G_val_without_honors <- sum(score_pre10_2017_G_without_honors)/length(score_pre10_2017_G_without_honors)
score_pre11_2017_G_val_without_honors <- sum(score_pre11_2017_G_without_honors)/length(score_pre11_2017_G_without_honors)
score_pre12_2017_G_val_without_honors <- sum(score_pre12_2017_G_without_honors)/length(score_pre12_2017_G_without_honors)
score_pre13_2017_G_val_without_honors <- sum(score_pre13_2017_G_without_honors)/length(score_pre13_2017_G_without_honors)
score_pre14_2017_G_val_without_honors <- sum(score_pre14_2017_G_without_honors)/length(score_pre14_2017_G_without_honors)
score_pre15_2017_G_val_without_honors <- sum(score_pre15_2017_G_without_honors)/length(score_pre15_2017_G_without_honors)
score_pre16_2017_G_val_without_honors <- sum(score_pre16_2017_G_without_honors)/length(score_pre16_2017_G_without_honors)
score_pre17_2017_G_val_without_honors <- sum(score_pre17_2017_G_without_honors)/length(score_pre17_2017_G_without_honors)
score_pre18_2017_G_val_without_honors <- sum(score_pre18_2017_G_without_honors)/length(score_pre18_2017_G_without_honors)
score_pre19_2017_G_val_without_honors <- sum(score_pre19_2017_G_without_honors)/length(score_pre19_2017_G_without_honors)
score_pre20_2017_G_val_without_honors <- sum(score_pre20_2017_G_without_honors)/length(score_pre20_2017_G_without_honors)
score_pre21_2017_G_val_without_honors <- sum(score_pre21_2017_G_without_honors)/length(score_pre21_2017_G_without_honors)
score_pre22_2017_G_val_without_honors <- sum(score_pre22_2017_G_without_honors)/length(score_pre22_2017_G_without_honors)

score_pre1_2017_NG_val_without_honors <- sum(score_pre1_2017_NG_without_honors)/length(score_pre1_2017_NG_without_honors)
score_pre2_2017_NG_val_without_honors <- sum(score_pre2_2017_NG_without_honors)/length(score_pre2_2017_NG_without_honors)
score_pre3_2017_NG_val_without_honors <- sum(score_pre3_2017_NG_without_honors)/length(score_pre3_2017_NG_without_honors)
score_pre4_2017_NG_val_without_honors <- sum(score_pre4_2017_NG_without_honors)/length(score_pre4_2017_NG_without_honors)
score_pre5_2017_NG_val_without_honors <- sum(score_pre5_2017_NG_without_honors)/length(score_pre5_2017_NG_without_honors)
score_pre6_2017_NG_val_without_honors <- sum(score_pre6_2017_NG_without_honors)/length(score_pre6_2017_NG_without_honors)
score_pre7_2017_NG_val_without_honors <- sum(score_pre7_2017_NG_without_honors)/length(score_pre7_2017_NG_without_honors)
score_pre8_2017_NG_val_without_honors <- sum(score_pre8_2017_NG_without_honors)/length(score_pre8_2017_NG_without_honors)
score_pre9_2017_NG_val_without_honors <- sum(score_pre9_2017_NG_without_honors)/length(score_pre9_2017_NG_without_honors)
score_pre10_2017_NG_val_without_honors <- sum(score_pre10_2017_NG_without_honors)/length(score_pre10_2017_NG_without_honors)
score_pre11_2017_NG_val_without_honors <- sum(score_pre11_2017_NG_without_honors)/length(score_pre11_2017_NG_without_honors)
score_pre12_2017_NG_val_without_honors <- sum(score_pre12_2017_NG_without_honors)/length(score_pre12_2017_NG_without_honors)
score_pre13_2017_NG_val_without_honors <- sum(score_pre13_2017_NG_without_honors)/length(score_pre13_2017_NG_without_honors)
score_pre14_2017_NG_val_without_honors <- sum(score_pre14_2017_NG_without_honors)/length(score_pre14_2017_NG_without_honors)
score_pre15_2017_NG_val_without_honors <- sum(score_pre15_2017_NG_without_honors)/length(score_pre15_2017_NG_without_honors)
score_pre16_2017_NG_val_without_honors <- sum(score_pre16_2017_NG_without_honors)/length(score_pre16_2017_NG_without_honors)
score_pre17_2017_NG_val_without_honors <- sum(score_pre17_2017_NG_without_honors)/length(score_pre17_2017_NG_without_honors)
score_pre18_2017_NG_val_without_honors <- sum(score_pre18_2017_NG_without_honors)/length(score_pre18_2017_NG_without_honors)
score_pre19_2017_NG_val_without_honors <- sum(score_pre19_2017_NG_without_honors)/length(score_pre19_2017_NG_without_honors)
score_pre20_2017_NG_val_without_honors <- sum(score_pre20_2017_NG_without_honors)/length(score_pre20_2017_NG_without_honors)
score_pre21_2017_NG_val_without_honors <- sum(score_pre21_2017_NG_without_honors)/length(score_pre21_2017_NG_without_honors)
score_pre22_2017_NG_val_without_honors <- sum(score_pre22_2017_NG_without_honors)/length(score_pre22_2017_NG_without_honors)


#Transforms the post-recitation scores columns on each data frame into a numeric vector
score_post1_2017_G_without_honors <- as.numeric(as.character(df_2017_G_by_year_without_honors$Post_1))
score_post2_2017_G_without_honors <- as.numeric(as.character(df_2017_G_by_year_without_honors$Post_2))
score_post3_2017_G_without_honors <- as.numeric(as.character(df_2017_G_by_year_without_honors$Post_3))
score_post4_2017_G_without_honors <- as.numeric(as.character(df_2017_G_by_year_without_honors$Post_4))
score_post5_2017_G_without_honors <- as.numeric(as.character(df_2017_G_by_year_without_honors$Post_5))
score_post6_2017_G_without_honors <- as.numeric(as.character(df_2017_G_by_year_without_honors$Post_6))
score_post7_2017_G_without_honors <- as.numeric(as.character(df_2017_G_by_year_without_honors$Post_7))
score_post8_2017_G_without_honors <- as.numeric(as.character(df_2017_G_by_year_without_honors$Post_8))
score_post9_2017_G_without_honors <- as.numeric(as.character(df_2017_G_by_year_without_honors$Post_9))
score_post10_2017_G_without_honors <- as.numeric(as.character(df_2017_G_by_year_without_honors$Post_10))
score_post11_2017_G_without_honors <- as.numeric(as.character(df_2017_G_by_year_without_honors$Post_11))
score_post12_2017_G_without_honors <- as.numeric(as.character(df_2017_G_by_year_without_honors$Post_12))
score_post13_2017_G_without_honors <- as.numeric(as.character(df_2017_G_by_year_without_honors$Post_13))
score_post14_2017_G_without_honors <- as.numeric(as.character(df_2017_G_by_year_without_honors$Post_14))
score_post15_2017_G_without_honors <- as.numeric(as.character(df_2017_G_by_year_without_honors$Post_15))
score_post16_2017_G_without_honors <- as.numeric(as.character(df_2017_G_by_year_without_honors$Post_16))
score_post17_2017_G_without_honors <- as.numeric(as.character(df_2017_G_by_year_without_honors$Post_17))
score_post18_2017_G_without_honors <- as.numeric(as.character(df_2017_G_by_year_without_honors$Post_18))
score_post19_2017_G_without_honors <- as.numeric(as.character(df_2017_G_by_year_without_honors$Post_19))
score_post20_2017_G_without_honors <- as.numeric(as.character(df_2017_G_by_year_without_honors$Post_20))
score_post21_2017_G_without_honors <- as.numeric(as.character(df_2017_G_by_year_without_honors$Post_21))
score_post22_2017_G_without_honors <- as.numeric(as.character(df_2017_G_by_year_without_honors$Post_22))

score_post1_2017_NG_without_honors <- as.numeric(as.character(df_2017_NG_by_year_without_honors$Post_1))
score_post2_2017_NG_without_honors <- as.numeric(as.character(df_2017_NG_by_year_without_honors$Post_2))
score_post3_2017_NG_without_honors <- as.numeric(as.character(df_2017_NG_by_year_without_honors$Post_3))
score_post4_2017_NG_without_honors <- as.numeric(as.character(df_2017_NG_by_year_without_honors$Post_4))
score_post5_2017_NG_without_honors <- as.numeric(as.character(df_2017_NG_by_year_without_honors$Post_5))
score_post6_2017_NG_without_honors <- as.numeric(as.character(df_2017_NG_by_year_without_honors$Post_6))
score_post7_2017_NG_without_honors <- as.numeric(as.character(df_2017_NG_by_year_without_honors$Post_7))
score_post8_2017_NG_without_honors <- as.numeric(as.character(df_2017_NG_by_year_without_honors$Post_8))
score_post9_2017_NG_without_honors <- as.numeric(as.character(df_2017_NG_by_year_without_honors$Post_9))
score_post10_2017_NG_without_honors <- as.numeric(as.character(df_2017_NG_by_year_without_honors$Post_10))
score_post11_2017_NG_without_honors <- as.numeric(as.character(df_2017_NG_by_year_without_honors$Post_11))
score_post12_2017_NG_without_honors <- as.numeric(as.character(df_2017_NG_by_year_without_honors$Post_12))
score_post13_2017_NG_without_honors <- as.numeric(as.character(df_2017_NG_by_year_without_honors$Post_13))
score_post14_2017_NG_without_honors <- as.numeric(as.character(df_2017_NG_by_year_without_honors$Post_14))
score_post15_2017_NG_without_honors <- as.numeric(as.character(df_2017_NG_by_year_without_honors$Post_15))
score_post16_2017_NG_without_honors <- as.numeric(as.character(df_2017_NG_by_year_without_honors$Post_16))
score_post17_2017_NG_without_honors <- as.numeric(as.character(df_2017_NG_by_year_without_honors$Post_17))
score_post18_2017_NG_without_honors <- as.numeric(as.character(df_2017_NG_by_year_without_honors$Post_18))
score_post19_2017_NG_without_honors <- as.numeric(as.character(df_2017_NG_by_year_without_honors$Post_19))
score_post20_2017_NG_without_honors <- as.numeric(as.character(df_2017_NG_by_year_without_honors$Post_20))
score_post21_2017_NG_without_honors <- as.numeric(as.character(df_2017_NG_by_year_without_honors$Post_21))
score_post22_2017_NG_without_honors <- as.numeric(as.character(df_2017_NG_by_year_without_honors$Post_22))

#Gets score for that question
score_post1_2017_G_val_without_honors <- sum(score_post1_2017_G_without_honors)/length(score_post1_2017_G_without_honors)
score_post2_2017_G_val_without_honors <- sum(score_post2_2017_G_without_honors)/length(score_post2_2017_G_without_honors)
score_post3_2017_G_val_without_honors <- sum(score_post3_2017_G_without_honors)/length(score_post3_2017_G_without_honors)
score_post4_2017_G_val_without_honors <- sum(score_post4_2017_G_without_honors)/length(score_post4_2017_G_without_honors)
score_post5_2017_G_val_without_honors <- sum(score_post5_2017_G_without_honors)/length(score_post5_2017_G_without_honors)
score_post6_2017_G_val_without_honors <- sum(score_post6_2017_G_without_honors)/length(score_post6_2017_G_without_honors)
score_post7_2017_G_val_without_honors <- sum(score_post7_2017_G_without_honors)/length(score_post7_2017_G_without_honors)
score_post8_2017_G_val_without_honors <- sum(score_post8_2017_G_without_honors)/length(score_post8_2017_G_without_honors)
score_post9_2017_G_val_without_honors <- sum(score_post9_2017_G_without_honors)/length(score_post9_2017_G_without_honors)
score_post10_2017_G_val_without_honors <- sum(score_post10_2017_G_without_honors)/length(score_post10_2017_G_without_honors)
score_post11_2017_G_val_without_honors <- sum(score_post11_2017_G_without_honors)/length(score_post11_2017_G_without_honors)
score_post12_2017_G_val_without_honors <- sum(score_post12_2017_G_without_honors)/length(score_post12_2017_G_without_honors)
score_post13_2017_G_val_without_honors <- sum(score_post13_2017_G_without_honors)/length(score_post13_2017_G_without_honors)
score_post14_2017_G_val_without_honors <- sum(score_post14_2017_G_without_honors)/length(score_post14_2017_G_without_honors)
score_post15_2017_G_val_without_honors <- sum(score_post15_2017_G_without_honors)/length(score_post15_2017_G_without_honors)
score_post16_2017_G_val_without_honors <- sum(score_post16_2017_G_without_honors)/length(score_post16_2017_G_without_honors)
score_post17_2017_G_val_without_honors <- sum(score_post17_2017_G_without_honors)/length(score_post17_2017_G_without_honors)
score_post18_2017_G_val_without_honors <- sum(score_post18_2017_G_without_honors)/length(score_post18_2017_G_without_honors)
score_post19_2017_G_val_without_honors <- sum(score_post19_2017_G_without_honors)/length(score_post19_2017_G_without_honors)
score_post20_2017_G_val_without_honors <- sum(score_post20_2017_G_without_honors)/length(score_post20_2017_G_without_honors)
score_post21_2017_G_val_without_honors <- sum(score_post21_2017_G_without_honors)/length(score_post21_2017_G_without_honors)
score_post22_2017_G_val_without_honors <- sum(score_post22_2017_G_without_honors)/length(score_post22_2017_G_without_honors)

score_post1_2017_NG_val_without_honors <- sum(score_post1_2017_NG_without_honors)/length(score_post1_2017_NG_without_honors)
score_post2_2017_NG_val_without_honors <- sum(score_post2_2017_NG_without_honors)/length(score_post2_2017_NG_without_honors)
score_post3_2017_NG_val_without_honors <- sum(score_post3_2017_NG_without_honors)/length(score_post3_2017_NG_without_honors)
score_post4_2017_NG_val_without_honors <- sum(score_post4_2017_NG_without_honors)/length(score_post4_2017_NG_without_honors)
score_post5_2017_NG_val_without_honors <- sum(score_post5_2017_NG_without_honors)/length(score_post5_2017_NG_without_honors)
score_post6_2017_NG_val_without_honors <- sum(score_post6_2017_NG_without_honors)/length(score_post6_2017_NG_without_honors)
score_post7_2017_NG_val_without_honors <- sum(score_post7_2017_NG_without_honors)/length(score_post7_2017_NG_without_honors)
score_post8_2017_NG_val_without_honors <- sum(score_post8_2017_NG_without_honors)/length(score_post8_2017_NG_without_honors)
score_post9_2017_NG_val_without_honors <- sum(score_post9_2017_NG_without_honors)/length(score_post9_2017_NG_without_honors)
score_post10_2017_NG_val_without_honors <- sum(score_post10_2017_NG_without_honors)/length(score_post10_2017_NG_without_honors)
score_post11_2017_NG_val_without_honors <- sum(score_post11_2017_NG_without_honors)/length(score_post11_2017_NG_without_honors)
score_post12_2017_NG_val_without_honors <- sum(score_post12_2017_NG_without_honors)/length(score_post12_2017_NG_without_honors)
score_post13_2017_NG_val_without_honors <- sum(score_post13_2017_NG_without_honors)/length(score_post13_2017_NG_without_honors)
score_post14_2017_NG_val_without_honors <- sum(score_post14_2017_NG_without_honors)/length(score_post14_2017_NG_without_honors)
score_post15_2017_NG_val_without_honors <- sum(score_post15_2017_NG_without_honors)/length(score_post15_2017_NG_without_honors)
score_post16_2017_NG_val_without_honors <- sum(score_post16_2017_NG_without_honors)/length(score_post16_2017_NG_without_honors)
score_post17_2017_NG_val_without_honors <- sum(score_post17_2017_NG_without_honors)/length(score_post17_2017_NG_without_honors)
score_post18_2017_NG_val_without_honors <- sum(score_post18_2017_NG_without_honors)/length(score_post18_2017_NG_without_honors)
score_post19_2017_NG_val_without_honors <- sum(score_post19_2017_NG_without_honors)/length(score_post19_2017_NG_without_honors)
score_post20_2017_NG_val_without_honors <- sum(score_post20_2017_NG_without_honors)/length(score_post20_2017_NG_without_honors)
score_post21_2017_NG_val_without_honors <- sum(score_post21_2017_NG_without_honors)/length(score_post21_2017_NG_without_honors)
score_post22_2017_NG_val_without_honors <- sum(score_post22_2017_NG_without_honors)/length(score_post22_2017_NG_without_honors)


#Preparing vectors for the dataframe
#Order based on Figure2a -Key concepts| Table 3 - Misconceptions
questionA2T3 <- c("Q1","Q3","Q15","Q4","Q10","Q13","Q16","Q7","Q5","Q6","Q8","Q2","Q9","Q12","Q17","Q20","Q14","Q19","Q22","Q11","Q18","Q21")
time_pre <- rep("Pre", 22)
time_post <- rep("Post", 22)

#Vector of pre-recitation and post-recitation scores by question on 2017 genie classes
pre_recitation_scores_2017_G_without_honorsA2T3 <- c(score_pre1_2017_G_val_without_honors,
                                  score_pre3_2017_G_val_without_honors,
                                  score_pre15_2017_G_val_without_honors,
                                  score_pre4_2017_G_val_without_honors,
                                  score_pre10_2017_G_val_without_honors,
                                  score_pre13_2017_G_val_without_honors,
                                  score_pre16_2017_G_val_without_honors,
                                  score_pre7_2017_G_val_without_honors,
                                  score_pre5_2017_G_val_without_honors,
                                  score_pre6_2017_G_val_without_honors,
                                  score_pre8_2017_G_val_without_honors,
                                  score_pre2_2017_G_val_without_honors,
                                  score_pre9_2017_G_val_without_honors,
                                  score_pre12_2017_G_val_without_honors,
                                  score_pre17_2017_G_val_without_honors,
                                  score_pre20_2017_G_val_without_honors,
                                  score_pre14_2017_G_val_without_honors,
                                  score_pre19_2017_G_val_without_honors,
                                  score_pre22_2017_G_val_without_honors,
                                  score_pre11_2017_G_val_without_honors,
                                  score_pre18_2017_G_val_without_honors,
                                  score_pre21_2017_G_val_without_honors)


post_recitation_scores_2017_G_without_honorsA2T3 <- c(score_post1_2017_G_val_without_honors,
                                   score_post3_2017_G_val_without_honors,
                                   score_post15_2017_G_val_without_honors,
                                   score_post4_2017_G_val_without_honors,
                                   score_post10_2017_G_val_without_honors,
                                   score_post13_2017_G_val_without_honors,
                                   score_post16_2017_G_val_without_honors,
                                   score_post7_2017_G_val_without_honors,
                                   score_post5_2017_G_val_without_honors,
                                   score_post6_2017_G_val_without_honors,
                                   score_post8_2017_G_val_without_honors,
                                   score_post2_2017_G_val_without_honors,
                                   score_post9_2017_G_val_without_honors,
                                   score_post12_2017_G_val_without_honors,
                                   score_post17_2017_G_val_without_honors,
                                   score_post20_2017_G_val_without_honors,
                                   score_post14_2017_G_val_without_honors,
                                   score_post19_2017_G_val_without_honors,
                                   score_post22_2017_G_val_without_honors,
                                   score_post11_2017_G_val_without_honors,
                                   score_post18_2017_G_val_without_honors,
                                   score_post21_2017_G_val_without_honors)

#Pre, post-, and combined recitations dataframes for plot 2017 genie class
df_scores_by_question_pre_2017_G_without_honorsA2T3 <- data.frame(scores_2017_G_without_honors = pre_recitation_scores_2017_G_without_honorsA2T3,
                                               time_of_recitation = time_pre,
                                               question=questionA2T3)

df_scores_by_question_post_2017_G_without_honorsA2T3 <- data.frame(scores_2017_G_without_honors = post_recitation_scores_2017_G_without_honorsA2T3,
                                                time_of_recitation = time_post,
                                                question=questionA2T3)

df_scores_by_question_combined_2017_G_without_honorsA2T3 <- rbind(df_scores_by_question_pre_2017_G_without_honorsA2T3, df_scores_by_question_post_2017_G_without_honorsA2T3)


#Vector of pre-recitation and post-recitation scores by question on 2017 no genie classes
pre_recitation_scores_2017_NGA2T3 <- c(score_pre1_2017_NG_val_without_honors,
                                   score_pre3_2017_NG_val_without_honors,
                                   score_pre15_2017_NG_val_without_honors,
                                   score_pre4_2017_NG_val_without_honors,
                                   score_pre10_2017_NG_val_without_honors,
                                   score_pre13_2017_NG_val_without_honors,
                                   score_pre16_2017_NG_val_without_honors,
                                   score_pre7_2017_NG_val_without_honors,
                                   score_pre5_2017_NG_val_without_honors,
                                   score_pre6_2017_NG_val_without_honors,
                                   score_pre8_2017_NG_val_without_honors,
                                   score_pre2_2017_NG_val_without_honors,
                                   score_pre9_2017_NG_val_without_honors,
                                   score_pre12_2017_NG_val_without_honors,
                                   score_pre17_2017_NG_val_without_honors,
                                   score_pre20_2017_NG_val_without_honors,
                                   score_pre14_2017_NG_val_without_honors,
                                   score_pre19_2017_NG_val_without_honors,
                                   score_pre22_2017_NG_val_without_honors,
                                   score_pre11_2017_NG_val_without_honors,
                                   score_pre18_2017_NG_val_without_honors,
                                   score_pre21_2017_NG_val_without_honors)

post_recitation_scores_2017_NGA2T3 <- c(score_post1_2017_NG_val_without_honors,
                                    score_post3_2017_NG_val_without_honors,
                                    score_post15_2017_NG_val_without_honors,
                                    score_post4_2017_NG_val_without_honors,
                                    score_post10_2017_NG_val_without_honors,
                                    score_post13_2017_NG_val_without_honors,
                                    score_post16_2017_NG_val_without_honors,
                                    score_post7_2017_NG_val_without_honors,
                                    score_post5_2017_NG_val_without_honors,
                                    score_post6_2017_NG_val_without_honors,
                                    score_post8_2017_NG_val_without_honors,
                                    score_post2_2017_NG_val_without_honors,
                                    score_post9_2017_NG_val_without_honors,
                                    score_post12_2017_NG_val_without_honors,
                                    score_post17_2017_NG_val_without_honors,
                                    score_post20_2017_NG_val_without_honors,
                                    score_post14_2017_NG_val_without_honors,
                                    score_post19_2017_NG_val_without_honors,
                                    score_post22_2017_NG_val_without_honors,
                                    score_post11_2017_NG_val_without_honors,
                                    score_post18_2017_NG_val_without_honors,
                                    score_post21_2017_NG_val_without_honors)


#Pre, post-, and combined recitations dataframes for plot 2017 no genie class
df_scores_by_question_pre_2017_NGA2T3 <- data.frame(scores_2017_NG = pre_recitation_scores_2017_NGA2T3,
                                                time_of_recitation = time_pre,
                                                question=questionA2T3)

df_scores_by_question_post_2017_NGA2T3 <- data.frame(scores_2017_NG = post_recitation_scores_2017_NGA2T3,
                                                 time_of_recitation = time_post,
                                                 question=questionA2T3)

df_scores_by_question_combined_2017_NGA2T3 <- rbind(df_scores_by_question_pre_2017_NGA2T3, df_scores_by_question_post_2017_NGA2T3)


#Diff without honors
score_diff1_2017_G_without_honors <- score_post1_2017_G_val_without_honors-score_pre1_2017_G_val_without_honors
score_diff2_2017_G_without_honors <- score_post2_2017_G_val_without_honors-score_pre2_2017_G_val_without_honors
score_diff3_2017_G_without_honors <- score_post3_2017_G_val_without_honors-score_pre3_2017_G_val_without_honors
score_diff4_2017_G_without_honors <- score_post4_2017_G_val_without_honors-score_pre4_2017_G_val_without_honors
score_diff5_2017_G_without_honors <- score_post5_2017_G_val_without_honors-score_pre5_2017_G_val_without_honors
score_diff6_2017_G_without_honors <- score_post6_2017_G_val_without_honors-score_pre6_2017_G_val_without_honors
score_diff7_2017_G_without_honors <- score_post7_2017_G_val_without_honors-score_pre7_2017_G_val_without_honors
score_diff8_2017_G_without_honors <- score_post8_2017_G_val_without_honors-score_pre8_2017_G_val_without_honors
score_diff9_2017_G_without_honors <- score_post9_2017_G_val_without_honors-score_pre9_2017_G_val_without_honors
score_diff10_2017_G_without_honors <- score_post10_2017_G_val_without_honors-score_pre10_2017_G_val_without_honors
score_diff11_2017_G_without_honors <- score_post11_2017_G_val_without_honors-score_pre11_2017_G_val_without_honors
score_diff12_2017_G_without_honors <- score_post12_2017_G_val_without_honors-score_pre12_2017_G_val_without_honors
score_diff13_2017_G_without_honors <- score_post13_2017_G_val_without_honors-score_pre13_2017_G_val_without_honors
score_diff14_2017_G_without_honors <- score_post14_2017_G_val_without_honors-score_pre14_2017_G_val_without_honors
score_diff15_2017_G_without_honors <- score_post15_2017_G_val_without_honors-score_pre15_2017_G_val_without_honors
score_diff16_2017_G_without_honors <- score_post16_2017_G_val_without_honors-score_pre16_2017_G_val_without_honors
score_diff17_2017_G_without_honors <- score_post17_2017_G_val_without_honors-score_pre17_2017_G_val_without_honors
score_diff18_2017_G_without_honors <- score_post18_2017_G_val_without_honors-score_pre18_2017_G_val_without_honors
score_diff19_2017_G_without_honors <- score_post19_2017_G_val_without_honors-score_pre19_2017_G_val_without_honors
score_diff20_2017_G_without_honors <- score_post20_2017_G_val_without_honors-score_pre20_2017_G_val_without_honors
score_diff21_2017_G_without_honors <- score_post21_2017_G_val_without_honors-score_pre21_2017_G_val_without_honors
score_diff22_2017_G_without_honors <- score_post22_2017_G_val_without_honors-score_pre22_2017_G_val_without_honors


diff_recitation_scores_2017_G_without_honors2AT3 <- c(score_diff1_2017_G_without_honors,
                                   score_diff3_2017_G_without_honors,
                                   score_diff4_2017_G_without_honors,
                                   score_diff10_2017_G_without_honors,
                                   score_diff13_2017_G_without_honors,
                                   score_diff15_2017_G_without_honors,
                                   score_diff16_2017_G_without_honors,
                                   score_diff7_2017_G_without_honors,
                                   score_diff5_2017_G_without_honors,
                                   score_diff6_2017_G_without_honors,
                                   score_diff8_2017_G_without_honors,
                                   score_diff2_2017_G_without_honors,
                                   score_diff9_2017_G_without_honors,
                                   score_diff12_2017_G_without_honors,
                                   score_diff17_2017_G_without_honors,
                                   score_diff20_2017_G_without_honors,
                                   score_diff14_2017_G_without_honors,
                                   score_diff19_2017_G_without_honors,
                                   score_diff22_2017_G_without_honors,
                                   score_diff11_2017_G_without_honors,
                                   score_diff18_2017_G_without_honors,
                                   score_diff21_2017_G_without_honors)


diff_recitation_scores_2017_G_without_honorsA2 <- c(score_diff1_2017_G_without_honors,
                                                  score_diff3_2017_G_without_honors,
                                                  score_diff4_2017_G_without_honors,
                                                  score_diff10_2017_G_without_honors,
                                                  score_diff13_2017_G_without_honors,
                                                  score_diff15_2017_G_without_honors,
                                                  score_diff16_2017_G_without_honors,
                                                  score_diff21_2017_G_without_honors,
                                                  score_diff22_2017_G_without_honors,
                                                  score_diff19_2017_G_without_honors,
                                                  score_diff7_2017_G_without_honors,
                                                  score_diff9_2017_G_without_honors,
                                                  score_diff20_2017_G_without_honors,
                                                  score_diff18_2017_G_without_honors,
                                                  score_diff11_2017_G_without_honors,
                                                  score_diff2_2017_G_without_honors,
                                                  score_diff14_2017_G_without_honors,
                                                  score_diff5_2017_G_without_honors,
                                                  score_diff17_2017_G_without_honors,
                                                  score_diff12_2017_G_without_honors,
                                                  score_diff6_2017_G_without_honors,
                                                  score_diff8_2017_G_without_honors)


###Order Fig2|Table3 in Price et al 2014
time_diffG <- rep("Diff-score Genie", 22)
time_diffNG <- rep("Diff-score no Genie", 22)

df_diff_scores_by_question_G_without_honors2AT3 <- data.frame(diff_recitation_scores = diff_recitation_scores_2017_G_without_honors2AT3,
                                                           question=question2AT3,
                                                           time=time_diffG)

df_diff_scores_by_question_NG2AT3 <- data.frame(diff_recitation_scores = diff_recitation_scores_2017_NG2AT3,
                                            question=question2AT3,
                                            time=time_diffNG)


df_diff_scores_by_question_G_without_honors2AT3$question <- factor(df_diff_scores_by_question_G_without_honors2AT3$question,
                                                               levels = df_diff_scores_by_question_G_without_honors2AT3$question)

df_diff_scores_by_question_NG2AT3$question <- factor(df_diff_scores_by_question_NG2AT3$question,
                                                  levels = df_diff_scores_by_question_NG2AT3$question)

df_diff_scores_by_question_combined2AT3 <- rbind(df_diff_scores_by_question_G_without_honors2AT3, df_diff_scores_by_question_NG2AT3)


pdf(file="nohonors_F5.pdf")

my_purple_colors <- c("orchid4","plum")
diff_per_question_plot2AT3 <- ggplot(data=df_diff_scores_by_question_combined2AT3, aes(x=question, y=diff_recitation_scores, fill=time)) +
  geom_bar(width=0.4, stat="identity", position=position_dodge(0.5)) +
  scale_fill_manual(values = my_purple_colors) +
  xlab("Question") +
  ylab("Scores") +
  labs(fill = "Recitation") +
  scale_x_discrete(limits= df_diff_scores_by_question_combined2AT3$p.question) + 
  theme(panel.grid.major = element_blank(), panel.grid.minor = element_blank(),
        panel.background = element_blank(), axis.line = element_line(colour = "black"))


print(diff_per_question_plot2AT3)
dev.off()

time_postG <- rep("Post-score Genie", 22)
time_postNG <- rep("Post-score no Genie", 22)

df_post_scores_by_question_G_without_honors2AT3 <- data.frame(post_recitation_scores = post_recitation_scores_2017_G_without_honorsA2T3,
                                                          question=question2AT3,
                                                          time=time_postG)

df_post_scores_by_question_NG2AT3 <- data.frame(post_recitation_scores = post_recitation_scores_2017_NG2AT3,
                                            question=question2AT3,
                                            time=time_postNG)


df_post_scores_by_question_G_without_honors2AT3$question <- factor(df_post_scores_by_question_G_without_honors2AT3$question,
                                                               levels = df_post_scores_by_question_G_without_honors2AT3$question)

df_post_scores_by_question_NG2AT3$question <- factor(df_post_scores_by_question_NG2AT3$question,
                                                 levels = df_post_scores_by_question_NG2AT3$question)

df_post_scores_by_question_combined2AT3 <- rbind(df_post_scores_by_question_G_without_honors2AT3, df_post_scores_by_question_NG2AT3)


pdf(file="nohonors_F4.pdf")

my_green_colors <- c("darkgreen","darkseagreen3")
post_per_question_plot2AT3 <- ggplot(data=df_post_scores_by_question_combined2AT3, aes(x=question, y=post_recitation_scores, fill=time)) +
  geom_bar(width=0.4, stat="identity", position=position_dodge(0.5)) +
  scale_fill_manual(values = my_green_colors) +
  xlab("Question") +
  ylab("Scores") +
  labs(fill = "Recitation") +
  scale_x_discrete(limits= df_post_scores_by_question_combined2AT3$p.question) +
  theme(panel.grid.major = element_blank(), panel.grid.minor = element_blank(),
      panel.background = element_blank(), axis.line = element_line(colour = "black"))


print(post_per_question_plot2AT3)
dev.off()
```
