## Supplementary figures and images for "Genie: An interactive real-time simulation for teaching genetic drift"

### SFile1.tiff

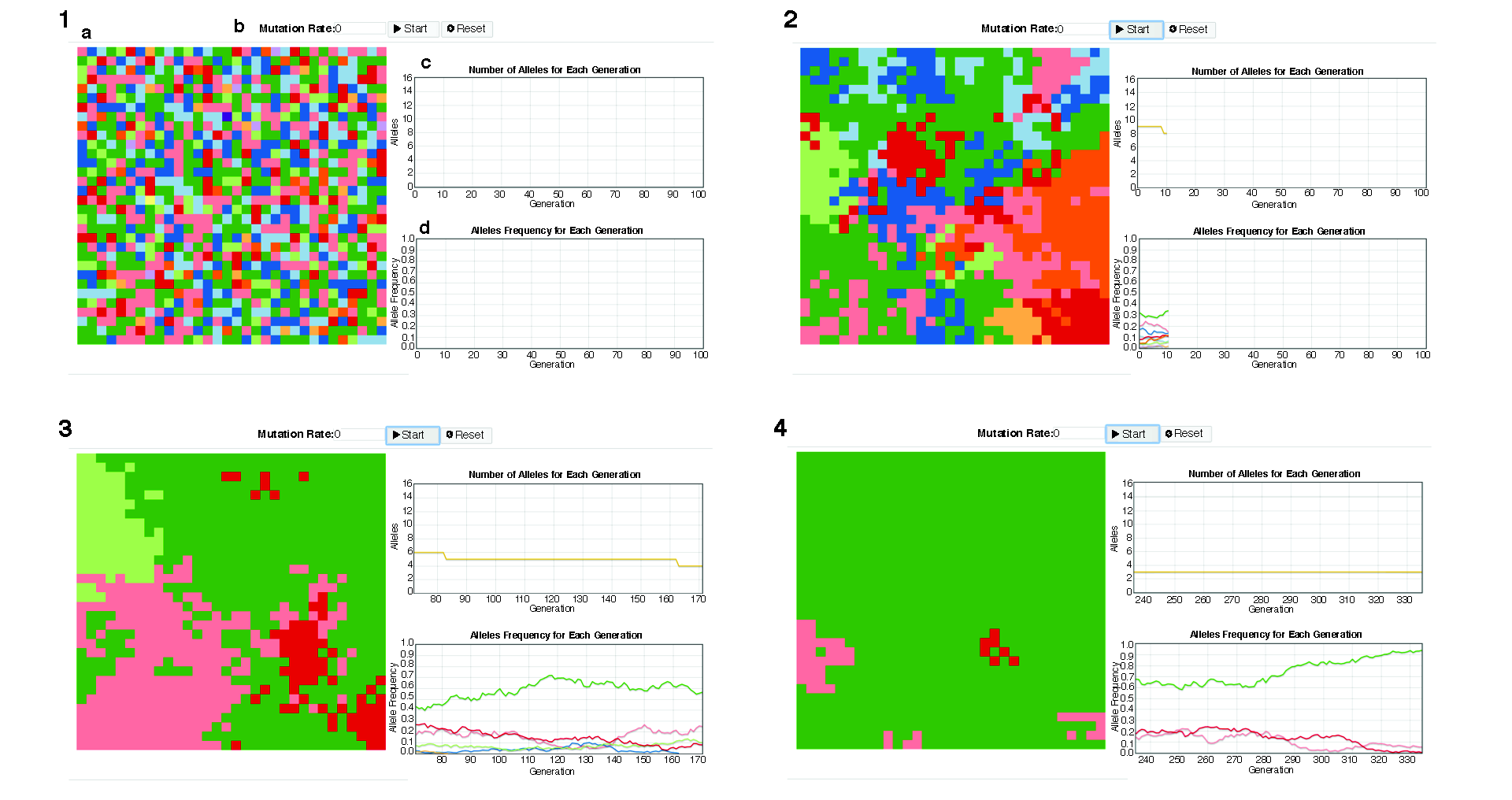

### SFile13.tif

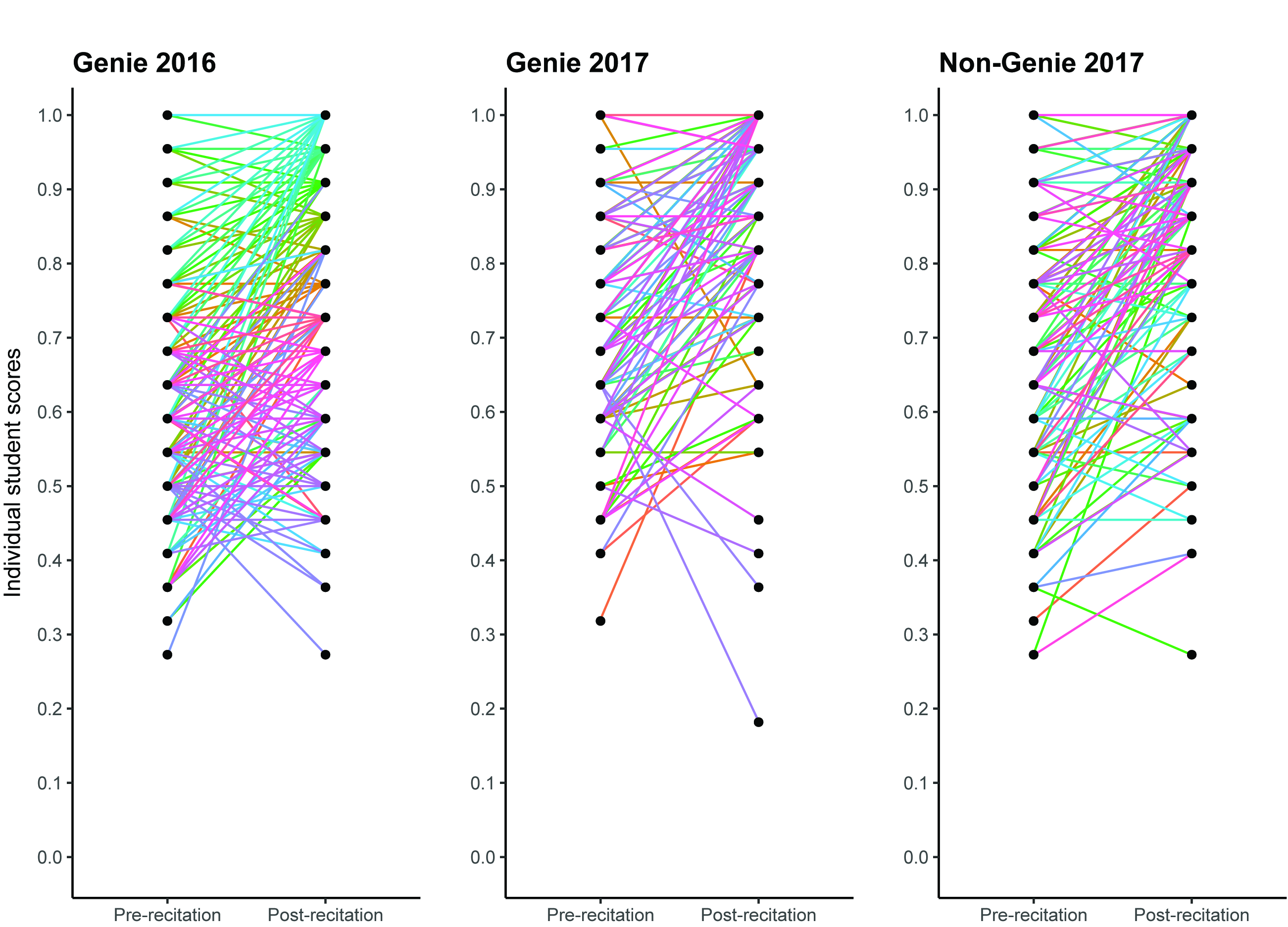
